# Pulse-Width Modulation of Gene Expression in Budding Yeast

**DOI:** 10.1101/2021.05.26.444658

**Authors:** Rainer Machné, Douglas B. Murray, Stephan H. Bernhart, Ilka M. Axmann, Peter F. Stadler

## Abstract

Metabolic oscillations are characterized by alternating phases of high and low respiratory activity, associated with transcription of genes involved in biosynthetic pathways and growth, and in catabolism and stress response. However, the functional consequences of transcriptome oscillations remain unclear, since most proteins are too stable to be affected by oscillatory transcript abundances. In this work, we investigate a transcriptome time series during an unstable state of the oscillation. Our analyses confirm previous suggestions that the relative times spent in the alternative transcription states are coupled to growth rate. This pulse-width modulation of transcription provides a simple mechanism for the long-standing question of how cells adjust their ribosome content and growth rate to environmental conditions. A mathematical model of this idea reproduces both the almost linear relation of transcript and protein abundances and the non-linear relation of oscillation periods to growth rate.

## Introduction

When yeast cultures are grown to a high cell density they tend to show collective metabolic dynamics, alternating between phases of high oxygen consumption (HOC) and low oxygen consumption (LOC). Numerous studies have shown that these oscillatory dynamics propagate throughout the metabolome, transcriptome and impinge on chromatin organization. The cycle period is dependent on growth conditions and the strain employed. Long period cycles (periods *τ_osc_* = 3h-8h) were explained by a partial synchronization of the cell division cycle (CDC). Glycogen stores are filled during LOC phase, which corresponds to the G1 phase of the CDC, and mobilized during HOC phase, which corresponds to the budding phase (***Küenzi and Fiechter, 1969***; ***von Meyenburg, 1969a***; ***Sonnleitner and Käppeli, 1986***; ***Münch et al., 1992***; ***Bellgardt, 1994***; ***Hjortso and Nielsen,1995***; ***Futcher, 2006***). ***Satroutdinov et al.*** (***1992***) then observed much shorter periods in the strain IFO 0233 (*τ_osc_* = 0.7h-1 h). IFO 0233, a distillery strain, questioned these prior models, as the glycogen storage cycle is reversed between the phases and ethanol is produced in LOC phase.

However, a similar temporal program is observed in both short period and long period experimental systems (***Machné and Murray, 2012***). Both show maxima of the cellular ATP/ADP ratio, followed by amino acid synthesis and a TORC1-mediated pulse of protein translation during the HOC phase (***von Meyenburg, 1969b***; ***Satroutdinov et al., 1992***; ***Hans et al., 2003***; ***Xu et al., 2004***; ***Müller, 2006***; ***Murray et al., 2007***; ***Machné and Murray, 2012***; ***Amariei et al., 2014***; ***O’Neill et al., 2020***). Global remodeling of promoter and gene body nucleosome organization occurs during the late LOC phase (***Amariei et al., 2014***; ***Nocetti and Whitehouse, 2016***). During the HOC phase, transcription progresses from a ribosome biogenesis cohort (`Ribi`) and cytoplasmic ribosomal protein genes (`RP`), to amino acid synthesis genes (`AA`) and mitochondrial ribosomal protein genes (`mtRP`) at the transition to the LOC phase. During the LOC phase, transcripts of a large group of stress-response and catabolic proteins (S/C) peak (***Klevecz et al., 2004***; ***Tu et al., 2005***; ***Slavov et al., 2011***; ***Machné and Murray, 2012***).

Several hypotheses on putative functions of the temporal transcription program have been suggested. The functional profiles of co-expressed cohorts match metabolic activity, and the initial hypothesis was a “just-in-time” model of gene expression (JIT), where enzymes are expressed when required within the metabolic cycle (***Klevecz et al., 2004***; ***Tu et al., 2005***; ***Murray et al., 2007***). However, protein half-lives in yeast are now thought to be much longer than initially reported (***Christiano et al., 2014***). This dampens the effect of periodic transcript on protein abundances (***Lück et al., 2014***). Indeed, recent proteomic studies found no (**preprint:** ***Feltham et al.*** (***2019***)) or only few (***O’Neill et al., 2020***) periodic protein abundances in long period systems. ***Slavov and Botstein (2011)*** and ***Burnetti et al.*** (***2016***) suggested an alternative hypothesis, based on the observation that the relative duration of the LOC phase varies strongly with growth rate while HOC phase duration only subtly changes. This would result in different absolute abundances of the proteins produced from HOC- and LOC-specific transcripts and could underlie growth rate-dependent cellular resource allocation (***Maaløe, 1979***; ***Molenaar et al., 2009***). Due to the analogy to electrical engineering we refer to this idea as the pulse-width modulation (PWM) hypothesis (Fig. 1).

**Figure 1.**
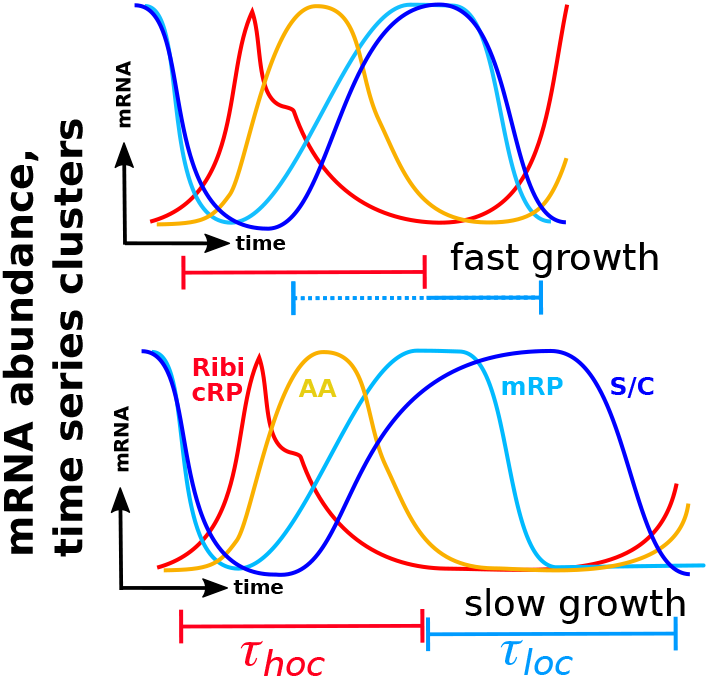
PWM, Pulse-Width Modulation of Transcription: The oscillation period in continuous culture is related to the culture growth rate. At slower growth the period is longer, reflected in a longer LOC phase, while the HOC phase stays approximately constant. The conserved temporal program of transcription is coupled to HOC and LOC phases. Thus, at a longer LOC phase the LOC phase-specific transcript abundances stay high for a longer time. This should lead to a overall higher abundance of LOC phase-specific transcripts (cohort S/C) and lower abundance of HOC phase-specific transcripts (cohort Ribi/RP), and thereby also to higher and lower abundances of the protein products produced (translated) from these transcripts. This is equivalent to the modulation of visually perceived intensity of LED lights by varying the fraction of time they are switched on, i.e., the pulse width.

To test above (non-exclusive) hypotheses, we performed strand-specific RNA sequencing (RNAseq) in high temporal resolution during an unstable state of the short period cycle of the strain IFO 0233. Only a few genes that combine high transcript abundance amplitudes with short protein half-lives are compatible with the JIT hypothesis. These may point to a feedforward control of the transition from catabolic to anabolic flux. However, the bulk of the protein-coding transcriptome codes for long-lived proteins. The duration of the LOC phase transcript abundance peak increased, and the duration of the HOC phase transcript abundance peak decreased within just two cycles of the oscillation. This preceded the transition to a longer period, compatible with the PWM hypothesis. Finally, we present a novel mathematical model of the PWM hypothesis that correctly predicts the correlations of growth-related protein abundances and oscillation periods to growth rate.

## Results and Discussion

### Metabolic Context: Period Drift and a Bifurcation

Previously, stable oscillations have been used to elucidate the transcriptome dynamics of continuously grown yeast (***Klevecz et al., 2004***; ***Li and Klevecz, 2006***). Here we observed more complex transient dynamics, that occurred spontaneously (Fig.2A, S1-S3). We first calculated the oscillation periods and metabolic rates from real-time measurements of the culture (Appendix A, Dataset S1) to characterize these dynamics. The culture cycled between a phase of low oxygen consumption (LOC) and a phase of high oxygen consumption (HOC). The period was 0.6h-0.7h (Fig.2B-C), i.e., the typically observed period for this strain and condition (***Satroutdinov et al., 1992***; ***Murray et al.,2001***). The respiratory quotient, 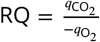, allows to infer details of the catabolic flux. During the LOC phase *RQ* > 1, i.e., cells produced ethanol and excess CO_2_ (fermentation). During the HOC phase, RQ decreased below 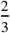, i.e., below the stoichiometry of complete ethanol oxidation (Fig.2D). This is consistent with a re-uptake of ethanol during the HOC phase (***Satroutdinov et al., 1992***) but points to additional contributions to CO_2_ turnover, i.e., an additional uptake of CO_2_ during HOC phase. Proton export (*q*_H_+, Fig.2C, E) peaked in early HOC phase, consistent with a higher intracellular pH during HOC in both short period and long period oscillations (***Keulers et al., 1996a***; ***O’Neill et al., 2020***). The concentration of H_2_S peaked at ≈ 3 **μM** with a sharp increase upon transition to LOC (Fig.2C, F), consistent with its release during amino acid biosynthesis in this transition phase (***Murray et al., 2007***) and its suggested role in population synchronization (***Murray et al., 2003***). The estimated ATP turnover rates (Fig.2D) were in phase with previously measured ATP/ADP ratios, peaking in early to mid HOC phase (***Machné and Murray, 2012***; ***Amariei et al., 2014***). Thus, the overall properties of the oscillations were consistent with previous data. During the whole run, oscillations appeared and vanished spontaneously twice. Both these events were similar. First, period decreased from 0.7h to 0.6h within ≈ 30h (Fig.2B). This period decrease was reflected in a decrease of the LOC phase length, while the HOC phase length even increased. At the end of this transient a sudden bifurcation of the dynamics occured. Afterwards periods were longer with a maximum of 1 h, but the oscillation was unstable and disappeared within a few cycles. This bifurcation was preceded by an increased and phase-shifted peak of CO_2_ release at the transition from HOC to LOC (Fig.2C, E, F). The peak of H_2_S release was delayed, and a novel third phase appeared between the peaks of CO_2_ and H_2_S release. This intermediate phase was purely respiratory at RQ = 1, and all metabolic rates had intermediate values.

**Figure 2.**
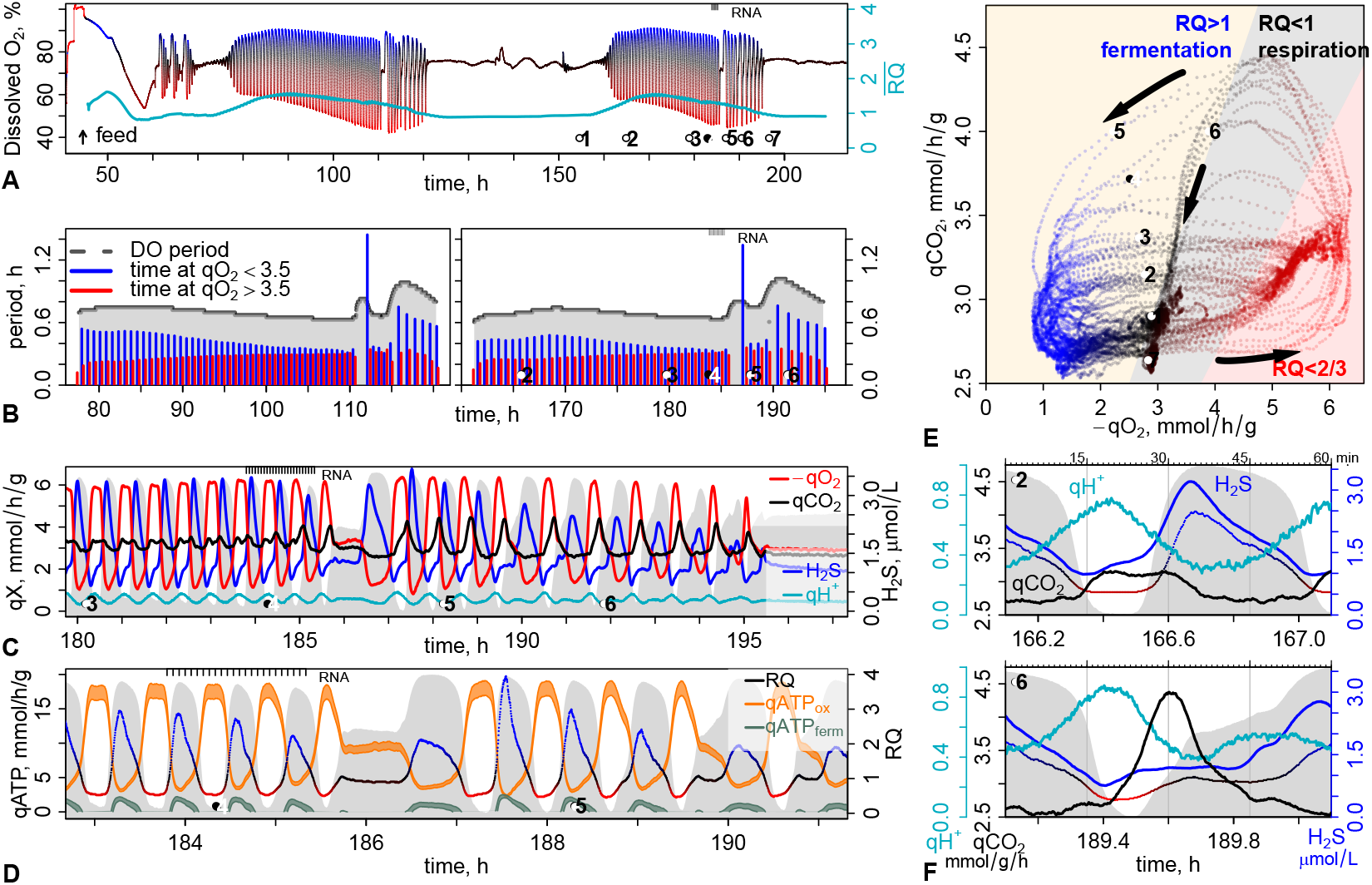
Complex Dynamics: Slow Transients and a Sudden Bifurcation. Metabolic dynamics during continuous culture of the budding yeast strain IFO 0233: panel A shows the full recorded time-series of the culture, and panels B-F zoom in on the time axis; bullet points P1-P7 serve as a guide between panels and are discussed in the text. The gray backgrounds show the dissolved O_2_ concentration (see A for axis) and serves as a reference to oscillation phase. **A:** Dissolved O_2_ (DO) measurement from the start of continuous feeding (dilution rate *ϕ* = 0.089 h^-1^). Line colors are derived from the respiratory quotient RQ (D) and indicate phases of high O_2_ consumption (HOC: red) and low O_2_ consumption (LOC: blue). The cyan line and right axis show the temporal mean 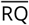, a moving average over ca. 10 h. **B:** The cycle periods were derived from a Wavelet transform of the DO signal and the phase lengths are the time spans of each cycle where oxygen uptake (−*q*_O_2__) stayed below (red, HOC) or above (blue, LOC) 3.5mmol/h/g_DCW_. **C:** Zoom on P3-P6 for measured metabolic rates and concentrations; *q*_O_2__, *q*_CO_2__ and H_2_S were measured in the offgas of the reactor, corrected for the measurement delay and H_2_S concentration was derived via its solubility. Proton export (*q*_H_+) was calculated from the NaOH addition rate. **D:** Zoom on P4-P5 for calculated rates. The respiratory quotient (RQ) and ATP production rates by respiration (*q*_ATPox_) or by fermentation (*q*_ATPferm_) were calculated from *q*_O_2__ and *q*_CO_2__ (Eq. S9-S14 in Appendix A). The RQ color gradient serves as a reference in (A, E,F). **E:** Phase portrait of *q*_O_2__ and *q*_CO_ over the time range indicated by bullet points in (A); points are colored by RQ (D) and in 10 s resolution; background colors indicate RQ ranges and arrows indicate time direction. **F:** One-hour snapshots at different times (bullet points 2 and 5). Data are indicated by colored axes and labels, except for RQ which is shown without axis but color-coded (red-black-blue) as in D and E. All reactor data is available as Datafile S1.

In summary, our experiment reflects the previously studied oscillation of the IFO 0233 strain, however, we describe complex transient dynamics that appeared twice. Emergence and disappearance of the oscillations could originate from a loss of oscillatory metabolic dynamics in single cells. We favor the alternative hypothesis that culture level oscillations result from a synchronization between individually oscillating single cells (***Silverman et al., 2010***). During the synchronous phases, the oscillation period first drifted slowly to a minimum of ≈ 0.6h; then system dynamics rapidly changed (bifurcated) to an unstable state with a longer period (≈ 1 h) and an intermediate phase that was purely respiratory (RQ ≈ 1). Low but purely respiratory activity at RQ ≈ 1 is characteristic of the LOC phase in CDC-coupled (long period) systems (***Münch et al., 1992***). The bifurcation was accompanied by the appearance of a pulse of CO_2_ release before, and a delayed pulse of H_2_S release after the intermediate RQ ≈ 1 phase. We interpret the period drift and sudden transition as an imbalance between catabolic and anabolic flux.

### Transcriptome Oscillation: A Universal Temporal Program

Numerous time series of the protein-coding transcriptome have revealed a universal temporal program of defined transcript cohorts but with periods ranging from 40 min to 7.5 h (***Machné, 2017***). Transient states of the oscillation or non-coding transcription have not been studied. We sampled for RNAseq analysis every 4min for 2.5 cycles, just preceding the bifurcation of system dynamics (P4 in Fig.2). The strand-specific sequencing reads were mapped to the reference genome (strain S288C, R64-1-1), yielding reads for 76 %of the genome (Fig. S5A). A similarity-based segmentation algorithm (***Machné et al., 2017***) yielded ca. 37k segments (Fig. S5D), each a putative individual transcript. All segments were classified by their oscillation p-values, calculated with the rain package (***Thaben and Westermark, 2014***), and by their overlaps with annotated genome features (Tab. S2). 4,489 segments were classified as open reading frame (ORF) transcripts; 3,378 of these showed oscillation and reproduced the previously characterized temporal sequence (Fig.3A; ***Machné and Murray*** (***2012***)). Oscillating non-coding (811 of 9,051) and antisense (232 of 569) segments predominantly peaked in the LOC phase. Very short and weakly expressed segments were removed from further analyses, the remaining 11k segments (Fig. S5G-I) were clustered into ten co-expressed clusters, and these were sorted and colored by their peak phase (Fig. S6-S8). These ten clusters can be further classified (Fig. S6C) into two groups of five clusters each. The first group (Fig.3B) comprises of longer segments with high amplitudes, and most assigned to protein-coding genes (Fig. S7). The second group (Fig.3C) contains shorter and weakly expressed segments with lower amplitudes, mostly non-coding and peaking during the LOC phase (Fig. S8).

**Figure 3.**
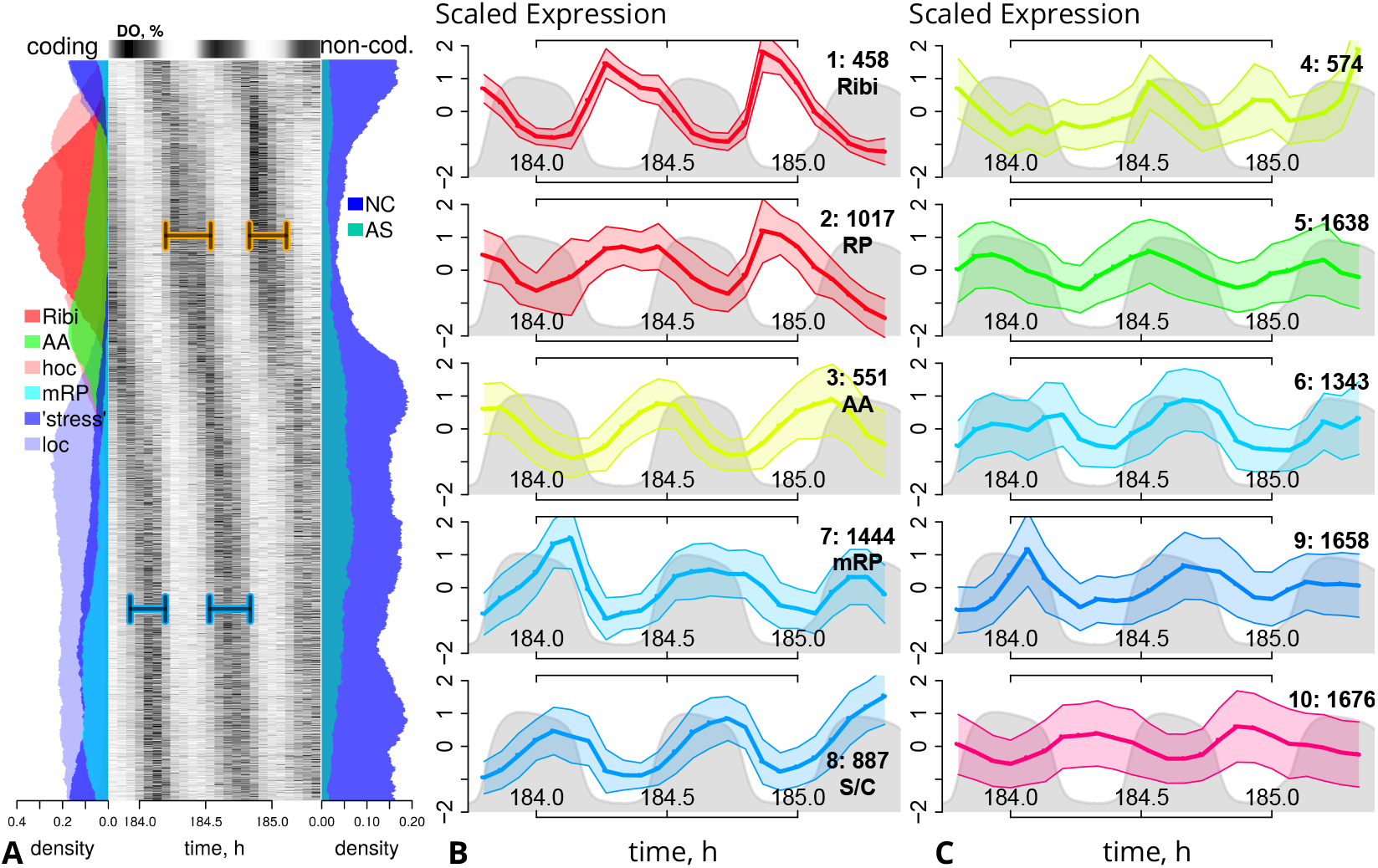
RNAseq Time Series Clustering. **A:** Phase-ordered heatmap of the time-courses of segments with oscillating abundance levels (6344 segments at *p*_rain_ < 0.05, Fig. S5E)). The dissolved O_2_ (DO, %) is shown as a color gradient (black: high DO) on the top axis. Left and right panels show local densities (circular moving average of counts over 1/10 of the total number) of segments overlapping with previously defined (***Machné and Murray, 2012***) classes of coding genes (left: Ribi, ribosomal biogenesis and cytoplasmic ribosomal proteins;AA: amino acid synthesis;mRibi: mitochondrial genes, incl. ribosomal proteins;stress: catabolic and protein homeostasis genes), or non-coding segments (right: AS, antisense to ORF;NC, no overlap with any annotated transcribed feature). **B:** Time series of the five major periodic co-expression clusters. Segment time-courses (mean RPM) were scaled to a mean of 0 and divided by their standard deviation. The mean of each cluster is shown as a solid line with points indicating the sampling times, and standard deviations are shown as transparent ranges;the legends indicate the cluster label, the number of segments in the cluster and the posterior functional cohort assignment. The gray background indicates the dissolved O_2_ (DO) concentration. **C:** Time series for the cluster 4, 5, 6, 9, 10, which comprise mostly non-coding segments; plotted as described for (B).

#### A Conserved Temporal Program Runs at Different Time Scales

Gene Ontology (GO) enrichment analysis of the protein coding cohorts (Fig.4A and S9) recapitulates previous data (***Klevecz et al., 2004***; ***Machné and Murray, 2012***). The ribosomal biogenesis regulon (***Jorgensen et al., 2004***) peaks in early to mid HOC phase (cluster 1: `Ribi`), followed by clusters encoding for cytoplasmic ribosomal proteins (cluster 2: `cRP`), and amino acid biosynthetic pathways (cluster 3: `AA`) at the transition to LOC phase. During the LOC phase, mitochondrial proteins, including mitochondrial ribosomal proteins (cluster 7: `mRP`) are co-expressed with a regulon associated with stress response (***Gasch et al., 2000***; ***Brauer et al., 2005***) and G1 phase (***O’Duibhir et al., 2014***). The latter comprises of proteins involved in the general stress response (chaperones) and in carbohydrate, fatty acid and protein catabolism (cluster 8: S/C). We used this clustering to reanalyze eight data sets from different strains and conditions and with periods ranging from 40min to 7.5 h (Fig. S10–S11, data from ***Li and Klevecz*** (***2006***); ***Tu et al.*** (***2005***); ***Slavov et al.*** (***2011***); ***Chin et al.*** (***2012***); ***Kuang et al.*** (***2014***); ***Wang et al.*** (***2015***); ***Nocetti and Whitehouse*** (***2016***)). This meta-analysis reveals common patterns. A temporally constrained program (0.5h–2h) leads from Ribi/cRP *via* AA to `mRP`, ending with the transition from HOC to LOC phase. Increases of the total period are mostly reflected by increased duration of the LOC phase and the associated `S/C` cohort expression. The same temporal program can be observed in six distinct cell cycle arrest & release experiments (Fig. S12) (***Orlando et al., 2008***; ***Bristow et al., 2014***).

**Figure 4.**
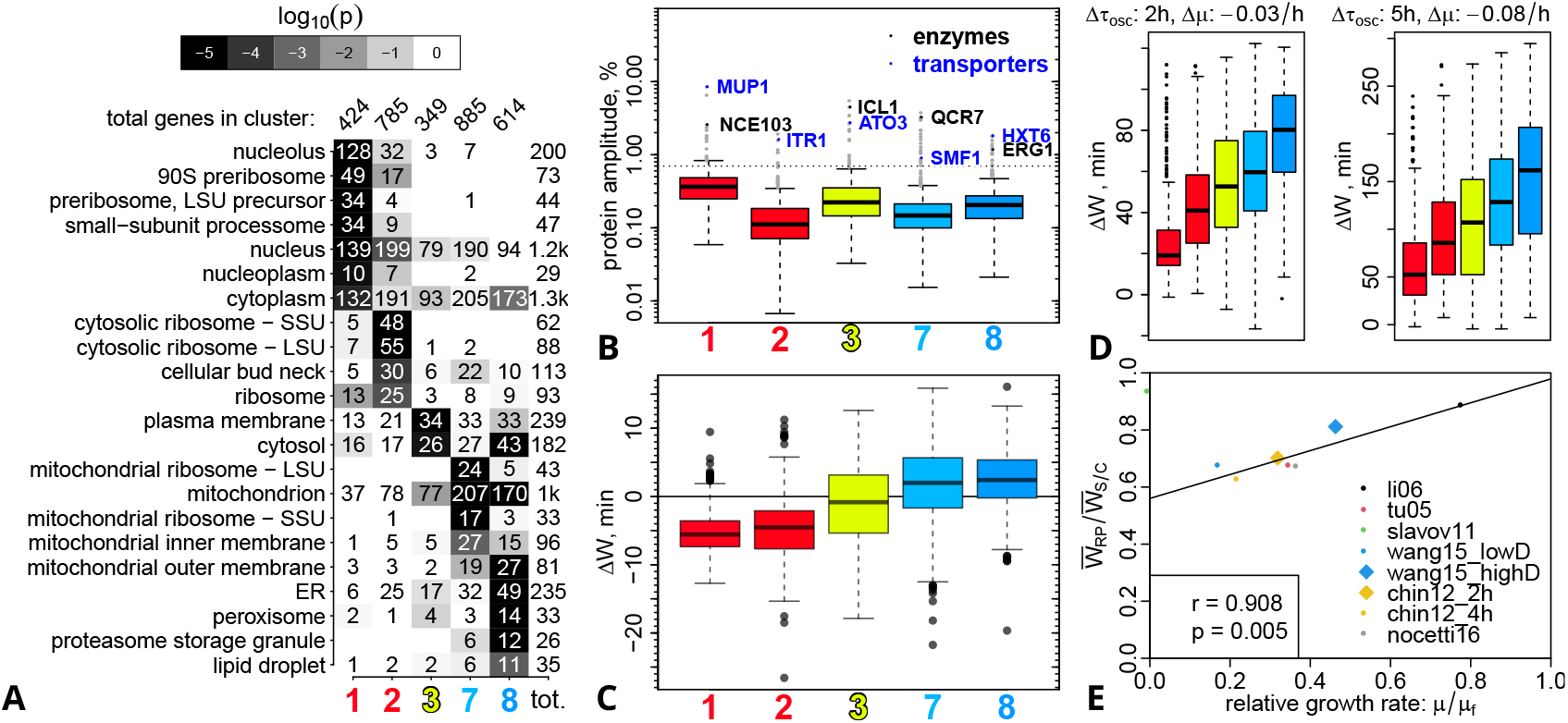
A Universal Temporal Program. **A:** Sorted cluster enrichment profiles for the GO category “cellular component”. The p-values were calculated by cumulative hypergeometric distribution tests and are shown as gray-scale. Only categories with *p* < 0.001 (white text) and more then 10 genes in one of the clusters are shown. See Figure S9 for all clusters and GO categories. **B:** Boxplots of cluster distributions of predicted relative protein amplitudes (*A*_P_, in % of their mean abundance), estimated from transcript amplitudes and protein half-lives (Fig. S13). The horizontal line indicates the the top 100 oscillators listed in Table S3. The top predicted oscillators of class enzyme or membrane transporter of each cluster are indicated. **C:** Boxplots of cluster distributions of transcript abundance peak width differences Δ*W* = *W*_2_ – *W*_1_ between the second and first full expression cycle. Figure S16 provides details on the calculation. **D:** Boxplots of the transcript abundance peak width differences between two experiments from the same culture but at different growth rates (left: ***Chin et al.*** (***2012***), right: ***Wang et al.*** (***2015***)). See Fig. S10B for raw peak widths. **E:** Peak width ratio vs. growth rate for all experiments analyzed in Figure S10B. The ratio of the mean peak widths ofcluster 2 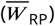 and cluster 8 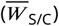 is correlated to the strain-specific relative growth rate (*μ/μ_f_*, see Fig. S10B for *μ* and Tab. S5 for *μ_f_* values). Experiments are indicated by the first author and year in the legend. The line indicates a linear regression, and *r* and *p* are the Pearson correlation and p-value, all calculated without the outlier at *μ* = 0h^-1^ (`slavov11`) which was taken from an oscillation at the end of a batch growth phase on ethanol medium.

### Testing Hypotheses: Putative Functions of the Temporal Program

Next, we analzyed the two hypotheses on putative functions of this universal temporal program; the just-in-time production (JIT) and the pulse-width modulation (PWM) hypothesis.

#### Carbonic Anhydrase and the Glyoxylate Cycle are Novel Feedback Candidates

The temporal order of mRNA abundances makes intuitive sense as a just-in-time gene expression program (JIT) coordinated with metabolic events. However, oscillations on transcript level are dampened by long protein half-lives (***Lück et al., 2014***). Thus, we estimated relative protein amplitudes (Fig.4B, S13A-C) from our RNA abundance time series and from protein half-life data by ***Christiano et al.*** (***2014***), using a mathematical model of periodic gene expression by ***Lück et al.*** (***2014***). Most proteins are predicted to vary by 0.1 %-0.5 %of their mean abundance (Fig.4B and S13C). Only 23 proteins have predicted relative protein amplitudes ≥2%; and oscillators are enriched in the `Ribi` and `AA` cohorts (Fig. S13C). These low amplitudes probably do not have a strong effect on metabolic dynamics, but the model is based on sine approximations of transcript time series and protein half-lives measured in asynchronous conditions; it may understimate amplitudes and it completely neglects potential effects of induced protein degradation and post-translational modifications. Thus, we tested our predicted against measured protein amplitudes in a long period oscillation (***O’Neill et al., 2020***). The genome-wide correlation between these amplitude sets was weak but significantly positive (Fig. S14). However, the top oscillator estimates of both data sets overlapped (Fig. S13D, E). Notably, 60 of the top 100 oscillators in our analysis were not detected in the proteomics measurement. These include several transcription factors (e.g. BDF2, CLB2, GZF3, MET28, SWI5, MSN4) which are known to be expressed at low levels. Thus, our analysis reveals putative oscillators that are potentially missed by proteomics analysis.

Both top oscillator lists share cell wall proteins, nutrient transporters, and metabolic enzymes. Several enzymes of the sulfate uptake pathway (MET genes) are expressed in Ribi and peaked prior to the pathway intermediate H_2_S at the HOC/LOC transition (Fig.2). The carbonic anhydrase (NCE103, in `Ribi`) catalyzes the interconversion of carbon dioxide and bicarbonate 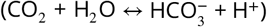, and is essential in aerated cultures (***Aguilera et al., 2005***). During the second sampled cycle the Ribi cohort was downregulated early (Fig.3A, Fig.4C) and this correlated with the appearance of the CO_2_ and the delay of H_2_S release pulses at transition to LOC (Fig.2C–F). Both, CO_2_ and H_2_S, were previously suggested to contribute to population synchronization (***Keulers et al., 1996a***; ***Murray et al., 1999, 2003***), and both are substrates of biosynthetic metabolism. However, the strongest synchronizing activity was found for the acetaldehyde (***Murray et al., 2003***), a futile intermediate offermentation or, more generally, of overflow metabolism around the pyruvate node of metabolism, between glycolysis, respiration and biosynthesis (***Pronk et al., 1996***; ***Sonnleitner and Käppeli, 1986***). The switch from catabolism in HOC phase to anabolism at the transition to LOC phase likely involves regulation around this central node of metabolism. We find several biosynthetic enzymes among the top 100 predicted oscillators (Fig. S15), most notably three enzymes of the glyoxylate cycle (ICL1, CIT2, MDH2, all in the `AA` cohort), a shorter and purely biosynthetic version of the tricarboxylic acid cycle. It is for example required to synthesize glucose, when ethanol is the only carbon source. This cycle is autocatalytic (***Barenholz et al., 2017***) and serves as metabolic switch in response to changes in carbon source (***Nakatsukasa et al., 2015***).

All discussed pathways also appear in the proteome-based list of top oscillators (Fig. S13D,E, S14), supporting their general relevance for metabolic oscillations. As outlined in Figure S15, these short-lived enzymes could be involved in gating the transition from the catabolic to the anabolic phase of the cycle.

#### Resource Allocation by Pulse-Width Modulation (PWM)

Most proteins are too stable for an effect of oscillatory transcript on protein abundances.***Slavov and Botstein*** (***2011***) and ***Burnetti et al.***(***2016***) suggested an alternative interpretation of periodic transcription. Variation of the relative times spent in HOC phase- and LOC phase-specific transcription states could serve to tune steady-state protein abundances. The LOC phase duration decreases with increasing growth rates, while HOC phase duration remains approximately constant or even slightly increases (***von Meyenburg, 1969a***; ***Strässle et al., 1989***; ***Slavov and Botstein, 2011***; ***Burnetti et al., 2016***; ***O’Neill et al., 2020***). This would lead to a higher relative biomass fraction of proteins from HOC phase-specific transcripts, i.e. of the `Ribi` and the `cRP` cohorts. During our experiment a similar shift of the relative times spent with HOC or LOC phase-specific expression occured (horizontal bars in Fig.3A). We quantified and compared the peak widths between the two cycles (Fig. S16). The `S/C` cohort peak width increased on average by ≈ 3 min, while the Ribi cohort peak width decreased by ≈ −5 min (Fig.4C). This occured without comparable changes of the duration of HOC and LOC phases, i.e., the transcription was not merely an output of respiratory dynamics. Thus, the relative duration of expression phases can be adapted rapidly and affect the metabolic dynamics in subsequent cycles.

So we next looked for evidence of PWM of transcription in the previous data sets and calculated peak widths for all transcripts (Fig. S10B). When growth rate was decreased in dilution rate shift experiments (***Chin et al., 2012***; ***Wang et al., 2015***) the period increased, as expected. Most transcript abundance peak widths increased with period, but this increase was significantly higher for the LOC-phase specific cohorts (Fig.4D). Thus, the peak widths of HOC phase-specific and LOC phase-specific co-expression cohorts indeed changed with growth rate. The oscillation periods tend to reach a minimum towards a strain-specific critical growth rate (*μ_f_*) where fermentation sets in (***Burnetti et al., 2016***; ***Machné et al., 2017***). We calculated the mean peak widths of the RP cohort (cluster 2) and the `S/C` cohort (cluster 8), and a relative growth rate for each experiment, i.e., the growth rate (dilution rate) of the continuous culture divided by the strain-specific critical growth rate 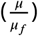. This reveals a good correlation between the `RP` to `S/C` peak width ratio and the relative growth rate of the cultures (Fig.4E). The only outlier is the transcriptome data taken at the end of a batch growth phase, i.e. at *μ*≈ 0h^-1^, on ethanol medium (***Slavov et al., 2011***).

An increase of ribosome content is directly and causally related to higher growth rates, constituting a fundamental principle of microbial growth physiology (***Schaechter et al., 1958***; ***Waldron and Lacroute, 1975***; ***Koch, 1988***; ***Scott et al., 2010***). This relation is reflected in continuous changes of relative abundances of different transcript and protein classes with growth rate (***Brauer et al.,2005***; ***Airoldi et al., 2009***; ***Molenaar et al., 2009***; ***Metzl-Raz et al., 2017***). No mechanism for this continous variation of gene expression is known in eukaryotes. A temporal regulation, via continuous changes of the relative durations of LOC and HOC phases generates this relation in synchronously oscillating continous culture. Even in asynchronous cultures, individual cells appear to oscillate (***Silverman et al., 2010***), thus this mechanism is likely general.

### The PWM Model Explains Period and Proteome Relations to Growth Rate

#### Consistent Prediction of Transcript and Protein Abundances

Next, we set out to explore the predictive power of the PWM hypothesis. In short, we assume a step function of transcriptional activity, such that genes are transcribed at maximal rate during their respective expression phase (HOC or LOC) and not transcribed in the other phase. The mean concentrations (over time) of an mRNA that is transcribed only in HOC phase (*R_hoc_*), and of its protein product (*P_hoc_*) are:

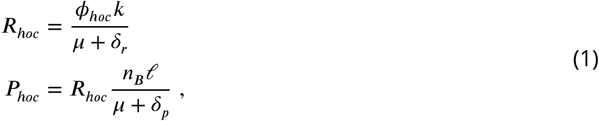

where *ϕ_hoc_* = *τ_hoc_/τ_osc_* is the fraction of the total period (*τ_osc_*) spent in HOC phase (*τ_hoc_*), *k* and *ℓ* are transcription and translation elongation rates, *n_B_* is the ribosome density (ribosomes per RP mRNA); *δ_r_* and *δ_p_* are the mRNA and protein degradation rates; and *μ* is the culture growth rate. The same model can be used for LOC phase-specific genes, with transcription restricted to *ϕ_loc_* = 1–*ϕ_hoc_*. See Appendix B for a detailed derivation of the model.

The period *τ_osc_* decreases with increasing growth rate (Fig.5A, S17A). This period decrease is reflected in a decrease of the time spent in LOC phase (*τ_loc_*), while the duration of the HOC phase stays approximately constant or even slightly increases (***von Meyenburg, 1969a***; ***Strässle et al.,1989***; ***Bellgardt, 1994***; ***Slavov and Botstein, 2011***; ***Burnetti et al., 2016***; ***O’Neill et al., 2020***). Similarly, *τ_loc_* decreased with period *τ_osc_*, while *τ_hoc_* changed less and in opposite direction during our experiment (Fig.2B). We thus estimated a *ϕ_hoc_* = *f*(*μ*) from data from the IFO 0233 strain (Fig. 5A, ***Murray et al.*** (***2001***)), used our classification into HOC phase and LOC phase genes (Fig.3,4), and collected gene-specific parameters for the production and degradation rates for each gene (Fig. S18). The model assumes that all regulation occurs through initiation of transcription at a maximal rate in HOC or LOC phase. The maximal transcription and translation rates merely depend on the gene and proteins lengths, while ribosome densities (per mRNA) and degradation rates are derived from genome-wide experimental data (Tab. S5). These assumptions and data allowed to estimate growth rate-dependent mean transcript and protein abundances from Eq.1 for 1,197 genes (Fig.5C-F, S21). To estimate the predictive power we calculated the slopes 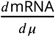, and find a good correlation (Spearman’s *ρ* = 0.66) with the slopes reported for 35 signature genes of the Universal Growth Rate Response (UGRR) model (Fig.5D, ***Airoldi et al.*** (***2009***); ***Slavov and Botstein*** (***2011***)). Next, we calculated slopes for absolute transcript counts measured in chemostat cultures at different growth rates by ***Xia et al. (2022)***. The correlation is overall weak (*ρ* = 0.35, Fig. S21C), but better for smaller gene sets from a more stringent consensus classification (*ρ* = 0.74, Fig. S21D). Similarly, we found good overall agreement of the relative proteome fractions at different growth rates of gene groups selected by ***Metzl-Raz et al.*** (***2017***) (Fig.5E). For example, the pro-teome fraction of mitochondrial genes decreases, while the fraction of genes involved with translation increases with growth rate, reflecting measurements (***Metzl-Raz et al., 2017***). The correlation with measured growth rate-slopes of proteins (***Xia et al., 2022***) were higher than for transcripts (*ρ*= 0.42, and *ρ* = 0.78 for the consensus set; Fig. S21G,H). However, the strongest contribution to these correlations comes from our accurate classification into HOC and LOC phase genes, while the correlation for the HOC phase-specific transcripts was even negative (Fig. S21C).

**Figure 5.**
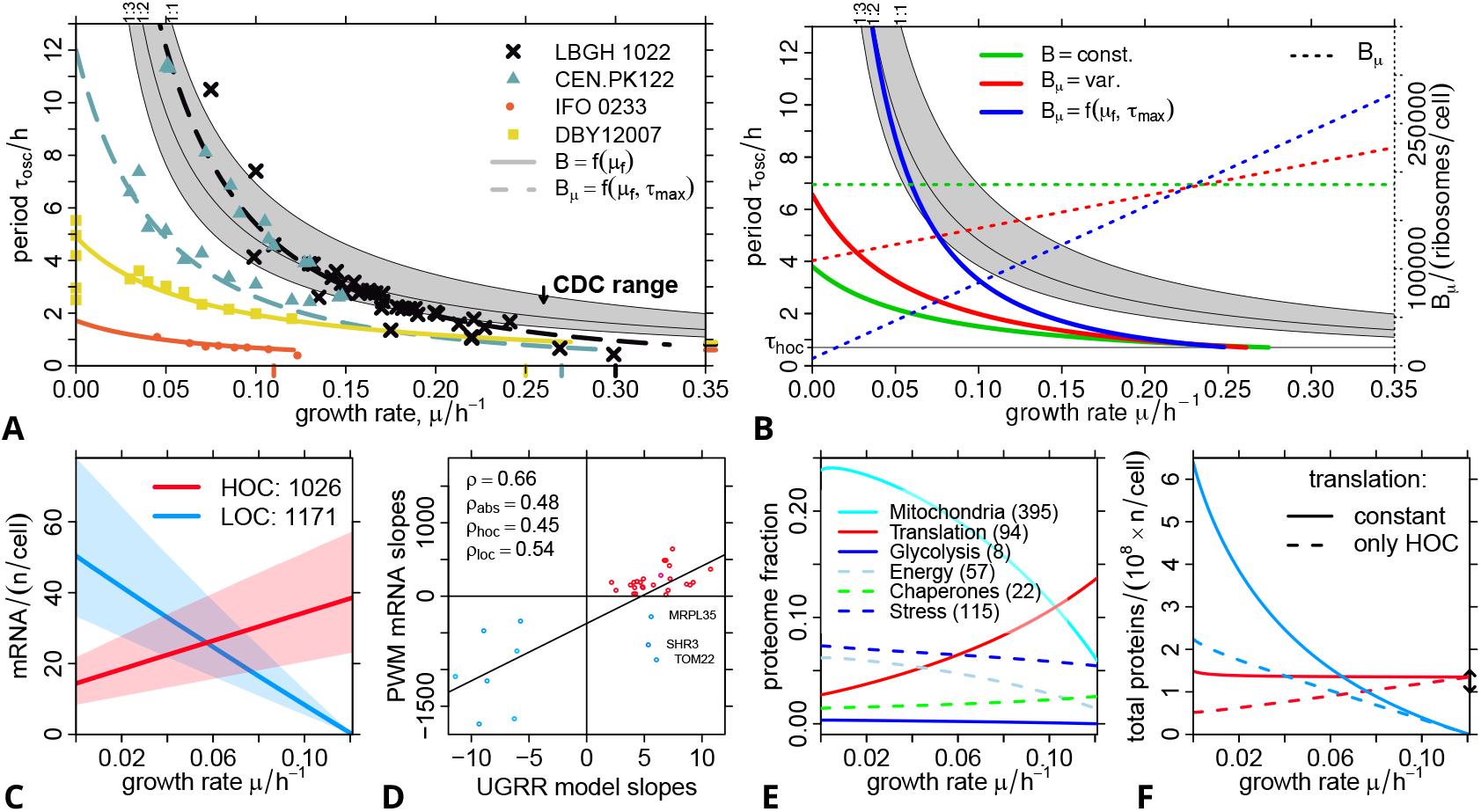
The PWM Model. **A:** Oscillation periods are non-linearly related to growth rate, here shown for four different strains (colored points, cf. Fig. S17A). The periods expected from partial CDC-synchronization (CDC range) in modes 1:1,1:2 and 1:3 are shown as black solid lines (***Bellgardt, 1994***), via Eq. S31-S32. The PWM model (colored lines, Tab. S6) can re-produce the observed periods, incl. at *μ* → 0, and the relation to the strain-specific critical growth rate *μ_f_* (colored ticks on the x-axis). Solid lines indicate a PWM model with constant ribosome concentration calculated via *μ_f_*, and dashed lines with linearly increasing ribosome concentration and the additional assumption of a *τ_max_*; *τ_hoc_* (colored ticks on the right y-axis) was manually adjusted. **B:** Periods predicted by the base model (Eq.2, solid green line) and the extended model with variable ribosome concentration *B*(*μ*) (Eq. S25, solid red line), with parameters from Table S5. The colored dashed lines are the ribosome concentrations (right y-axis) used for each model. Alternatively, ribosome parameters can be estimated via the *μ_f_*-constraint (Eq. S26, solid blue line). **C:** Median (lines) and 25%/75%quantiles (transparent range) of all mRNA abundances predicted by the PWM model (Eq.1) from *ϕ_hoc_* (IFO 0233 parameters in (A)), and from gene-specific production and degradation rates (Tab. S18B), and classification to either HOC (clusters 1, 2 and 10) or LOC (clusters 6, 7, 8 and 8) phase. The legend indicates the number of genes for which all data was available. **D:** Comparison of the mRNA slopes, derived from a linear regression of the data in (C), and the slopes provided for signature genes of the UGRR model (***Slavov and Botstein, 2011***). All data required for the PWM-based prediction was available for the shown 35 of 58 signature genes. Gene names are provided for the outliers, two mitochondrial and one ER-associated. The straight line is a linear regression, and *ρ* is the Spearman correlation, *ρ_abs_* removes the influence of the classification by taking the absolute slopes in both data sets, and *ρ_hoc_* and *ρ_loc_* are correlations calculated for only the HOC- or LOC-specific signature genes (red and blue point symbols). **E:** Fractions of the total protein abundance predicted by the PWM model (Eq.1) for the gene lists used to analyze proteome fractions in ***Metzl-Raz et al.*** (***2017***); gene numbers in brackets. **F:** Total protein abundances predicted by the PWM model for HOC and LOC phase genes; calculated without (solid lines, Eq.1) or with (dashed lines, Eq. S27) an additional restriction of translation to HOC phase. The vertical black arrow indicates the total protein content estimation by ***Milo*** (***2013***). All rates required for mRNA and protein prediction are available in Dataset S3.

The model neglects all other types of regulation such as targeted degradation, or intrinsic bias such as sequence-dependent differences of elongation rates; thus, it is not suprising that on a genome-wide scale the predictive power is weak. Appendix B.6 discusses potential reasons for these discrepancies. A more fundamental problem of the model is that translation is unlimited. The total protein abundance increases strongly at *μ* →0, while it is very close to estimates from experimental data (***Milo, 2013***) at high *μ* (Fig.5F). As outlined in Appendix B.5, previous data point to a pulse of translational activity at the HOC-to-LOC transition. The majority of ATP synthesis occurs in HOC phase; as a simple approximation, we restricted the translation of all transcripts (HOC and LOC phase) to HOC phase (Eq. S27). This reduced the total protein abundance at *μ* → 0 to about twice the estimate for cells in exponential growth (***Milo, 2013***), thus into a more realistic range.

#### The PWM Model Predicts Oscillation Periods

We further noted, that the model yields a strict constraint between oscillation parameters, the life cycle rates and concentration of proteins, and growth rate. Ribosomal proteins (RP) are (a) transcribed within the HOC phase clusters (***Klevecz et al., 2004***; ***Machné and Murray, 2012***) (Fig.4A, D), and (b) their relative fraction of total biomass increases with growth rate (Fig. S18C-E, (***Waldron and Lacroute, 1975***)). Thus, we can use this constraint to predict oscillation periods from measured ribosome concentrations and life cycle parameters. Assuming that each RP is associated with one ribosome (Appendix B.2), we get:

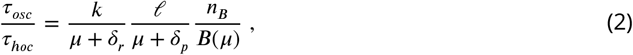

where *B*(*μ*) is the total concentration of (cytoplasmic) ribosomes, and all other parameters refer to an average RP (Tab. S5, Fig. S18A). Remarkably, the collected literature parameters already yield (i) realistic periods and (ii) the non-linear dependence of periods on growth rates (green line, Fig. 5B). Linearly varying the ribosome concentration with growth rate (Fig. S18C) makes the period function steeper (red line, Fig.5B).

Experimentally observed periods reach a minimum towards the strain-specific growth rates *μ_f_*, where yeast metabolism switches from purely respiratory to respiro-fermentative metabolism of glucose (***Burnetti et al., 2016***; ***Machné, 2017***). In the IFO 0233 strain, fermentation sets in early, at growth rates *μ_f_* = 0.11 h^-1^-0.15h^-1^ (***Hansson and Häggström, 1983***; ***Satroutdinov et al., 1992***) consistent with its short period cycles. This constraint allows to estimate strain-specific values for the RP and ribosome-related parameters via published values for *μ_f_* (Appendix B.3, Fig.5A, S17D, Tab. S6). The model with variable ribosomes is required to fit data from the two strains with longer periods (CDC range), or, alternatively, very low degradation rates (Fig. S17C). This pattern is confirmed when fitting Eq.2 separately to 20 independent data sets (Fig. S19, S20). Long period data sets require to set at least one of the degradation parameters (*δ_r_*, *δ_p_*) to 0. This may be due to a phase-locking with the CDC (gray areas in Fig.5A, B), where the HOC phase aligns with the budding phase of the CDC and LOC phase is purely respiratory (RQ ≈ 1) and corresponds to the G1 phase of the CDC (***Münch et al., 1992***). When such phase-locking with the CDC occurs, PWM and oscillation parameters may not be directly coupled anymore.

Previously suggested models based on partial synchrony of the asymmetric cell division cycle fit long period data well (***Bellgardt, 1994***; ***Hjortso and Nielsen, 1995***; ***Duboc and von Stockar, 2000***). However, these models can not account for oscillations in batch culture and without division (***Mochan and Pye, 1973***; ***Murray, 2004***; ***Slavov et al., 2011***) and for periods that are longer than the culture doubling time (***Heinzle et al., 1983***; ***Porro et al., 1988***). ***Burnetti et al.*** (***2016***) suggested a purely empirical model for these relations. The PWM model is the first mechanistic model of the oscillation that can account for all experimentally observed periods; although only with unrealistic parameter choices for long periods. Future work based on this novel theoretical framework should explicitly account for energetic constraints on the protein synthesis capacity during the cycle, and could explore the effects of additional regulatory mechanisms or systematic differences in production and degradation rates.

## Conclusion

The phenomenon of metabolic auto-synchronization in budding yeast continuous culture was instrumental for the clarification of the asymmetric CDC of budding yeast (***Küenzi and Fiechter, 1969***; ***von Meyenburg, 1969a***). The discovery of stable short period cycles in the distillery strain IFO 0233 (***Satroutdinov et al., 1992***) fortified early indications (***von Meyenburg, 1969b***; ***Mochan and Pye,1973***) that the system is more than just synchronization of the CDC. Here, we first explored the complexity of dynamics observable in budding yeast continuous culture, a long-term transient and a sudden bifurcation. We then tested the two main hypotheses on putative functions of the periodic transcriptome (JIT and PWM).

We presented four independent lines of evidence in support of the PWM hypothesis ***(Burnetti et al., 2016***; ***Slavov and Botstein, 2011***): (i) transcript abundance peak widths changed as predicted in two dilution rate shift experiments, (ii)the relative peak widths correlated very well to the relative growth rate, i.e., the growth rate divided by the strain-specific critical growth rate, (iii) the PWM model predicts measured growth rate-dependent transcript and protein abundances reasonably well, despite its simplicity, and (iv) the PWM model predicts the dependence of oscillation periods on growth rate. The coupling is consistent over periods ranging from 40min to 7.5h. We further note that circadian biology faces a similar problem, low protein abundance amplitudes despite significant transcript abundance oscillations (***Lück et al., 2014***; ***Wang et al., 2018***; ***Krahmer et al.,2021***; ***Karlsen et al., 2021***). While the period is fixed, seasonal variation of light/dark cycle phase lengths could mediate PWM-based control of steady state protein abundances.

And finally, the prediction of periodic proteins (JIT analysis) and the metabolic dynamics during our experiment underpin previous data on H_2_S and CO_2_ as population synchronizers (***Keulers et al., 1996a***; ***Murray et al., 2003, 2007***). The accumulating evidence suggests that the involved pathways could gate the switching from catabolic to anabolic flux at the transition from HOC phase to LOC phase (Fig. S15). The metabolic mechanisms behind CDC-coupled long period oscillations were considered to lie in a cycle of glycogen build-up during LOC phase and mobilization during HOC phase, where the respiratory electron transport chain becomes limiting and overflow metabolism at the pyruvate node (ethanol, acetate or acetaldehyde accumulation and secretion) induces the switch to LOC phase and synchronizes the culture (***Küenzi and Fiechter, 1969***; ***Strässle et al., 1989***; ***Münch et al., 1992***). However, glycogen content oscillates at low amplitude and peaks in the wrong phase in IFO 0233 (***Satroutdinov et al., 1992***) and glycogen is not produced during oscillatory growth on ethanol-based medium (***Keulers et al., 1996b***). Thus, the glycogen cycle model is either wrong or not generally valid. The next big question is thus to clarify the metabolic mechanisms behind switching between the HOC and LOC phases of this cycle. What is the nature of the metabolic limitation in continuous culture, and how does it determine the relative lengths of the phases?

## Supporting Information and Data

The RNA sequencing reads are available at ArrayExpress (http://www.ebi.ac.uk/arrayexpress/, ***Athar et al***. (***2019***)) with accession number E-MTAB-11901.

### Supporting Information File

Appendices A (calculation of metabolic rates from bioreactor online measurements), B (detailed formulation of the PWM model), and all Supporting Figures.

### Dataset S1

Reactor data, including all calculated rates and RNAseq sampl timess.

### Dataset S2

All 36,928 segments reported by `segmenTier`, incl. genome coordinates, cluster labels, read-counts, oscillation values (amplitude *A*_2_, phase *ϕ*_2_, p-value *p*_rain_), coding gene and SUT overlaps, and all time points, using the sampling IDs (2–25) indicated in the reactor data.

### Dataset S3

Data for 3,849 coding genes that overlap with a segment with *J* > 0: relative mRNA and protein amplitudes and protein half-lives for prediction of protein amplitudes (Fig. S13), and production and degradation rates as used for period, mRNA and protein abundance predictions, incl. cluster associations and classification as RP gene (Fig. S18A/B, Tab. S5).

## Materials and Methods

### Strain History

Kuriyama’s lab first reported oscillations in continuous culture of the *Saccharomyces cerevisiae* strain IFO 0233 (***Satroutdinov et al., 1992***). The strain number is from the Japanese culture collection NBRC and is identified there as “`Distillery yeast Rasse II`”, “accepted” in 1941, and as ATCC 560 in the US American culture collection. These strains can be traced back to the “Brennereihefe, Rasse II” isolated as “Hefe 128” axenic culture by Paul Lindner at the Berlin Institut für Gärungsgewerbe in 1889 from samples of a distillery in Gronowo (West Prussia, now Poland) which obtained their yeast from a dry yeast supplier in the city Thorn (now Toruń, Poland) (***Lindner, 1895***). The strain and its descendant “Rasse XII” became commercially successful distillery strains within hybrid formulations (“Rasse M”), and was at the time an intensively studied strain in basic research, *e.g*., in the search for the nature of “bios” (***Lindner, 1919***).

### Continuous Culture

#### Pre-Culture

*Saccharomyces cerevisiae* (strain IFO 0233) were maintained on yeast nitrogen base agar plates (2%glucose, 1.5%agar; Difco, Japan) at 4 °C, sub-cultured from frozen stock cultures (−80 °C; 1mL; 15 %glycerol; 5 × 10^8^cells). Pre-cultures were inoculated into Yeast Extract Peptone Dextrose media (10mL; 1 %yeast extract, 2%peptone, 2%glucose) and grown at 30°C in an orbital incubator (200rpm) for 24 h.

#### Continuous Culture Medium & Inoculation

The culture medium consisted of D-glucose (20 gL^-1^), (NH4)2SO4(5 gL^-1^), KH2PO4(2gL^-1^), MgSO4.7H2O (0.5gL^-1^), CaCl2.2H2O (0.1gL^-1^), FeSO4.7H2O (20mgL^-1^), ZnSO4.7H2O (10mgL^-1^), CuSO4.5H2O (5mgL^-1^), MnCl2.4H2O (1 mgL^-1^), 70%H2SO4, (1 mLL^-1^), Difco yeast extract (1 gL^-1^) and Sigma Antifoam A (0.2 mLL^-1^). All chemicals were supplied by Wako Pure Chemical Industries Ltd., Japan. The medium prepared with this recipe has a pH of ca. 2.5 which allows for autoclaving of media with both sugar and ammonium without browning (caramelization) and further avoids precipitation of salts in feed medium bottles during continuous culture. A custom-built bioreactor as outlined below was filled with 0.635L of medium and autoclaved (121 °C; 15min). Aeration (0.15Lmin^-1^), agitation (750rpm), and temperature (30°C) and pH (3.4) control were switched on, until the system was equilibrated. Then, the dissolved oxygen probe was 2-point calibrated by flushing with pure nitrogen (0%) and switching back to air (100%). The equilibrated and fully calibrated reactor was inoculated with ≈ 1 × 10^9^pre-culture yeast cells. A batch phase continued for ≈40h until the cells had reached stationary phase, indicated by a sharp decrease in respiratory activity. Then continuous culture, *i.e*., feeding with fresh medium, was initiated (at 44.5 h in Figure 2).

#### Culture Control & Monitoring

Continuous culture was performed in a custom-built bioreactor. The culture vessel was a jar fermentor(Eyela, Japan) with a total volume of 2.667 L. Culture volume was measured using a balance (SB16001, MettlerToledo, Japan), and continuous dilution with fresh medium was performed using a peristaltic pump (AC2110, ATTA, Japan) with a six roller planetary design which minimizes pulsing during rotation (about 10rpm), and medium was pumped through 1 mm tubing (inner diameter; Masterflex, Cole Palmer, USA) and a 23 gauge steel needle. This ensured that the media was introduced in a stream of <20 μL droplets and just under a droplet per second at the operating dilution rate. Feed medium bottle weight was monitored by a balance (PMK-16, Mettler Toledo, Japan), set up to read from unstable environments and shielded from direct breezes. The culture was agitated at 750rpm and aerated at 0.150Lmin^-1^ by a mass flow controller (B.E. Marubishi, Japan). Dissolved oxygen was measured using an InPro 6800 sensor and pH with an InPro 3030 (both: Mettler Toledo, Japan). Culture pH was maintained at 3.4 by the automatic addition of 2.5mol L^-1^ NaOH, and the weight of the NaOH bottle was monitored on a balance (PM400). Local control of agitation and pH was carried out by Labo controllers (B.E. Marubishi, Japan). The reactor pressure was monitored by a manometer (DM-760, Comfix, Japan) installed on a split outlet flow stream. The culture temperature was controlled at 30°C by an external sensor connected to a circulating water bath (F25-ME, Julabo, Japan). Partial pressure of oxygen and carbon dioxide in the off-gas were measured by an Enoki-III gas analyzer (Figaro engineering, Japan). The partial pressure of hydrogen sulfide in the off-gas was measured using an electrode based gas monitor (HSC-1050HL, GASTEC, Japan). Instruments were calibrated as per manufacturer’s instruction.

#### Reactor Data Acquisition and Calculation of Metabolic Rates

Data were acquired *via* the in-house `FERMtastic` software at 0.1 Hz. Metabolic rates were calculated as described previously (***von Meyenburg, 1969a***; ***Heinzle, 1987***; ***Verduyn et al., 1991***; ***Marison et al., 1998***; ***Murray et al., 2007***) from the online recorded data. Details and all equations are provided in Appendix A of the supporting information. All data were processed in the script `samplingSeq_2019.R` of the `yeastSeq2016` git repository. All calculated rates are provided in Dataset S1.

### RNA Sequencing & Read Mapping

#### Sampling, RNA Extraction & Sequencing Library Generation

Total RNA was extracted as previously described (***Sasidharan et al., 2012***) from 24 samples taken every 4 min, covering ca. 2.5 cycles of the respiratory oscillation.

Culture samples were immediately quenched in ethanol and disrupted using acid-washed zirconia/silica beads (0.5 mm; Tomy Seiko Co., Ltd., Japan) with sodium acetate buffer (250 μL; sodium acetate 300mM, Na2-EDTA 10mM, pH 4.5–5.0) and one volume of TE-saturated phenol (Nacalai Tesque) equilibrated with sodium acetate buffer (250 μL).

The samples were then centrifuged (12000g, 15min, 4°C) and the aqueous phase transferred to fresh 1.5mL microcentrifuge tubes. Back-extraction was performed by adding sodium acetate buffer (125 μL)to the bead-beat tubes, vortex (10 s), centrifuging (12 000 g, 15min, 4 °C) and adding the aqueous phase to the first aqueous phase. 2.5 volumes ice-cold 99.5 %ethanol were added to the aqueous phase and RNA/DNA precipitated at −20 °C overnight. The samples were then centrifuged (12000g, 30min, 4 °C), the supernatant removed by aspiration, and pellets washed 3× in 500 μL 70%ethanol and air-dried (10min, room temperature). DNAwas removed (RNase-Free DNase Set; Qiagen, Japan) and RNA recovered by column purification (QIAquick PCR Purification Kit; Qiagen, Japan) in 50 μL UltraPure water, and stored at-80 °C prior to analysis. Total RNA had an RNA integrity number >7 and 260nm:230nm and 260nm:230nm ratios >2.14. All cDNA libraries were then generated and sequenced by the Beijing Genome Institute (BGI), China. Strand specific cDNA libraries were created using the “dUTP method” (***Parkhomchuk et al., 2009***; ***Levin et al., 2010***) and sequencing was carried out on an Illumina 1G sequencer.

#### RNAseq Read Mapping

RNAseq reads were mapped against the yeast reference genome (strain S288C, release R64-1-1) using `segemehl` (version 0.1.4) (***Hoffmann et al., 2014***) with default parameters and spliced read mapping enabled. Initially unmatched reads were mapped again using the remapping tool from the `segemehl` package and the resulting files were merged. Coverage (read-counts per nucleotide) was normalized for total library size to reads-per-million (RPM) and RPM values were stored in a bedgraph file for further analysis.

### RNAseq Time Series Analysis

#### Analysis Strategy and R Code

All analyses were performed with bash and R. The full analysis pipeline is available in a git repository at https://gitlab.com/raim/yeastSeq2016. Analysis and plotting tools developed for this work are available in an git repository with scripts and an R package available at https://github.com/raim/segmenTools. RNAseq segmentation was performed with the `segmenTier` R package (***Machné et al., 2017***), available at https://cran.r-project.org/package=segmenTier. Scripts for genome-wide data collections and mapping to the yeast S288C reference genome (release R64-1-1) as well as the genomeBrowser plots are available at the git repository https://gitlab.com/raim/genomeBrowser. The collection of oscillation period data and the scripts for the PWM model analysis are available at the https://gitlab.com/raim/ChemostatData repository, generated originally for ***Machné*** (***2017***).

#### Additional Data Sources

Genome annotations including Gene Ontology (GO) terms were taken directly from the gff genome file from the *Saccharomyces* genome database (SGD, release R64-1-1, 2011-02-08, same as for RNAseq mapping). Published transcript data sets (XUT, SUT, etc.) were also obtained from SGD for the same genome release. Protein complex annotation CYC2008 (***Pu et al., 2009***) was downloaded from http://wodaklab.org/cyc2008/resources/CYC2008_complex.tab on 2019-06-04. All other data were obtained from the supporting material of publications: half-live data for mRNAs and proteins from ***Geisberg et al.*** (***2014***) and ***Christiano et al.*** (***2014***); ribosome density data from ***Arava et al.*** (***2003***); the consensus clustering of periodically expressed transcripts from ***Machné and Murray*** (***2012***); UGRR expression data and slopes from ***Slavov and Botstein*** (***2011***); protein abundance data from ***Paulo et al.*** (***2016***), where growth rate data was sent in personal communication; and functional gene groups from (***Metzl-Raz et al., 2017***).

#### Discrete Fourier Transform

A time series of ***N*** measurements *x* = {*x*_0_,…, *x*_*N*–1_}, taken at equally spaced time points {*t*_0_,…, *t*_*N*-1_}, can be transformed to frequency-space by the Discrete Fourier Transform (DFT):

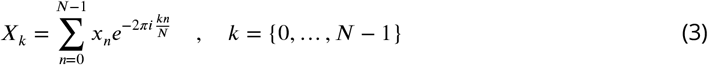

where ***X**_k_* is a vector of complex numbers representing the decomposition of the original time series into a constant (mean) component (at *k*= 0) and a series of harmonic oscillations around this mean with periods ***P**_k_*, amplitudes ***A**_k_* and phase angles *ϕ_k_*:

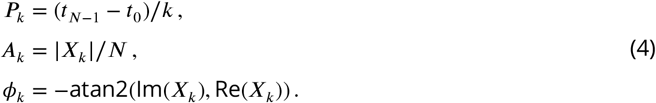

All DFT were performed with R’s `fft` function.

For DFT-based clustering and segmentation analysis, it proved useful to scale DFT components by the mean amplitude of all other components *k* > 0:

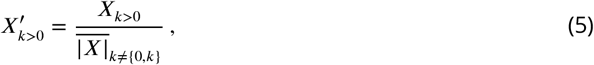

and the constant component (*k* = 0) by the arcsinh transformation:

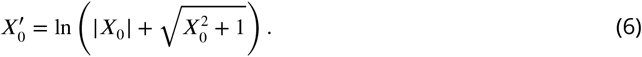

For analysis of read-count data *x_n_* were the raw read-counts, for analysis of segments *x_n_* were the mean of all read-counts of the segment.

The index *k* corresponds to the number of full cycles with period ***P**_k_* in the time series. Only the first 19 time points, covering two full cycles of the oscillation were used for the calculation of phases and p-values, such that *k*= 2 reflects the main oscillation. For all plots, phases were shifted such that *ϕ*_2_= 0 corresponds to the transition from LOC to HOC.

#### Oscillation p-Values

For calculation of oscillation p-values *p*_DFT_ on read-count level the time series were permutated ***N*** = 10,000 times, and random amplitude *Ã*_2_ calculated. The p-value was estimated as the fraction of permutations for which the random amplitude was larger than the observed amplitude ***A***_2_ (eqn.4). This analysis was performed with the script `genomeOscillation.R` from the `segmenTools` git repository. Oscillation p-values *p*_rain_ on segment level were calculated with the R package `rain` (***Thaben and Westermark, 2014***) using period *P* = 0.65 h and time step *δt* = 4 min. This analysis was done with the script `segmentDynamics.R` from the `segmenTools` git repository.

#### Segmentation of RNAseq Read-Counts & Segment Classification

The data were pre-segmented into expressed and weakly expressed chromosomal domains by a previously described heuristic (***Machné et al., 2017***) with a minor correction that splits pre-segments at chromosome ends. Pre-segmentation was done with the script `presegment.R` from the `segmenTools` script collection; Figure S4 provides pre-segment length distributions and run parameters. Pre-segments were then individually split into non-overlapping segments with coherent temporal expression profiles by the `segmenTier` algorithm, using optimized parameters from our previous study (***Machné et al., 2017***). Shortly, the arcsinh-transformed read-count data was Fourier-transformed (Eq.3); the first component (*k* = 0), reflecting the mean expression level, was `arcsinh`-transformed (Eq.6); and all other (*k* > 0) components were amplitude-scaled (Eq. 5). The real and imaginary parts of the scaled DFT components 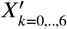 were then clustered into 12 groups with R’s implementation of `k-means` (using the `Hartigan-Wong` method or if that failed, the `MacQueen` method). This clustering then provided the anchors for the similarity-based segmentation by the `segmenTier`, where we used the `icor` scoring function with exponent *ϵ* = 2, length penalty ***M*** = 150, nuissance cluster penalty ***M***_0_ = 100, and nuissance cluster exponent *ν* = 3. This combination of parameters is achieved by arguments `–trafo “ash”–dc.trafo “ash”–dft.range 1,2,3,4,5,6,7 –K 12 –Mn 100 –scores “icor”–scales 2 –M 150 –nui.cr` 3 to the `runSegmentier.R` script in the `segmenTools/scripts` collection. All segments are provided in Dataset S2.

The resulting segments were then filtered and classified by their oscillation p-values (*p*_rain_, see above) and their overlaps with transcribed features annotated in the reference genome (release R64-1-1), using `segmentOverlaps.R` and `segmentAnnotation.R` in the `segmenTools/scripts` collection. Overlaps were quantified as the Jaccard index, 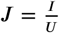, where ***I*** is the intersect, the number of overlapping nucleotides, and *U* the union, the number of nucleotides covered by both, the segment and the annotated feature. Table S2 provides details on filtering and the resulting sizes (numbers) of analyzed segment sets. Figure S5 provides the full data structure which guided these threshold choices.

#### Segment Clustering

The means of read-counts covered by a segment were taken as segment time series. Periodic expression was analyzed by permutation analysis and DFT and by the R package `rain`. 11,248 segments with *p*_rain_ < 0.85 were chosen for further analysis (Fig. S5E-F). The DFT of the segment time series was amplitude-scaled (Eq.5, Fig. S6A) and the first (constant) component (*k* = 0) was `arcsinh`-transformed (Eq.6). Real and imaginary parts of the scaled DFT components 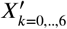 were then clustered with the `flowClust` algorithm (***Lo et al., 2009***) for cluster numbers *K* = 2,…, 16. The clustering with the maximal Bayesian Information Criterion, as reported by `flowClust` (Fig. S6B), was selected for further analysis. Clustering was performed by `clusterTimeseries2` function of `segmenTools` *via* the `segmentDynamics.R` script). The resulting clustering was sorted, re-labeled and colored automatically based on the means of their segments’ expression phases (Eq.4). The clustering was further sub-divided into high-amplitude clusters enriched for coding genes and low-amplitude clusters (compare Fig. S7 and S8).

#### Relative Protein Amplitudes

3,189 segments overlapping with a coding region with Jaccard index ***J*** > 0.5 and with protein halflive (*τ*_1/2_) annotation in (***Christiano et al., 2014***) were considered. Proteins with half-life annotation “>=100” were treated as *τ*_1/2_ = ∞. The relative mRNA amplitudes were calculated from the DFT *X_k_* (Eq. 3–4) of the first 19 time points (2 full cycles) of the RNAseq read count time series as *A*_R_ = ***X***_2_/***X***_0_, *i.e*., the ratio of the amplitudes of the *2^nd^* component ***X***_2_ (2 cycles) over the 0^*th*^ component ***X***_0_ (corresponds to the mean overall time points) of the DFT. Relative protein amplitudes ***A***_P_ were then calculated with the analytical solution to an ordinary differential equation of rhythmic production, after equation S8 of (***Lück et al., 2014***), as

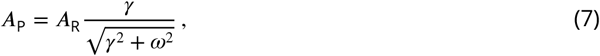

with angular frequency 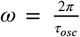 and *τ_osc_* = 0.67 h; the total protein degradation rate *γ* = *δ_p_* + *μ*, where the actual protein degradation rates *δ_p_* were taken from (***Christiano et al., 2014***); and the growth rate equals the chemostat dilution rate *μ* = *ϕ* = 0.089h^-1^. In this model, the relative amplitude ***A***_R_ is assumed to directly reflect periodic production, *i.e*., translational activity. Total amounts or translation rates are not required, and only a relative amplitude of protein amount can be calculated. Predicted protein amplitudes are provided in Dataset S3.

#### Transcript Abundance Peak Width Analysis

For each high-amplitude segment (2,505 segments with *p*_rain_ < 0.0001) the time series was interpolated to 1° resolution (0.105 min), and the oscillation phase *ϕ*_2_ (Eq.4) was used as anchors to scan for times spent above the temporal median ***X*** during the first and the second full cycle in the data set (horizontal arrows in Fig.3A). These times were recorded as the peak widths ***W***_1_ and ***W***_2_. The peak width change is the difference **Δ*W*** = ***W***_2_ – ***W***_1_. Only segments with peak phases with ≥60°distance to the start or end of the timeseries and where the median expression was traversed twice within one cycle were considered, resulting in 2,357 segments with **Δ*W*** values. See Figure S16 for an example and all data. Peak widths of other transcriptome data sets (Fig. S10B) were calculated for the first full cycle of each experiment, simply as the time spent above the mean of transcript abundance over the first cycle. Data that were not sampled equispaced were interpolated at equispaced time points using the minimal time step of the original sampling.

### Cluster Enrichment Analyses

#### Cluster-Cluster Enrichment Tests

Categorical enrichments, *e.g*. coding gene co-expression cohorts *vs*. gene annotations, were analyzed by cumulative hypergeometric distribution tests (R’s `phyper`) using `segmenTools`’s `clusterCluster` function and the `clusterAnnotation` wrapper for GO and and protein complex analysis, which compares overlaps of each pair of two distinct classifications into multiple classes, and stores overlap counts and p-values (“enrichment tables”) for informative plots (see “Enrichment Profiles”).

In these tests, the complete set of ORF annotated in the reference genome was analyzed (urn size: 5,795). Of these, 4,489 ORF that overlapped with an segment (Tab. S2) with a Jaccard index ***J*** > 0.5 were assigned to this segment’s cluster, where non-clustered segments (*p*_rain_ ≥ 0.85) were assigned to cluster “0”, and all non-overlapping ORF (***J***_ORF,max_ < 0.5) assigned to the “n.a” cluster. For the analysis of protein complex analysis, all 5,524 ORF that overlapped with a segment with ***J***_ORF_ > 0, and the one with the maximal ***J***_ORF_ was used for cluster assignment. This relaxed assignment was used to comprehensively capture complex co-expression and differential expression.

#### Enrichment Profiles

The results of multi-class enrichment tests (segment overlaps or cluster-cluster categorical overlaps) were visualized as colored table plots, *e.g*. Figure4A), using `segmenTools`’function `plotOverlaps`. The total counts of overlapping pairs are plotted as text, where the text color is selected based on a p-value cutoff *p*_txt_(as indicated). The background color gray level of each field scales with log_2_(*p*), such that fields with a minimal p-value *p*_min_(as indicated) are black.

For intuitively informative plots the enrichment tables were sorted. Table rows were sorted along the other dimension (table columns) such that all categories enriched above a certain threshold *p*_sort_ in the first column cluster are moved to the top, and, within, sorted by increasing p-values. Next, the same sorting is applied to all remaining row clusters for the second column cluster, and so on until the last column cluster. Remaining row clusters are either plotted unsorted below a red line or removed. This is especially useful to visualize enrichment of functional categories along the temporal program of co-expression cohorts, *e.g*., Figure 4A and D. This sorting is implemented in `segmenTools`’function `sortOverlaps`.

## Author Contributions

DBM designed and performed the experiment, SHB performed the RNAseq read mapping, RM and PFS developed the segmentation algorithm. RM and IMA analyzed the time series data. All authors contributed to data interpretation and to writing of the manuscript.

## Acknowledgments

We thank Sarah Lück, Oliver Ebenhöh, Wolfram Liebermeister, Oliver Bodeit, Ovidiu Popa, Chilperic Armel Foko Kuate and St. Elmo Wilken for inspiring discussions of the data and critical review of the manuscript. We are grateful to Martin Senz from the Berliner Institutf. Gärungsgewerbe und Biotechnologie, a successor of Paul Lindner, for help with clarifying the origin of the IFO 0233 strain.

