## Appendix A, B and Supporting Figures for "Pulse-Width Modulation of Gene Expression in Budding Yeast"

### RNAseq Time Series in IFO 0233
### Supporting Methods, Tables & Figures

Rainer Machné, Douglas B. Murray, Stephan Bernhart, Ilka Axmann, Peter F. Stadler

September 2, 2022

#### Appendix A  Calculation of Metabolic Rates

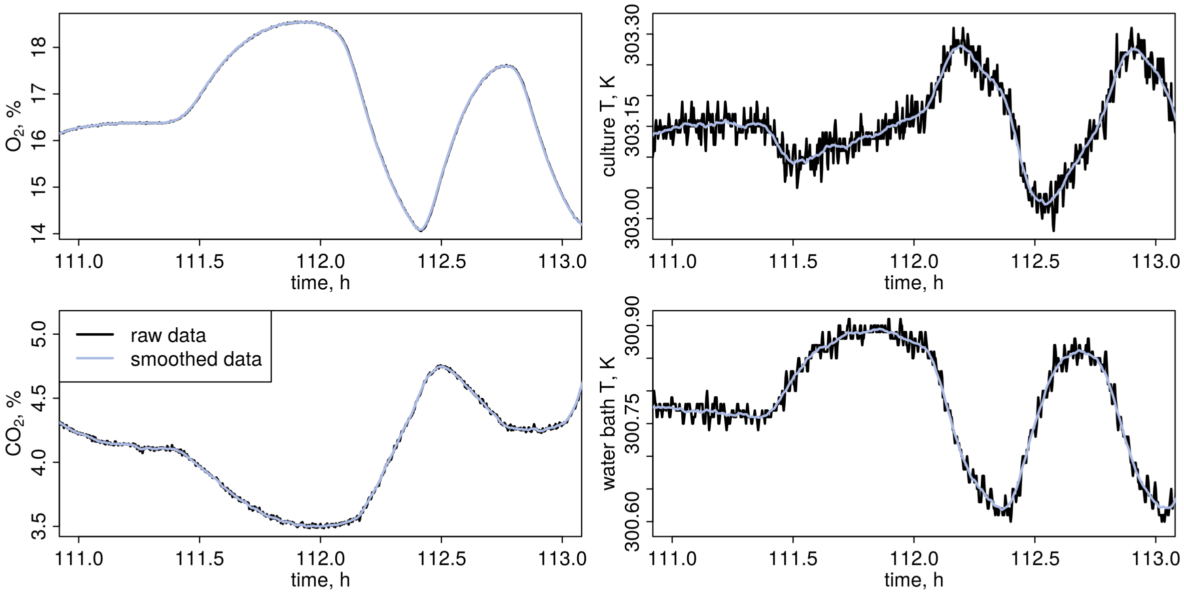


Figure A1: **Raw Data Smoothing.** Offgas partial pressures (left: $O_2$ and $CO_2$) were smoothed by a moving average over 10 measurements (90 s) and culture and water bath temperature (right panels) over 30 measurements (290 s).

**Data Pre-Processing.** Before all calculations some data were mildly smoothed (Fig. A1) by taking the means of moving windows of 10 measurements (90 s), for all offgas partial pressures ($p_{\text{out}}(t)$), and of 30 measurements (290 s) for the culture and water bath temperatures ($T(t)$ and $T_{\text{b}}(t)$). This was required to avoid noise in manual differentiation in offgas correction and in estimation of heat dissipation.

**Culture Dilution Rate $\phi$** The decreasing weights of the feed medium $W_{\text{feed}}(t)$ and NaOH (pH correction) bottles $W_{\text{NaOH}}(t)$ were monitored, and the total culture dilution rate $\phi$ was calculated (Fig. A2) from the change of the total weight, the densities $\rho_{\text{NaOH}}$, $\rho_{\text{H}_2\text{O}}$, and the set culture volume $V_\ell$:

$$\begin{aligned} V(t) &= \frac{W_{\text{NaOH}}(t)}{\rho_{\text{NaOH}}} + \frac{W_{\text{feed}}(t)}{\rho_{\text{water}}} \\ \phi_{\text{raw}}(t) &= -\frac{\mathrm{d}V(t)}{\mathrm{d}t}\frac{1}{V_\ell} \end{aligned} \tag{S1}$$

where $\mathrm{d}V(t)$ and $\mathrm{d}t$ were calculated as the means in moving windows of 200 s (21 measurements). The contribution of pH correction was oscillating, but the amplitude neglibible. The data showed constant feed and only the mean $\phi = \overline{\phi}_{\text{raw}} = 0.089\,\text{h}^{-1}$ was considered in all subsequent calculations.

**Period Determination from Dissolved $O_2$** Oscillation periods $\tau_{\text{osc}}(t)$ were calculated from the dissolved $O_2$ measurement using the `WaveletComp` R package [Roesch and Schmidbauer, 2018] with parameters `loess.span`=0 and `dj`=0.05.

**Calculation of $q_{O_2}$, $q_{CO_2}$ and RQ** The culture temperature $T(t)$, reactor overpressure $P_r(t)$, and partial pressures in the offgas $p_{\text{out}}(t)$ of gases $O_2$, $CO_2$ and $H_2S$ were recorded, and metabolic rates calculated after Heinzle [1987] but with correction for the delay of offgas measurements [Murray et al., 2007] in four steps as follows:

Table S1: Physical & Culture Parameters used in calculations.

| parameter | symbol | value ± S.D. | note or reference |
|---|---|---|---|
| PHYSICAL | | | |
| universal gas constant | $R$ | $8.314\,472\,\mathrm{J\,K^{-1}\,mol^{-1}}$ | |
| standard temperature | $T^{\ominus}$ | $298.15\,\mathrm{K}$ | |
| atmospheric pressure | $P_0$ | $101\,325\,\mathrm{Pa}$ | |
| atmospheric partial pressure, $O_2$ | $p_{\mathrm{in,O_2}}$ | 20.95% | |
| atmospheric partial pressure, $CO_2$ | $p_{\mathrm{in,CO_2}}$ | 383 ppm | |
| atmospheric partial pressure, $H_2S$ | $p_{\mathrm{in,H_2S}}$ | 0 | |
| Henry coefficient, $H_2S$ | $H^{\ominus}$ | $1\times10^{-3}\,\mathrm{mol\,m^{-3}\,Pa^{-1}}$ | Sander [2015] |
| -"- temperature factor | d$H$ | $2100\,\mathrm{K}$ | Sander [2015] |
| density of 10% NaOH at 27° C | $\rho_{\mathrm{NaOH}}$ | $1.105\,61\,\mathrm{kg\,l^{-1}}$ | `http://www.handymath.com/cgi-bin/naohtble3.cgi` |
| density of feed medium | $\rho_{\mathrm{medium}}$ | $1\,\mathrm{kg\,l^{-1}}$ | |
| REACTOR, static | | | |
| culture volume | $V_\ell$ | $0.635\,\mathrm{L}$ | |
| time delay, offgas measurement | $\tau$ | $13\,\mathrm{min}$ | |
| reactor inner surface area | $A$ | $0.132\,217\,9\,\mathrm{m^2}$ | |
| reactor wall thickness | $\Delta x$ | $3.2\,\mathrm{mm}$ | |
| Pyrex glass heat transfer coefficient | $k$ | $1.154\,784\,\mathrm{W\,m^{-1}\,K^{-1}}$ | Carwile and Hoge [1966] |
| REACTOR, calculated | | | |
| dilution rate | $\phi$ | $0.089\pm0.003\,\mathrm{h^{-1}}$ | |
| aeration rate | $\phi_g$ | $0.150\pm0.001\,\mathrm{L\,min^{-1}}$ | |
| reactor pressure | $P$ | $101\,707\pm102\,\mathrm{Pa}$ | |
| reactor temperature | $T$ | $303.15\pm0.04\,\mathrm{K}$ | * |
| water bath temperature | $T_\mathrm{b}$ | $300.79\pm0.05\,\mathrm{K}$ | * |
| culture pH | pH | 3.4±0.1 | * |
| REACTOR, from previous experiments | | | |
| biomass, dry cell weight | $X$ | $8\,\mathrm{g_{DCW}/L}$ | Sasidharan et al. [2012] |

*) these values varied with respiratory oscillation (see Text).

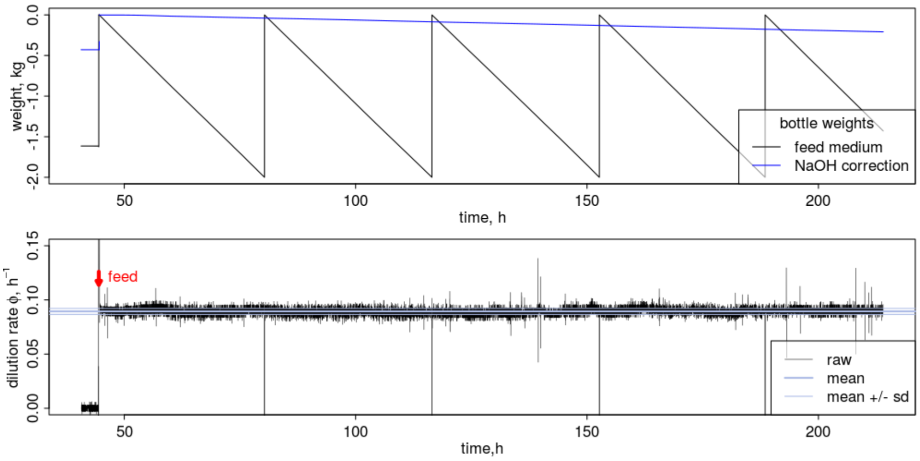


Figure A2: **Calculation of Dilution Rate $\phi$.** Upper panel: recorded weights of medium and NaOH correction bottles. The discrete changes stem from attachement of fresh medium bottles. Lower panel: the dilution rate $\phi$ (black lines) was calculated from the sum of both weights (see Methods). The noise is an artefact of manual differentiation. The global mean (after onset of feed, red arrow at 44.5 h) is the thicker solid blue line, and mean ± 1 standard deviation (sd) are the thinner blue lines.

Net total mass changes between ingas and offgas were corrected by assuming atmospheric partial pressures $p_{\mathrm{in}}$ and no net change in "inert" gases ($N_2$):

$$\begin{aligned} p_{\mathrm{in,inert}} &= 1-(p_{\mathrm{in,O_2}}+p_{\mathrm{in,CO_2}}+p_{\mathrm{in,H_2S}}) \\ p_{\mathrm{out,inert}}(t) &= 1-(p_{\mathrm{out,O_2}}(t)+p_{\mathrm{out,CO_2}}(t)+p_{\mathrm{out,H_2S}}(t)) \\ \hat{p}_{\mathrm{out}}(t) &= p_{\mathrm{out}}(t)\frac{p_{\text{in, inert}}}{p_{\text{out, inert}}(t)}\,. \end{aligned} \tag{S2}$$

The measurements were then corrected for a lag-time of $\tau = 13$ min between culture and measurement device, mostly determined by the head-space volume $V_h \approx 2$ L and aeration rate $\phi_g = 0.15\ \mathrm{L\,min^{-1}}$ as $\tau = V_h\phi_g$, and using a first-order transport equation:

$$p_{\mathrm{culture}}(t) = \hat{p}_{\mathrm{out}}(t) + \tau\frac{\mathrm{d}\hat{p}_{\mathrm{out}}(t)}{\mathrm{d}t}\,, \tag{S3}$$

where $\mathrm{d}\hat{p}_{\mathrm{out}}(t)$ and d$t$ were calculated as the means in moving windows of 200 seconds (21 measurements), resulting in a good match between $p_{\text{culture, O}_2}(t)$ and the measured dissolved $O_2$ concentration in the culture (Fig. A3).

The total metabolic rates $r(t)$ of the culture were then calculated in $\mathrm{mol\,h^{-1}}$ assuming the ideal gas law:

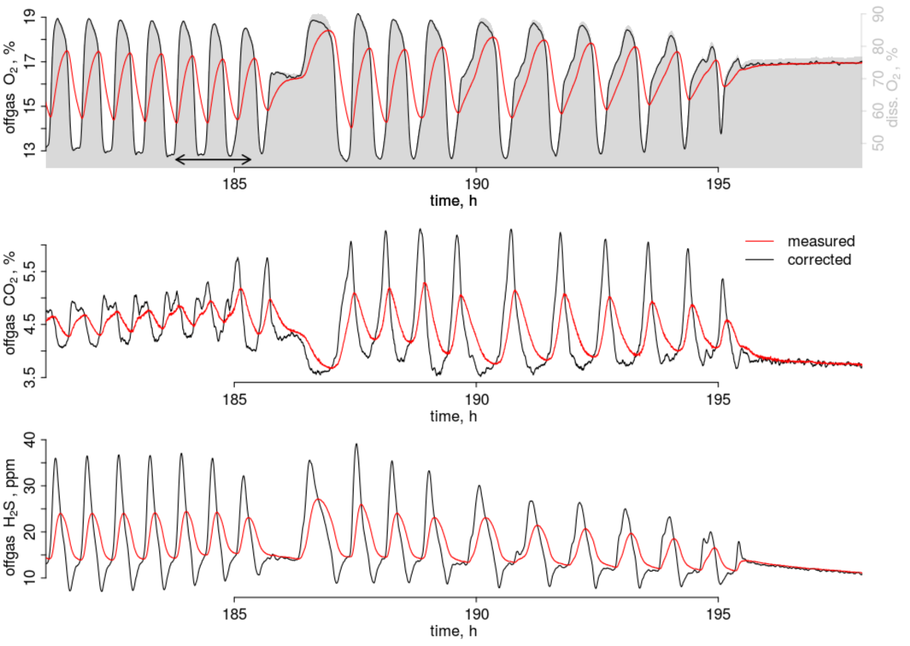


Figure A3: **Offgas Delay Correction.** Measured (red lines: $\hat{p}_{\text{out}}(t)$) and corrected (black lines: $p_{\text{culture}}(t)$) offgas concentrations (see Methods). The gray background in the top panel is the dissolved $O_2$ concentration (right y-axis), and comparison to corrected offgas data indicates a good match.

$$
\begin{aligned}
\Delta p(t) =& p_{\text{in}} - p_{\text{culture}}(t) \\
P(t) =& P_r(t) + P_0 \\
r(t) =& \phi_g \frac{\Delta p(t) P(t)}{RT(t)} \,,
\end{aligned}
\tag{S4}
$$

with the universal gas constant $R$, atmospheric pressure $P_0$, the set aeration rate of $\phi_g$ , and the measured total pressure $P(t)$ and culture temperature $T(t)$.

Specific metabolic rates were calculated by normalizing to the total biomass in the culture:

$$
q(t) = \frac{r(t)}{XV_\ell} \,, \tag{S5}
$$

with biomass concentration $X$ and culture volume $V_\ell$. The culture pH was maintained at 3.4 and thus $CO_2$/bicarbonate buffering in the culture medium is negligible for calculation of $q_{CO_2}$ [Heinzle, 1987].

The respiratory quotient is the ratio of $CO_2$ and $O_2$ turnover rates (excretion and uptake, respectively):

$$
\text{RQ}(t) = \frac{q_{CO_2}(t)}{-q_{O_2}(t)} \,. \tag{S6}
$$

The temporal mean over several cycles ($\overline{RQ}$ in Figure 1A are the means of moving windows over 3500 measurements (9.7 h of experiment time).

**$H_2S$ Liquid Concentration** $H_2S$ had been explored for its potential activity as a synchronizer of the system [Sohn and Kuriyama, 2001, Murray et al., 2003] due to its inhibitory activity on respiratory complex IV, *e.g.*, with reported $K_i = 0.2\,\mu\text{M}$ [Szabo et al., 2014]. Thus, we calculated the liquid concentration $C$ in the culture from the corrected offgas partial pressure $p_{\text{culture}}(t)$, total reactor pressure $P(t)$ and culture temperature $T(t)$ *via* Henry's law:

$$
C(t) = p_{\text{culture}}(t) P(t) H^{\ominus} e^{\text{d}H(1/T(t) - 1/T^{\ominus})} \,, \tag{S7}
$$

with Henry coefficient $H^{\ominus}$ and temperature factor d$H$ from Sander [2015] (Table S1).

**Stoichiometric ATP Production Rates $q_{\text{ATP,ox}}$ & $q_{\text{ATP,ferm}}$** ATP production rates of glucose-grown cultures can be estimated from the rates $q_{O_2}$ and $q_{CO_2}$ [von Meyenburg, 1969a, Verduyn et al., 1991], using the known $CO_2$ and $O_2$ stoichiometries of glucose fermentation and oxidation, and experimentally estimated P/O ratios $P_{\text{O}} = \frac{\text{ATP}}{\text{O}}$, mol ATP produced per half mol $O_2$.

Omitting inorganic phosphates and protons for clarity, the overall stoichiometry of complete oxidation of glucose $C_6H_{12}O_6$ in glycolysis, tricarboxylix acid cycle (TCA) and respiration is:

$$
C_6H_{12}O_6 + 6\ O_2 + 2\ \text{ADP} + 2\ \text{GDP} + 12{*}P_{\text{O}}\ \text{ADP} \rightarrow 6\ CO_2 + 6\ H_2O + 2\ \text{ATP} + 2\ \text{GTP} + 12{*}P_{\text{O}}\ \text{ATP} \,. \tag{S8}
$$

Per mol consumed $O_2$, metabolism releases 1 mol $CO_2$, and a respiratory quotient $RQ_{glc} = 1$ indicates purely respiratory use of glucose. Counting GTP as ATP we can summarize the rate of ATP production from complete oxidation of glucose as:

$$q_{\text{ATP,ox,glc}} = 2\left(P_{\text{O}} + \frac{1}{3}\right) q_{\text{O}_2} \,, \tag{S9}$$

and with a compromise $P_{\text{O}}$=1 (1.1 in [von Meyenburg, 1969a], 0.95 in [Verduyn et al., 1991] we get a simple scaling $q_{\text{ATP,ox}} \approx 2.667 \times q_{\text{O}_2}$.

Additional fermentation increases RQ. The overall reaction of glucose fermentation to ethanol ($C_2H_6O$), again omitting phosphates and protons for clarity, is:

$$C_6H_{12}O_6 + 2\ \text{ADP} \rightarrow 2\ C_2H_6O + 2\ CO_2 + 2\ \text{ATP}\,, \tag{S10}$$

*i.e.* 1 mol ATP per mol $CO_2$. Assuming that surplus $q_{\text{CO}_2}$ completely stems from additional fermentation, we can subtract the respiratory from the total rate, to estimate the additional ATP production in fermentation:

$$q_{\text{ATP,ferm,glc}} = q_{\text{CO}_2} - q_{\text{O}_2} \,. \tag{S11}$$

**Fermentation & Ethanol Oxidation.** During HOC, the RQ < 1 indicates re-uptake and oxidation of the ethanol produced in the preceding LOC phase. The overall stoichiometry of ethanol ($C_2H_6O$) oxidation has a theoretical $RQ_{etoh} = \frac{2}{3}$:

$$C_2H_6O + 3\ O_2 + 6P_{\text{O}}\ \text{ADP} \rightarrow 2\ CO_2 + 3\ H_2O + 6P_{\text{O}}\ \text{ATP}\,, \tag{S12}$$

where 1 ATP lost during the initial reactions of ethanol assimilation is compensated by 1 GTP gained in TCA, and thus:

$$q_{\text{ATP,ox,etoh}} = 2P_{\text{O,etoh}} q_{\text{O}_2} \,. \tag{S13}$$

Ethanol has a higher degree of reduction (6) than glucose (4) [Roels, 1980], and accordingly Verduyn et al. [1991] reported a ≈50% higher P/O ratio on ethanol ($P_{\text{O,etoh}}$=1.5) than on glucose ($P_{\text{O}}$=0.95) medium. Assuming this $P_{\text{O,etoh}}$, we would arrive at a 10–15% higher overall ATP stoichiometry for ethanol re-uptake and oxidation ($q_{\text{ATP,etoh}} \approx 3 \times q_{\text{O}_2}$) than for glucose oxidation. Assuming complete fermentation and re-uptake and oxidation of the produced ethanol we obtain a lower estimate for glycolytic ATP generation of:

$$q_{\text{ATP,ferm,etoh}} = q_{\text{CO}_2} - \frac{3}{2} q_{\text{O}_2} \,. \tag{S14}$$

When assuming, as an extreme alternative scenario, full glycolytic fermentation of glucose to ethanol and ethanol as the sole substrate of respiration, the relative contribution of fermentation to total ATP generation would thus be even lower.

In Figure 1D, $q_{\text{ATPox}}$ and $q_{\text{ATPferm}}$ are the ranges between $q_{\text{ATPox,glc}}$ (lower) $q_{\text{ATPox,etoh}}$ (upper) and $q_{\text{ATPferm,glc}}$ (upper) $q_{\text{ATPferm,etoh}}$ (lower), respectively.

### Appendix B Mathematical Formulation of the PWM Model

#### B.1 The PWM Model of Gene Expression

We describe transcription of an mRNA species with concentration $R$ and its translation to a protein with concentration $P$ as:

$$\begin{aligned}\frac{dR}{dt} &= \tilde{k} - (\mu + \delta_r)R \\ \frac{dP}{dt} &= \ell n_B R - (\mu + \delta_p)P \ ,\end{aligned} \tag{S15}$$

where $\tilde{k}$ and $\ell$ are (effective) transcription and translation rates, $n_B$ is the number of ribosomes per RNA (ribosome density), $\delta$ are the respective degradation rates, and $\mu$ is the growth rate. In steady state the protein concentration will be:

$$P = \frac{\tilde{k}}{\mu + \delta_r} \frac{n_B \ell}{\mu + \delta_p} \ .$$

Now we assume a respiratory or metabolic cycle with alternating phases HOC and LOC. The relative lengths of the phases can vary, but they sum up to the total period $\tau_{\text{osc}}$ of the cycle:

$$\begin{aligned}\tau_{\text{osc}} &= \tau_{\text{hoc}} + \tau_{\text{loc}} \\ \phi_{\text{hoc}} &= \frac{\tau_{\text{hoc}}}{\tau_{\text{hoc}} + \tau_{\text{loc}}} \\ \phi_{loc} &= 1 - \phi_{\text{hoc}} \ ,\end{aligned}$$

where $\phi$ are the fractions of $\tau_{\text{osc}}$ spent in HOC and LOC phases.

Next, we assume that the RNA species $R_{hoc}$ and $R_{loc}$ are only transcribed during these phases at maximal transcription rate $k$, and not transcribed in the other phase, i.e. a simple step function of transcriptional activity. When averaging over multiple stable cycles, the effective transcription rates are:

$$\begin{aligned}\tilde{k}_{hoc} &= \phi_{\text{hoc}}\, k \\ \tilde{k}_{loc} &= \phi_{loc}\, k \ ,\end{aligned}$$

where $k$ is the real transcription rate. The protein abundances can then be calculated as:

$$\begin{aligned}P_{hoc} &= \frac{\phi_{\text{hoc}} k}{\mu + \delta_r} \frac{n_B \ell}{\mu + \delta_p} \\ P_{loc} &= \frac{\phi_{loc} k}{\mu + \delta_r} \frac{n_B \ell}{\mu + \delta_p} \ .\end{aligned} \tag{S16}$$

#### B.2 Constraints on Oscillation Period

Equation S16 can be re-arranged to emphasize a constraint between oscillation parameters and the life-cycle parameters and cellular abundance (concentration) of every protein that is transcribed in HOC (or LOC) phase and under the PWM assumption:

$$\frac{\tau_{\text{osc}}}{\tau_{\text{hoc}}} = \frac{k}{\mu + \delta_r} \frac{n_B \ell}{\mu + \delta_p} \frac{1}{P_{hoc}} \ . \tag{S17}$$

Ribosomal protein (RP) genes are always transcribed with the large HOC co-expression cohort. The yeast ribosome consists of $\approx 80$ distinct RP, most present only once per ribosome [Verschoor et al., 1998]. Free RP that are not part of a ribosome are quickly degraded [Warner, 1999, Beyer et al., 2004, von der Haar, 2008]. Thus, we can assume a stoichiometric equivalence, i.e. the cellular abundance of an RP equals the cellular abundance of each single RP [von der Haar, 2008]. From hereon we specifically consider such an RP gene, i.e., the parameters in S17 refer to a limiting or an average RP with concentration $P_{hoc} := P_{RP}$, and we replace its protein concentration by the ribosome concentration $P_{RP} = B$. We further summarize the two critical parameters, most likely to vary with growth rate [Waldron and Lacroute, 1975, Metzl-Raz et al., 2017], as:

$$\alpha = \frac{n_B}{B} \ , \tag{S18}$$

which can here be considered an autocatalytic efficiency of ribosomes: how many ribosomes are invested per RP mRNA to produce new ribosomes.

The time spent in HOC, $\tau_{\text{hoc}}$, changes little or sometimes even slightly increases with growth rate [Burnetti et al., 2016], while $\tau_{\text{loc}}$ proportionally decreases with increasing growth rate and decreasing period. If we assume a constant $\tau_{\text{hoc}}$ the total period can now be calculated as:

$$\tau_{\text{osc}} = \frac{\tau_{\text{hoc}} k\, \alpha \ell}{(\mu + \delta_r)(\mu + \delta_p)} \ . \tag{S19}$$

#### B.3 Species- and Condition-Specific Constraints

**Maximal Period.** Oscillations were also observed at the end of batch phase cultures, when cells are not dividing anymore [Mochan and Pye, 1973, Murray, 2004, Slavov et al., 2011], with overall consistent transcriptome dynamics [Slavov et al., 2011]. Burnetti et al. [2016] and Machné [2017] suggested the existence of a condition-specific and strain-specific period $\tau_{\text{osc}} = \tau_{\text{max}}$ at $\mu = 0$; Eq. S19 then yields:

$$\tau_{\text{max}} = \tau_{\text{hoc}} \frac{k \alpha \ell}{\delta_r \delta_p} \ . \tag{S20}$$

This also provides an interesting boundary case. When either of the two degradation rates is 0, $\delta_r \to 0$ or $\delta_p \to 0$, Eq. S20 implies

$$\lim_{\delta \to 0} \tau_{\text{max}} = \infty \ . \tag{S21}$$

**Minimal Period and Maximal HOC Fraction)** Machné [2017] and Burnetti et al. [2016] noted that minimal oscillation periods are observed near $\mu_f$, the strain-specific growth rate where the specific respiration rate is at maximum and additional fermentation sets in [Rieger et al., 1983]. This observation yields another parameter constraint, where

$$\tau_{\text{min}} k \alpha \ell = \tau_{\text{hoc}} (\mu_f + \delta_r)(\mu_f + \delta_p) \ . \tag{S22}$$

Combined with the observation that $\tau_{\text{hoc}}$ is approximately constant [von Meyenburg, 1969a, Strässle et al., 1989, Slavov and Botstein, 2011, Burnetti et al., 2016, O' Neill et al., 2020], Burnetti et al. [2016] suggested that at this point the cycle would consist only of a HOC phase, , i.e., $\tau_{\text{min}} = \tau_{\text{hoc}}$ and $\phi_{\text{hoc}} = 1$. Adopting this assumption we get:

$$k \alpha \ell = (\mu_f + \delta_r)(\mu_f + \delta_p) \ . \tag{S23}$$

Estimating RP-specific transcription and translation rates, and using measured RP mRNA and protein degradation rates (Tab. S5) and strain-specific $\mu_f$, we can obtain values of $\alpha$ between 2e-05 and 9e-05.

Combining both constraints (Eq. S20 and S23) we get a relation between the maximal period and the critical growth rate where fermentation sets in:

$$\frac{\tau_{\text{max}}}{\tau_{\text{hoc}}} = \frac{(\mu_f + \delta_r)(\mu_f + \delta_p)}{\delta_r \delta_p} \ , \tag{S24}$$

which constrains the degradation parameters. Thus, the model could predict species- or condition-specific period using only $\tau_{\text{max}}$, $\mu_f$ and one of the two degradation rates.

#### B.4 Additional Growth-Rate Dependency

We can relax the constancy assumption and extend the relation by an an additional growth-rate dependence on any parameter, here e.g. the ribosome concentration $B$:

$$\tau_{\text{osc}} = \frac{\tau_{\text{hoc}} k}{\mu + \delta_r} \frac{\ell}{\mu + \delta_p} \frac{n_B}{B_0 + \beta \mu} \ . \tag{S25}$$

This extended model yields steeper curves and better fits to periods observed close to the range of CDC synchrony. We can apply the same constraints as above, $\tau_{\text{max}}$ at $\mu = 0$ and $\tau_{\text{min}} = \tau_{\text{hoc}}$ at $\mu_f$, and e.g., can assume both a $\mu_f$ and a $\tau_{\text{max}}$ to calculate the linear function for ribosome numbers:

$$\begin{aligned} B_0 &= \frac{\tau_{\text{hoc}}}{\tau_{\text{max}}} \frac{k \ell n_B}{\delta_r \delta_p} \\ \beta &= \left( \frac{k \ell n_B}{(\mu_f + \delta_r)(\mu_f + \delta_p)} - B_0 \right) \frac{1}{\mu_f} \ . \end{aligned} \tag{S26}$$

#### B.5 A Pulse of Translation in HOC Phase

The cycle is accompanied by strong variation of the ATP/ADP ratio, from a minimum of $\approx$2 in early/mid LOC phase, to $\approx 7$ in HOC phase, in both IFO 0233 short period [Satroutdinov et al., 1992, Machné and Murray, 2012, Amariei et al., 2014] and CDC-coupled [von Meyenburg, 1969b, Müller, 2006, Xu et al., 2004] experimental systems. Our estimation of ATP production rates (Fig. 1 of the main manuscript) is consistent with this, indicating that the contribution by fermentation in the LOC phase is minimal.

At the transition to the LOC phase NAD(P)H fluorescence peaks [Münch et al., 1992, Murray et al., 1999]. In the short period oscillation (IFO 0233) first citric acid cycle and other intermediates, then amino acid concentrations peaked [Murray et al., 2007] at the transition to and in LOC phase, and decreased in late LOC phase. In long period systems, nitrogen uptake [von Meyenburg, 1969a,b] and amino acids pools that oscillated peaked in the HOC phase [Hans et al., 2003] and all decreased upon transition to the LOC phase. Consistent with this, protein translation rate (puromycin uptake) peaked in the HOC phase O' Neill et al. [2020].

TORC1 is a conserved regulator of global translational activity, and the TORC1 inhibitor rapamycin; upon pulse injection to the culture, HOC phase is immediately stopped and cells enter a long LOC phase in both short and long period systems [Murray et al., 2007, O' Neill et al., 2020]. The activating TORC1 phosphorylation (phospho-Rps6, Ser235/236) peaked in the HOC phase and became un-detectable in LOC phase [O' Neill et al., 2020]. During the LOC phase, translational activity and TORC1 phosphorylation were lower, and instead protein aggregation and proteasomal activity were observed [O' Neill et al., 2020].

These observations all indicate biosynthesis occurring in the HOC or at the HOC to LOC phase transition in both short and long period systems, ending with a pulse of protein translation. As a first approximation of this phenomenon as well as a first attempt to build in the obvious energetic constraints during the oscillation between a high-energy (HOC) phase and a low energy (LOC) phase with variable lengths, one may assume that also translation is confined to HOC phase, and independent of the transcription phase of the mRNA: $\ell_{\text{effective}} = \phi_{\text{hoc}}\ell$. This brings in the relative duration of phases a second time, and the equations above become, e.g.:

$$\begin{aligned}
\text{Eq. S16:}\quad P_{hoc} &= \phi_{\text{hoc}}^2 \frac{k}{\mu+\delta_r}\frac{n_B\ell}{\mu+\delta_p} \\
P_{loc} &= \phi_{\text{hoc}}\phi_{loc} \frac{k}{\mu+\delta_r}\frac{n_B\ell}{\mu+\delta_p} \\
\text{Eq. S19:}\quad \tau_{\text{osc}} &= \tau_{\text{hoc}}\sqrt{\frac{k\,\alpha\ell}{(\mu+\delta_r)(\mu+\delta_p)}}\,.
\end{aligned} \tag{S27}$$

#### B.6 Quantitative Discrepancies of Model Predictions

Qualitatively the PWM model, parameterized with generic gene length-dependent elongation rates and specific measured degradation rates, can predict the observed growth rate-dependent abundance levels of large gene groups quite well (see main manuscript, Fig. 4C–F, S21).

However, quantitatively our predictions (Fig. 4C–F, are off for some gene groups. For example the "translation" gene group occupies only 3% to 14% of the proteome in the PWM prediction, while this ranged from 19% to 38% in the proteome data evaluated by Metzl-Raz et al. [2017]. In contrast, proteins from the "stress" group are overestimated $\approx$2x by the PWM prediction. Total mRNA abundances are within the range of published estimates [Hereford and Rosbash, 1977, Zenklusen et al., 2008] at high growth rates, but 1.5x too high at low growth rates (Fig. S21B). Total protein abundances are within the estimated range [Milo, 2013] only at the highest growth rate, and 6x fold too high at slow growth (Fig. 4F, S21F). When comparing all gene-wise predictions with data, the main contribution for the good overall correlation stems from correct classification into HOC phase or LOC phase-specific genes, while correlations are low, absent or even negative within the two groups (Fig. S21C,G). This improves when focusing on the subsets of genes in a previously defined consensus clustering [Machné and Murray, 2012] of co-expression cohorts (Fig. S21D,H).

These errors help to pinpoint weaknesses of the model, and suggest directions for future extensions. The `Ribi` and `RP` cohorts (HOC) are characterized by large nucleosome-free regions (NFR) in their promoter, while the `S/C` (LOC) cohorts have occupied promoters and a less defined gene body nucleosome structure [Lee et al., 2007, Machné and Murray, 2012]. Total nucleosome occupancy (histone H3) at transcription start sites was lower in HOC and reached a maximum in LOC phase, anti-phase to histone acetylation (H3K9ac) and RNA polymerase II binding [Amariei et al., 2014]. Nucleosomes act as a barrier to RNA polymerase [Jimeno-González et al., 2015], and different chromatin structures are accompanied by different modes of transcription [Kristjuhan and Svejstrup, 2004, Zenklusen et al., 2008]. These observations all suggest that transcription of LOC phase-associated genes occurs at lower overall rates, which would rectify some of the observed discrepancies between model and data.

Similarly, translation rates could differ between the co-expression cohorts; e.g. RP genes are often assumed to have optimized codon usage, contributing to high RP abundances [Jansen et al., 2003]. However, we believe a more significant over-simplification of the model leads to the overestimation of protein abundances at low growth rates. Protein biosynthesis is globally regulated by nutrient availability via the TOR1 kinase. Its inhibitor rapamycin immediately stops HOC phase and induces a long phase of low respiratory activity in both IFO 0233 and CDC-coupled systems [Murray et al., 2007, O' Neill et al., 2020]. Cumulative evidence suggests (Appendix B.5) that a TOR1-mediated translational pulse occurs during HOC phase or at the transition from HOC to LOC phase, when the ATP/ADP ratio is high and then rapidly drops. As a first approximation of this phenomenon, we additionally assumed that translation of all transcripts occurs exclusively in HOC. Indeed, this assumption (Eq. S27) leads to more realistic total protein abundances at low growth rates (dashed lines in Fig. 4F, S21F,J). However, when using this assumption in period estimation, the obtained results are confined to too short periods (Fig. S21K). Future work based on this novel theoretical framework should explicitly account for energetic constraints (Fig. 1) on the protein synthesis capacity during the cycle.

#### B.7 Comparison to Related Models

**Empirical Oscillation Models.** For comparison to other models, we can express Eq. S19 as the oscillation frequency $f_{osc} = \frac{1}{\tau_{\text{osc}}}$, and this frequenciy is a quadratic function of growth and degradation rates:

$$f_{osc} = \frac{1}{\tau_{\text{hoc}}} \frac{(\mu + \delta_r)(\mu + \delta_p)}{k\alpha\ell} \; . \tag{S28}$$

This dependency of oscillation frequency on growth rate lies between the linear and exponential dependencies in purely empirical formulations by Burnetti et al. [2016] and Machné [2017], respectively.

Burnetti et al. [2016] found that they could fit their own data set with:

$$f_{osc} = f_{min}(\ln(2)\kappa\mu + 1) \; , \tag{S29}$$

which is a linear inverted form of an assumed Michaelis-Menten-like dependence of period on growth rate, with half-maximal parameter $\kappa$ and maximal period $\frac{1}{f_{min}}$ at long division times.

Machné [2017] collected data[1] from different strains from literature which did not match the linear function, and instead suggested:

$$f_{osc} = f_{min} e^{\mu\gamma} \; , \tag{S30}$$

where data fits yielded a $\gamma$ of 10 h–20 h, which was speculated to reflect measured protein half-lives in yeast. It was further noted that $f_{min}$ of different strains followed a period doubling relation with circadian $\frac{1}{f_{min}} \approx 24\,\text{h}$ for the strain with the longest periods.

**Cell Division Synchrony Models.** von Meyenburg [1969b] observed that different growth rates are mostly realized as different lengths of the G1 phase, while the length of the budding phase, comprising of S, M and G2 phases of the cell cycle, did not vary strongly. Parent cells have a shorter G1 phase and the smaller offspring cells grow in G1 phase until they reach the cell size appropriate for initiating their own budding cycle. Hartwell and Unger [1977] provided the definition of exponential growth for asymmetric cell division:

$$e^{-\mu\tau_{\text{O}}} + e^{-\mu\tau_{\text{P}}} = 1 \; , \tag{S31}$$

In continuous culture with high cell density, parent and offspring division cycles become synchronized such that the oscillation period corresponds to the doubling time of parent cells, while offspring cells spent one or two cycles with cell size growth before joining the cycle, or generalized:

$$\tau_{\text{osc}} = \frac{\tau_{\text{P}}}{i} = \frac{\tau_{\text{O}}}{j} \; , \tag{S32}$$

where $i \leq j$ are integers. A given combination of $i$ and $j$ is denoted a 'mode' `i:j`. Combining these and solving the polynomial $a^i + a^j - 1 = 0$, we can calculate $\tau_{\text{osc}}$ for any given mode.

The fraction $\psi$ of parent (mother) over offspring (daughter) cell numbers in mode `1:2` corresponds to the *golden ratio* [Bellgardt, 1994, Hjortso and Nielsen, 1995]: $\psi = \frac{n_{\text{p}}}{n_{\text{o}}} = e^{\tau_{\text{osc}}\mu} = \frac{1+\sqrt{5}}{2}$, and the oscillation period is lower than the culture doubling time:

$$\tau_{\text{osc}} = \frac{\ln(\psi)}{\mu} \leq \frac{\ln(2)}{\mu} . \tag{S33}$$

Duboc and von Stockar [2000] suggested a probabilistic extension of the mode `1:2` model:

$$\psi = \frac{1 + \sqrt{1 + 4p}}{2} , \tag{S34}$$

where $p$ is the probability of a cell to enter CDC in one metabolic cycle.

[1] https://gitlab.com/raim/ChemostatData

### Appendix C Supporting Figures

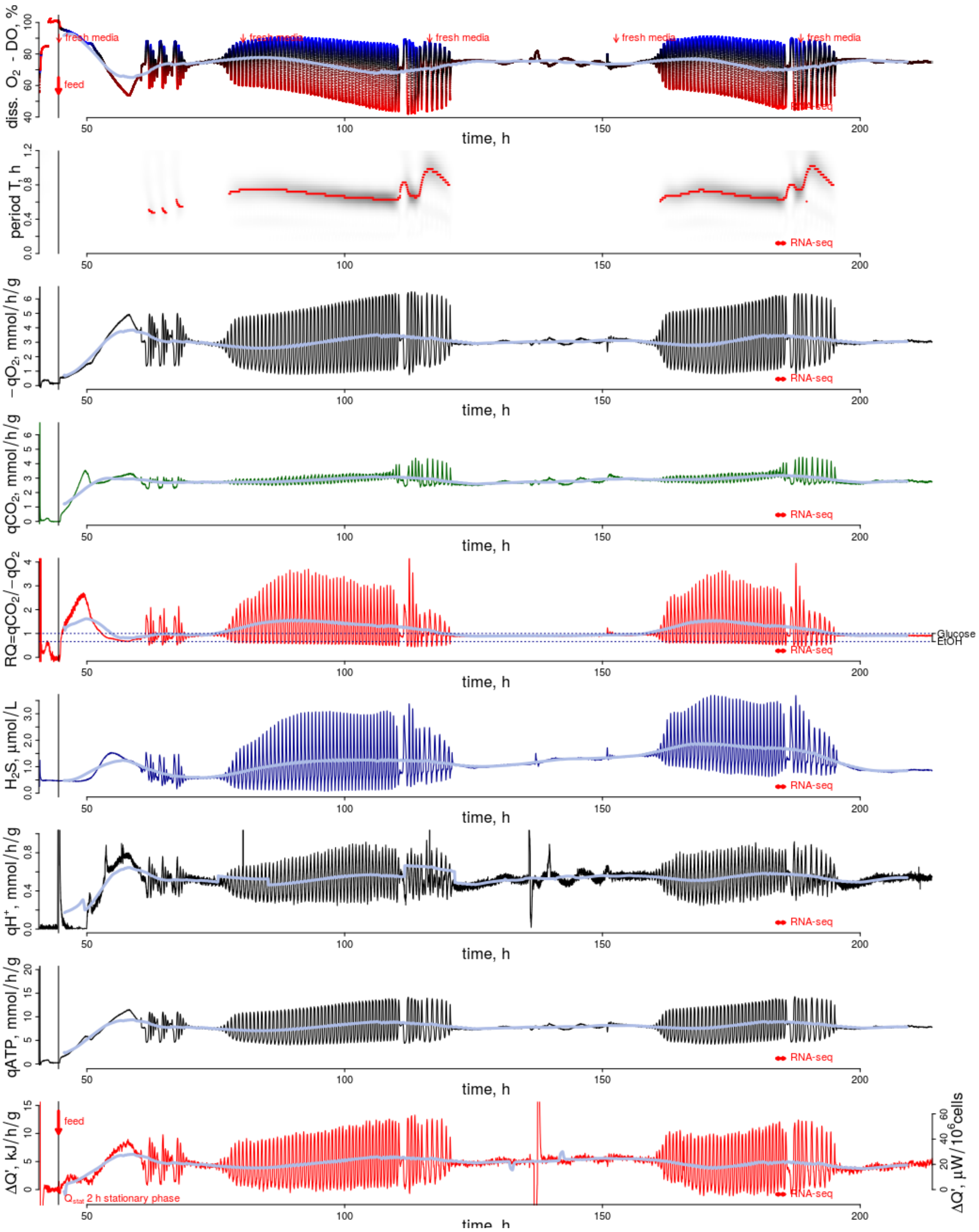


Figure S1: **Culture Dynamics - Time Series.** All data calculated from online measurements (see Methods) over the complete recorded time range, and normalized to 8 $g_{DCW}$/L [Sasidharan et al., 2012]. Red vertical arrows in the top panel indicate the start of continuous culture (feed) and changes of the media bottle (fresh media). The RNAseq experiment range is indicated in all panels by a horizontal arrow.

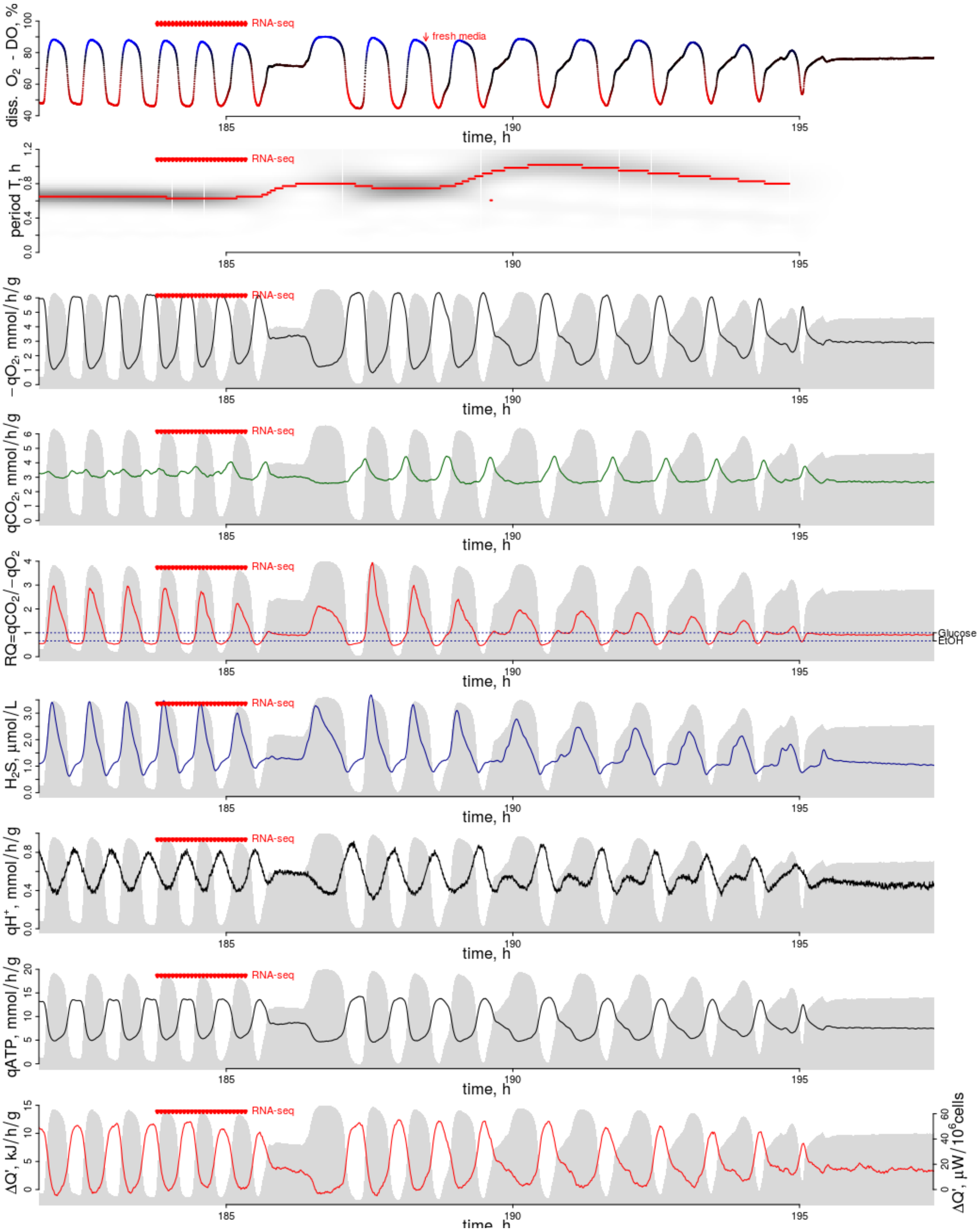


Figure S2: **Culture Dynamics - Time Series.** As Fig. S1 but zoomed in around RNAseq sampling. Each sample is indicated by a vertical arrow.

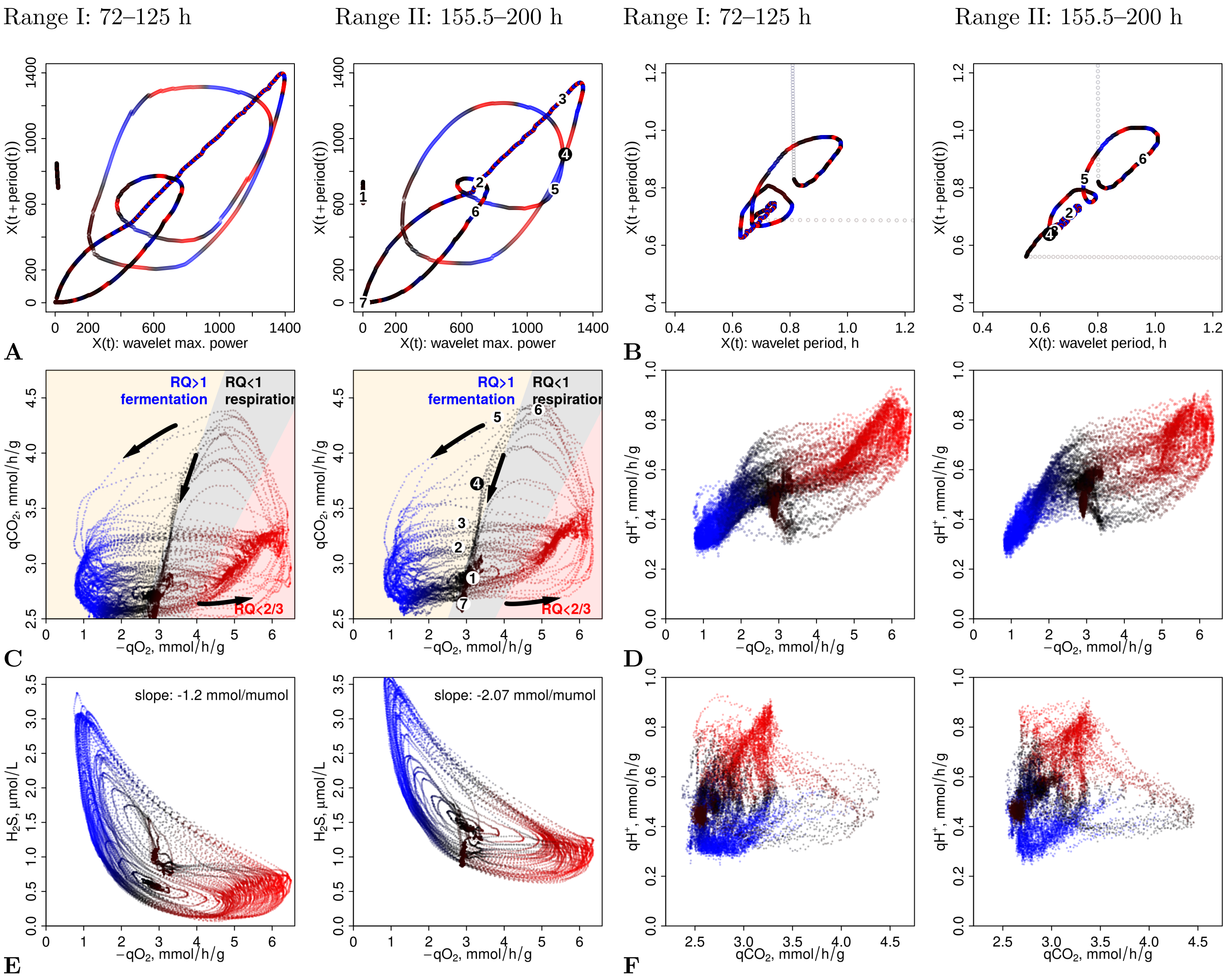


Figure S3: **Poincaré Maps & Phase Portraits** of various culture metabolic rates (Fig. S1 and S2) during the first (left in each panel) and the second (right) instant of the 30-50 h long transients. All data points are colored by the RQ value at the time (panel (B) of $q_{CO_2}$ *vs.* $q_{O_2}$ serves as a legend) and indicate phases of high $O_2$ consumption (red) and low $O_2$ consumption (blue). Background colors in the top panels indicate RQ ranges for respiratory (gray and red) and respiro-fermentative metabolism (blue). Bullet points (right panels only) map to their locations in Figure 1 of the main manuscript. Arrows indicate the time direction. All trajectories except $q_{CO_2}$ *vs.* $q_{H^+}$ (right bottom panel) move counter-clockwise. The phase advancement and delays at the bifurcation of system dynamics and the appearance of a novel phase at RQ $\approx$ 1, as outlined in the main manuscript, are indicated by trajectory points colored in black. **A & B:** Poincaré maps of the maximal power (A) of the wavelet transformed DO data and the period (B) with this power. **C–F:** Phase portraits, each comparing two metabolic rates. **C:** A $CO_2$ release pulse appears at transition to the RQ $\approx$ 1 phase. **D:** Phase-advanced proton export. **E:** Phase-delayed $H_2S$ pulse after the RQ $\approx$ 1 phase. **F:** Uncoupling of $q_{CO_2}$ and $q_{H^+}$.

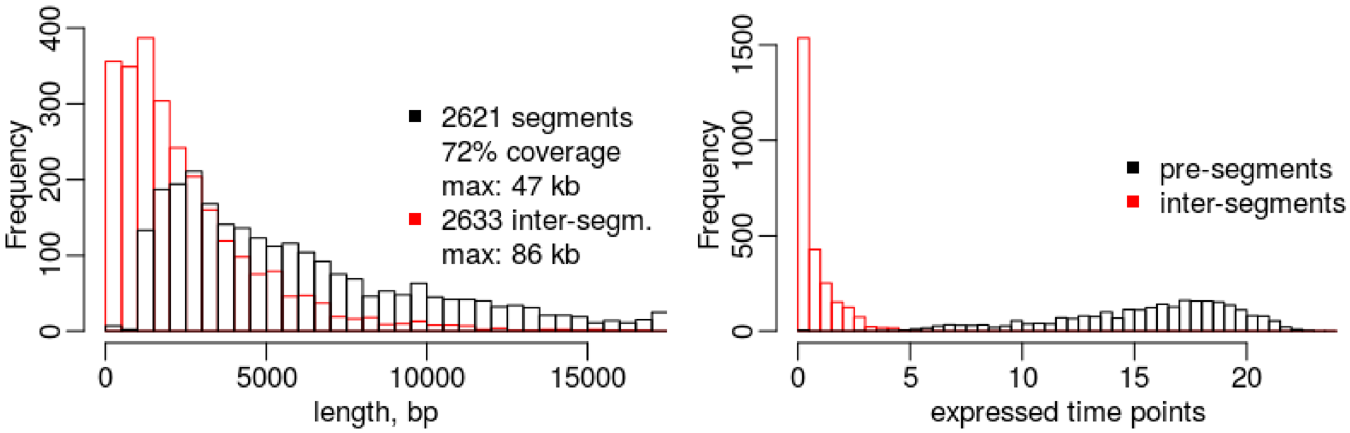


Figure S4: **Presegmentation of RNAseq Data.** The RNAseq read-count data was used to coarsely segment all chromosomes into expressed "segments" and lowly expressed "inter-segments" as described previously [Machné et al., 2017] using the `presegment.R` from the `segmenTools` script collection with parameters `--avg 1000 --minrd 8 --minds 1000 --minsg 1000 --rmlen 1 --favg 100`. **A:** pre-segment and inter-segment length distributions, **B:** Number of time-points (of 25 total) with >0 read-count per segment.

| Filter | Number | Description |
|---|---|---|
| `segmenTier` | 36,927 | output of the segmentation |
| $p_{\text{rain}} \geq 0.85$ | 25,681 | segment cluster "0" |
| $p_{\text{rain}} < 0.85$ | 11,246 | used for segment clustering |
| ORF | 5,795 | annotated ORF in genome release |
| $J_{\text{ORF}} > 0.5$ | 4,489 | segments in coding gene analyses (GO, half-lives) |
| $J_{\text{ORF}} > 0$ | 5,524 | segments in protein complex analysis |
| $J_{\text{AS}} > 0.5$ | 569 | segments considered as antisense transcripts |
| $J_{\Sigma} = 0$ | 9,051 | segments considered as non-coding transcripts |
| $p_{\text{rain}} < 0.05$ | 6,344 | all in Fig. 2A |
| $p_{\text{rain}} < 0.05$ & $J_{\text{ORF}} > 0.5$ | 3,378 | ORF by clusters in Fig. 2A |
| $p_{\text{rain}} < 0.05$ & $J_{\text{AS}} > 0.5$ | 232 | "AS" in Fig. 2A |
| $p_{\text{rain}} < 0.05$ & $J_{\Sigma} = 0$ | 811 | "NC" in Fig. 2A |
| $J_{\text{ORF}} > 0.5$ & half-life data* | 3,189 | for relative protein amplitude |
| $p_{\text{rain}} < 10^{-4}$ | 2,505 | considered in peak width analysis |
| $p_{\text{rain}} < 10^{-4}$ & QC** | 2,357 | two full cycles found |
| $J_{\text{ORF}} > 0.5$ & cluster 1,2,10 & required data*** | 1,026 | HOC genes in PWM prediction |
| $J_{\text{ORF}} > 0.5$ & cluster 6,7,8,19 & required data*** | 1,171 | LOC genes in PWM prediction |

Table S2: **Segment Filtering and Classification.** This table provides an overview of the analyzed total numbers of segments in all analyses of this work. Segments produced from RNAseq data by the `segmenTier` algorithm were classified by their oscillation p-values calculated by the `rain` package ($p_{\text{rain}}$) and by their overlaps with transcribed features annotated in the reference genome (release R64-1-1). quantified each overlap by the Jaccard index, $J = \frac{I}{U}$, where $I$ is the intersect, the number of overlapping nucleotides, and $U$ the union, the number of nucleotides covered by both, the segment and the annotated feature. Segments overlapping with $J_{\text{ORF}} > 0.5$ with an open reading frame (ORF) were classified as transcripts of this ORF and further analyzed. For a comprehensive picture all segments with $J_{\text{ORF}} > 0$, and for each ORF the one with the maximal $J$ were used in protein complex analysis. Segments overlapping with $J_{\text{AS}} > 0.5$ antisense to an ORF were classified as antisense transcripts, and segments not overlapping ($J_{\Sigma} = 0$) with any annotated feature (ORF, antisense to ORF, rRNA, tRNA, snoRNA, snRNA) were classifed as non-coding. *: calculation of relative protein amplitudes require protein half-live data in [Christiano et al., 2014]. **: calculation of peak-width changes required that two full cycles were detected and did not start or end too close the experiment start or end. ***: PWM model prediction of mRNA and protein abundances require mRNA [Geisberg et al., 2014] and protein [Christiano et al., 2014] half-life data, and ribosome density data [Arava et al., 2003].

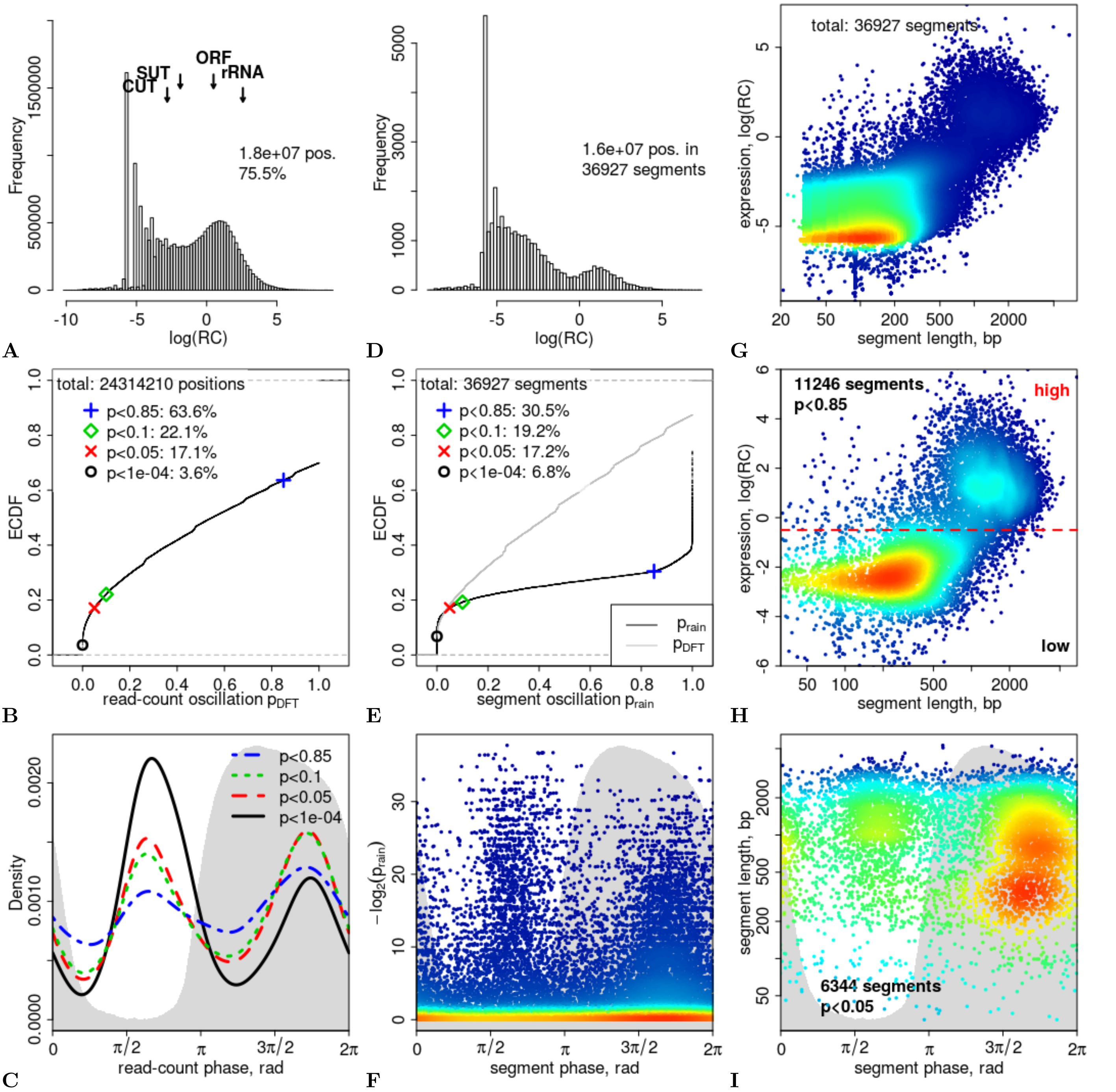


Figure S5: **Global Data Structure.** **A:** distribution of raw combined raw-counts (RC, in RPM), with mean values for annotated features indicated as arrows. $18 \times 10^6$ positions or 75.5% of the nuclear genome was covered by reads. **B:** empirical cumulative distribution function (ECDF) of p-values for oscillation at each genome position, based on the Discrete Fourier Transform (DFT) of the read-counts of the first 19 time-points (2 full cycles) and a permutation test. **C:** distribution of DFT-derived peak phases of oscillating read-counts at different p-value cutoffs. The gray background indicates the dissolved oxygen concentration as a phase reference. **D:** distribution of the mean read-counts (RC, in RPM) of all 36,927 segments. **E:** ECDF of p-values for oscillation of segments, calculated with `rain` [Thaben and Westermark, 2014] (black line) and with a DFT-based permutation test (gray line) as for (B). **F:** 2D histogram (red: highest and blue: lowest density of values) of segment oscillation p-values *vs.* DFT-based peak expression phase. The gray background indicates the dissolved oxygen concentration as a phase reference. **G:** 2D histogram (as in (F)) of segment read-counts (RC,in RPM) *vs.* segment lengths in basepairs (bp). **H:** same as (G) only for the 11,246 segments with $p_{\text{rain}} < 0.85$ (see E) used in all further analyses. **I:** 2D histogram (as in (F)) of segment lengths *vs.* segment phases for the 6,344 segments oscillating at $p_{\text{rain}} < 0.05$. The gray background indicates the dissolved oxygen concentration as a phase reference.

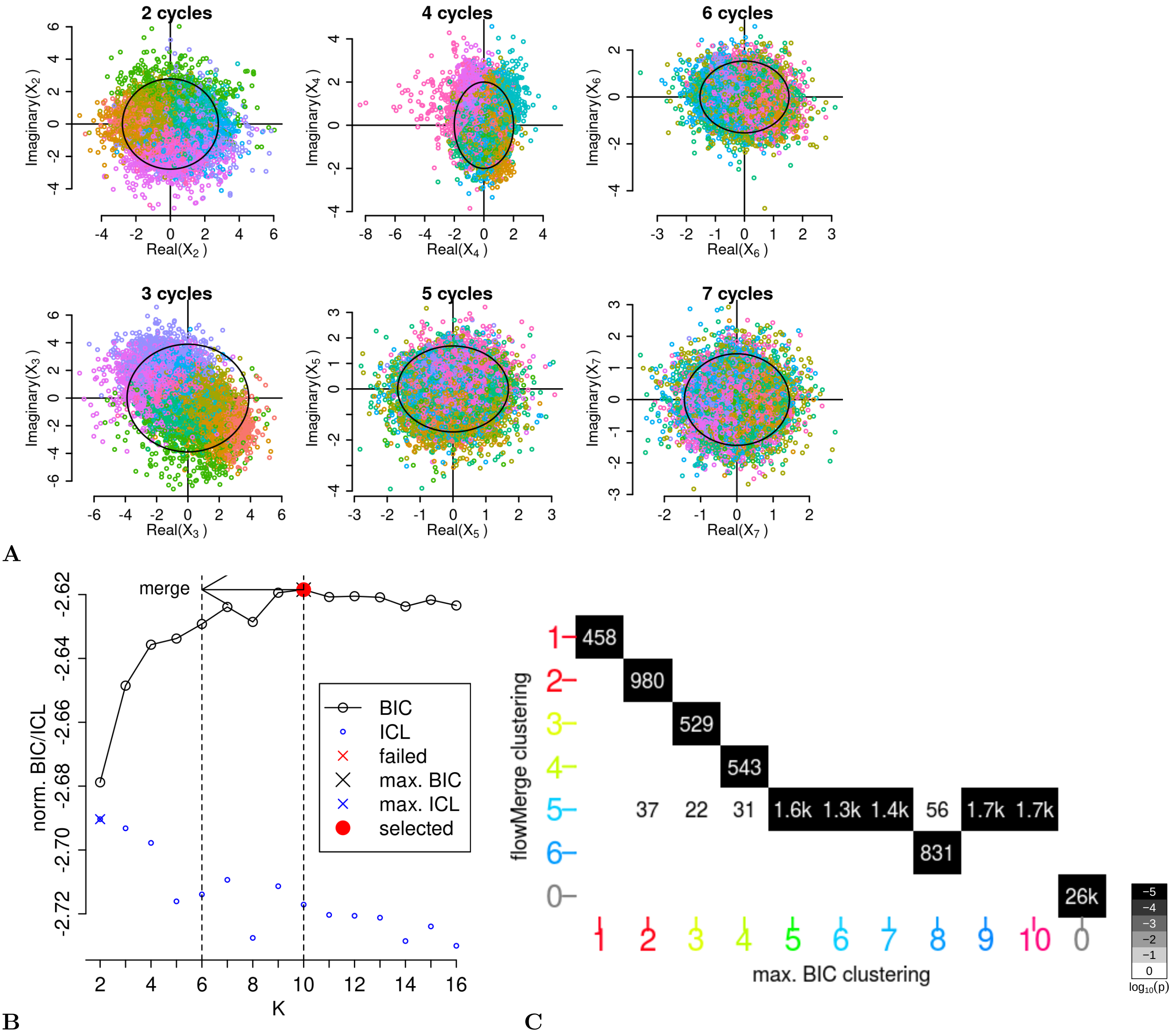


Figure S6: **DFT-based Clustering. A:** Clustering of real and imaginary parts of the Discrete Fourier Transform (DFT) of the time-series by a Gaussian mixture model (with tails) using the R package `flowClust` [Lo et al., 2009] (*via* the script `segmentDynamics.R` and function `clusterTimeSeries2` in `segmenTools`). **B:** The optimal cluster number (K=10) was selected where the maximum BIC (Bayesian Information Content) was reached (`segmenTools` function `plotBIC`). The accessory package `flowMerge` [Finak et al., 2009] (*via* `segmentDynamics.R`) merges 5 clusters while maintaining constant BIC and suggests a new classification into K=6 clusters. **C:** The original and merged clustering were ordered, re-labelled and colored by their mean peak expression phases. The four low-amplitude and mostly non-coding clusters (5, 7, 9 and 10) are merged with the coding cluster 6, here shown as an overlap enrichment profile (`segmenTools`'s `clusterCluster` and `plotOverlaps`).

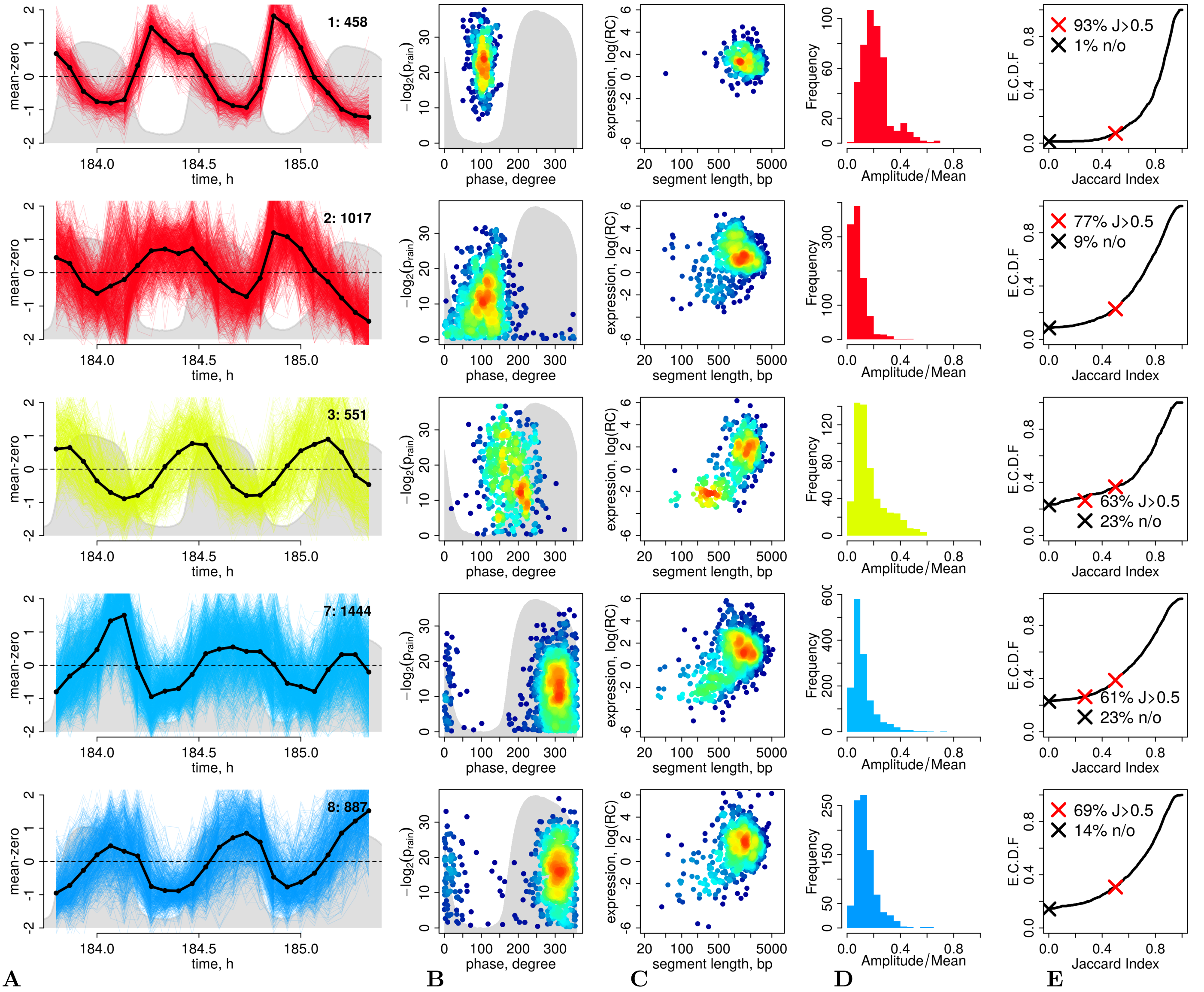


Figure S7: **Segment Cluster Statistics - High Amplitude Clusters.** The high amplitude clusters are shown ordered by their mean phase of expression from top to bottom. **A:** cluster means (black solid lines) and individual segment read-count time series, scaled to a mean of 0. The gray background indicates the dissolved $O_2$ concentration. Plots were generated with the `segmenTools` function `plotClusters`. **B:** 2D histogram of segment oscillation p-values, as calculated by the package `rain`, *vs.* the DFT-derived phases. The background is the averaged dissolved $O_2$ concentration of the first two cycles of the experiment. **C:** 2D histogram of segment expression strenghts (RC: read-counts, RPM) *vs.* segment lengths. **D:** histogram of the relative amplitude, *i.e.*, the ratio of DFT-derived amplitude over mean expression ($X_2/X_0$). **E:** empirical cumulative distribution function (E.C.D.F) of the segment Jaccard indices $J$ with overlapping ORFs. The red **x** indicates the fraction of segments assigned to coding genes at $J > 0.5$, the black **x** indicates the fraction of segments without any ORF overlap.

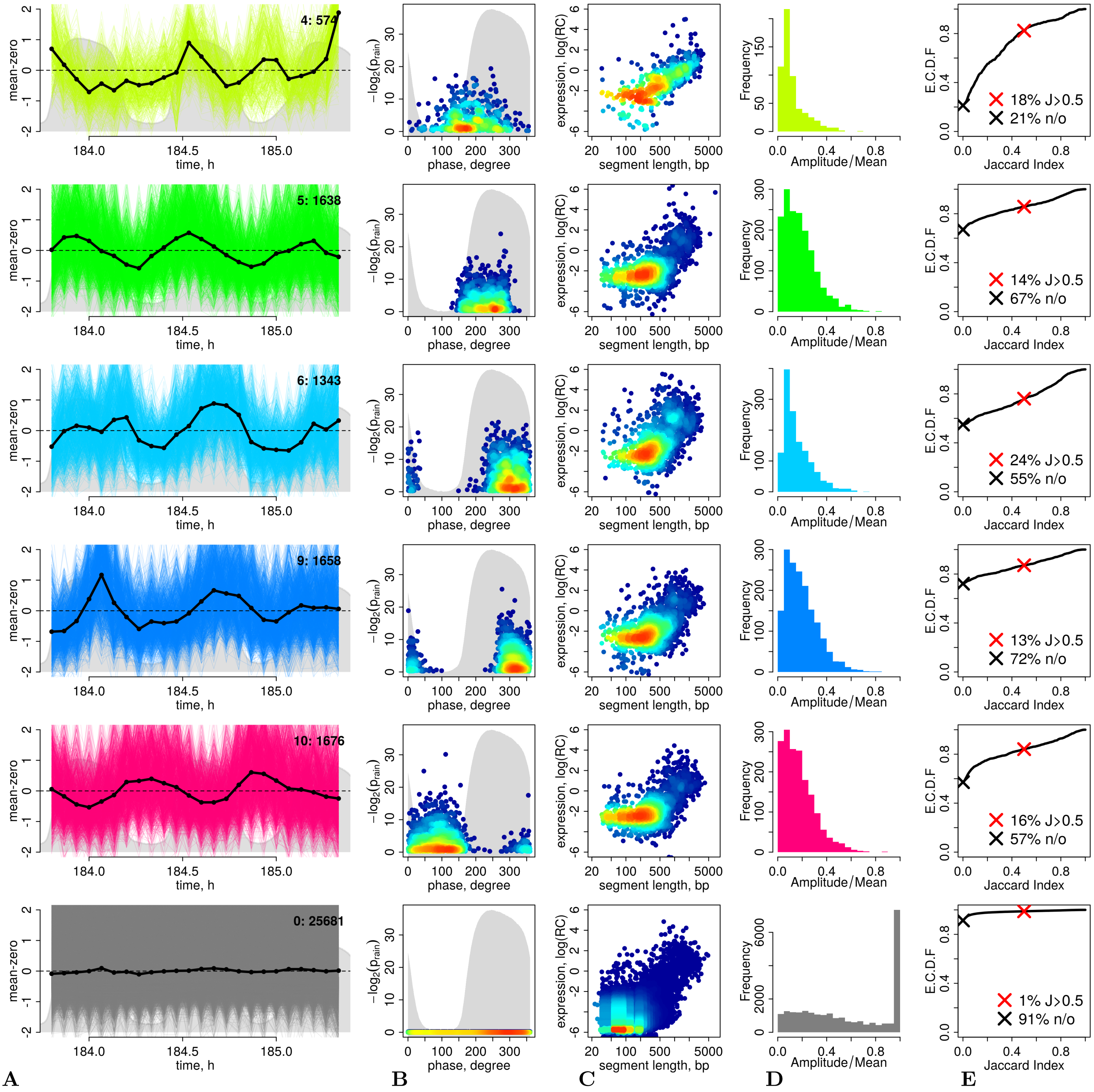


Figure S8: **Segment Cluster Statistics.** As Figure S7 but for low amplitude clusters.

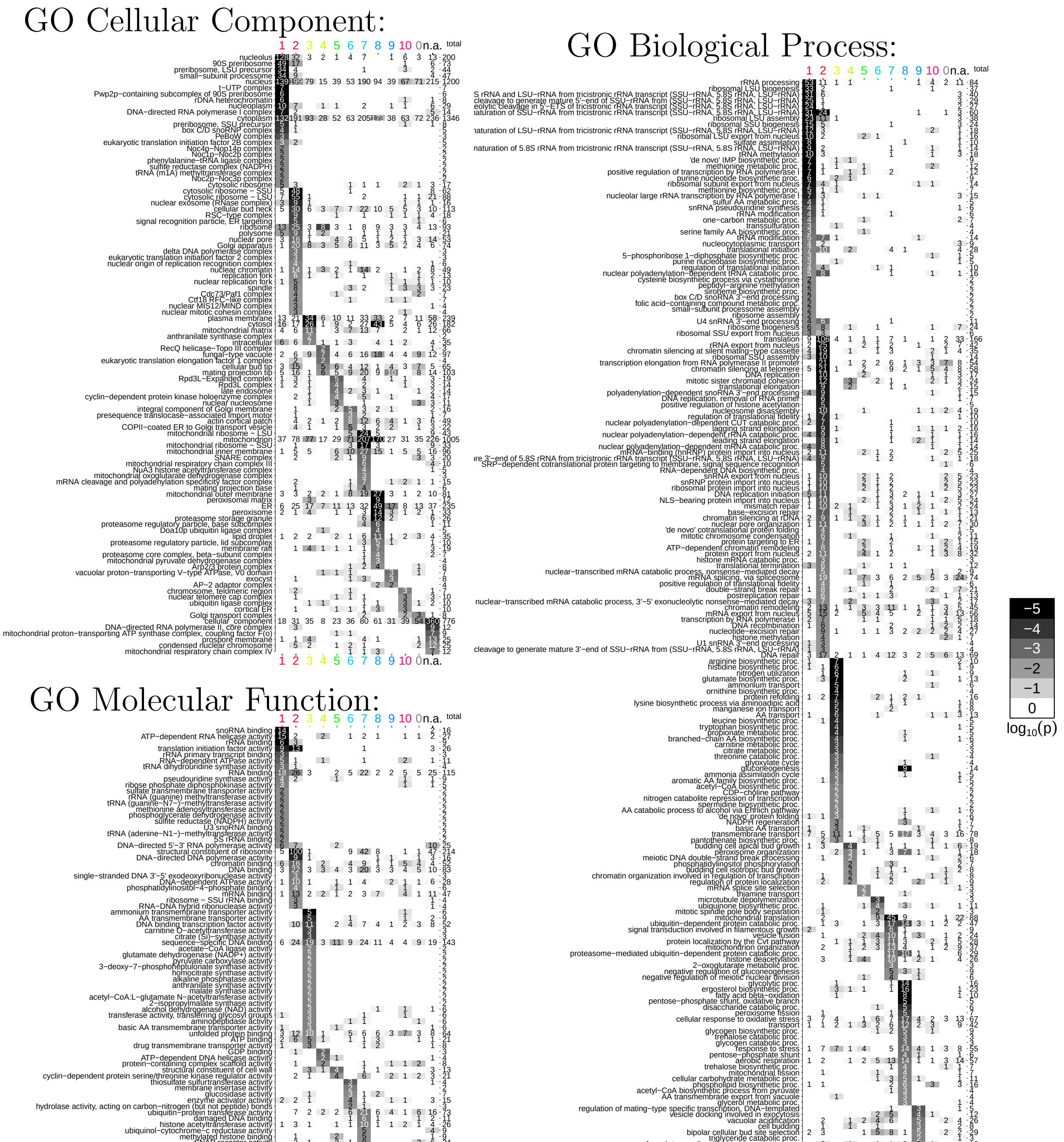


Figure S9: **GO Analysis Enrichment Profiles.** Gene Ontology enrichment profiles for all clusters and all three GO root categories. For cumulative hypergeometric distribution tests the 5,795 annotated protein coding genes was used as the total set (urn size). 4,489 genes that overlap with a clustered segment with $J_{\mathrm{ORF}} > 0.5$ were assigned to the segment's cluster, all other annotated genes are labelled as "n.a." (Tab. S2). The right y-axis provides the total number of genes annotated by each GO term. Only GO terms with any cluster overlap with $p < 0.01$ are shown, and $p_{\mathrm{sort}} = 0.01$ was also used for row sorting (see Methods, "Enrichment Profiles").

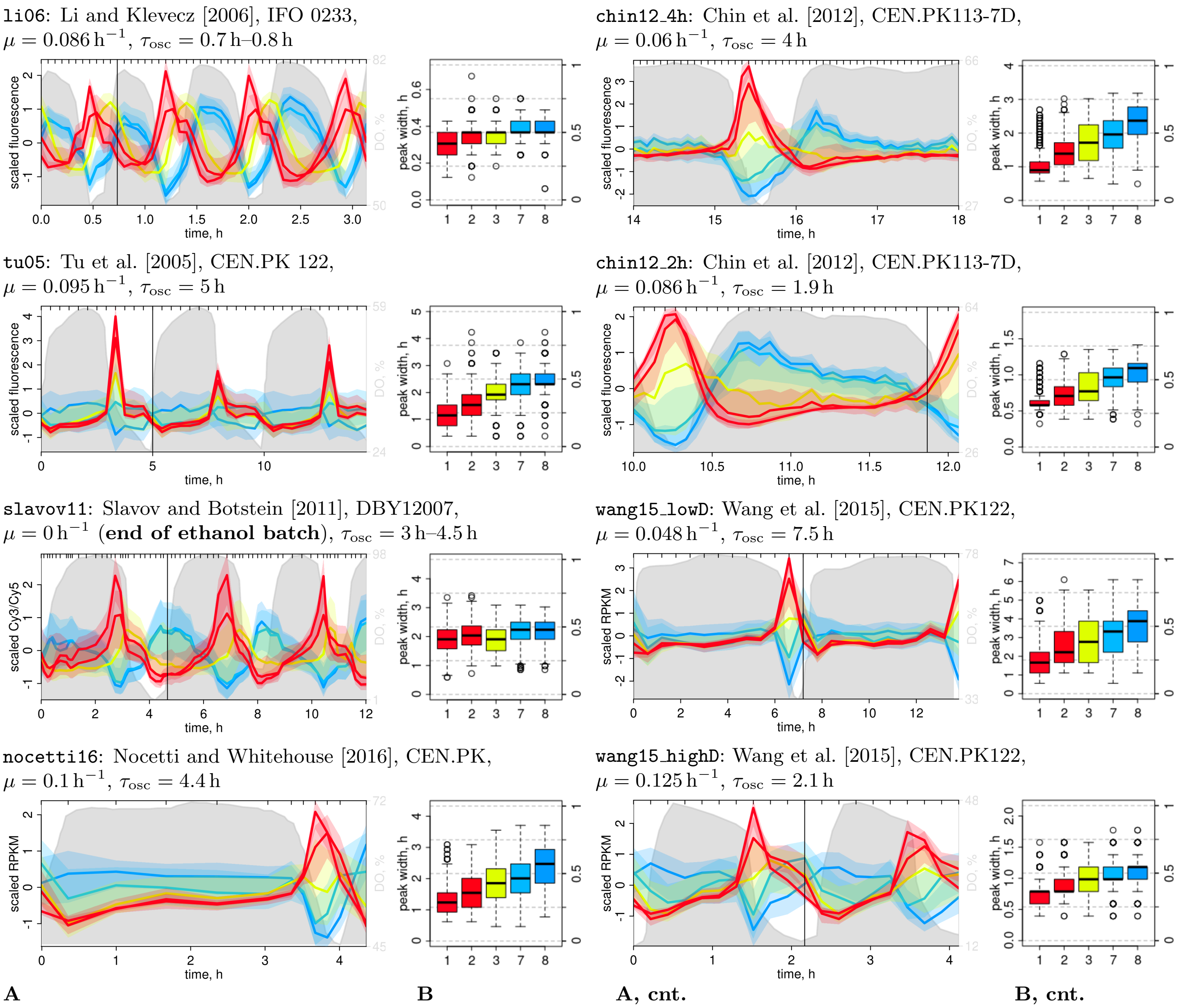


Figure S10: **Oscillatory Transcriptome Meta-Analysis.** Meta-analysis of the transcription program in previously published data sets from the indicated publications, using the clustering described herein and only showing the major protein coding clusters 1 (Ribi), 2 (RP), 3 (AA), 7 (mRP) and 8 (S/C). Note that the right columns (A, cnt. and B, cnt.) show experiments run at different dilution rates, these were compared for the analysis shown in Figure 3D. **A:** The yeast strains, growth rates $\mu$ (assuming stable cultures equal to the reported dilution rates $\phi$), and the oscillation periods $\tau_{osc}$ are indicated for each experiment. The horizontal black line demarks the first cycle, and only this first cycle was used for peak width estimation in (B). The li06 and tu05 data sets were taken from the supporting material from and re-normalized as described in Machné and Murray [2012]. Cluster means (solid lines) and 25%/75% quantiles (transparent areas) plotted with segmenTools's plotClusters. Gray areas indicate the dissolved oxygen concentrations digitized with engauge from the published figures. **B:** Boxplots of cluster distributions of transcript abundance peak widths. The peak widths of each transcript were calculated for the first cycle of each experiment (horizontal black lines in (A)) as the time spent above the temporal mean of the transcript. The right y-axis shows peak widths as a fraction of $\tau_{osc}$. Data that were not sampled equispaced were interpolated at equispaced time points using the minimal time step of the original sampling. The means of cluster 2 and 8 were used for the analysis shown in Figure 3E.

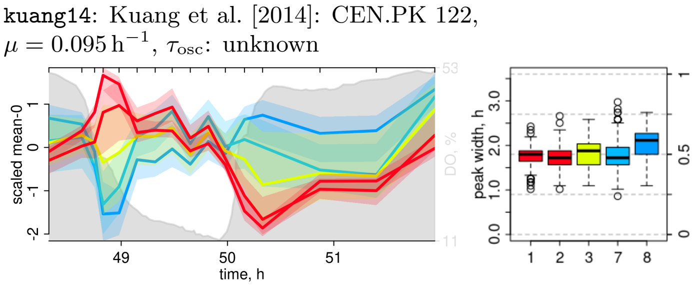


Figure S11: For the experiment by Kuang et al. [2014] the publication (Fig. 1 therein) leaves it unclear, whether a full cycle was sampled and what the period was. Note that cluster 7 and 8 (cyan and blue) transcript abundances are higher at the last time point than at the first time point, indicating that the experiment did not cover a full cycle. We therefore excluded this data set from peak width analysis.

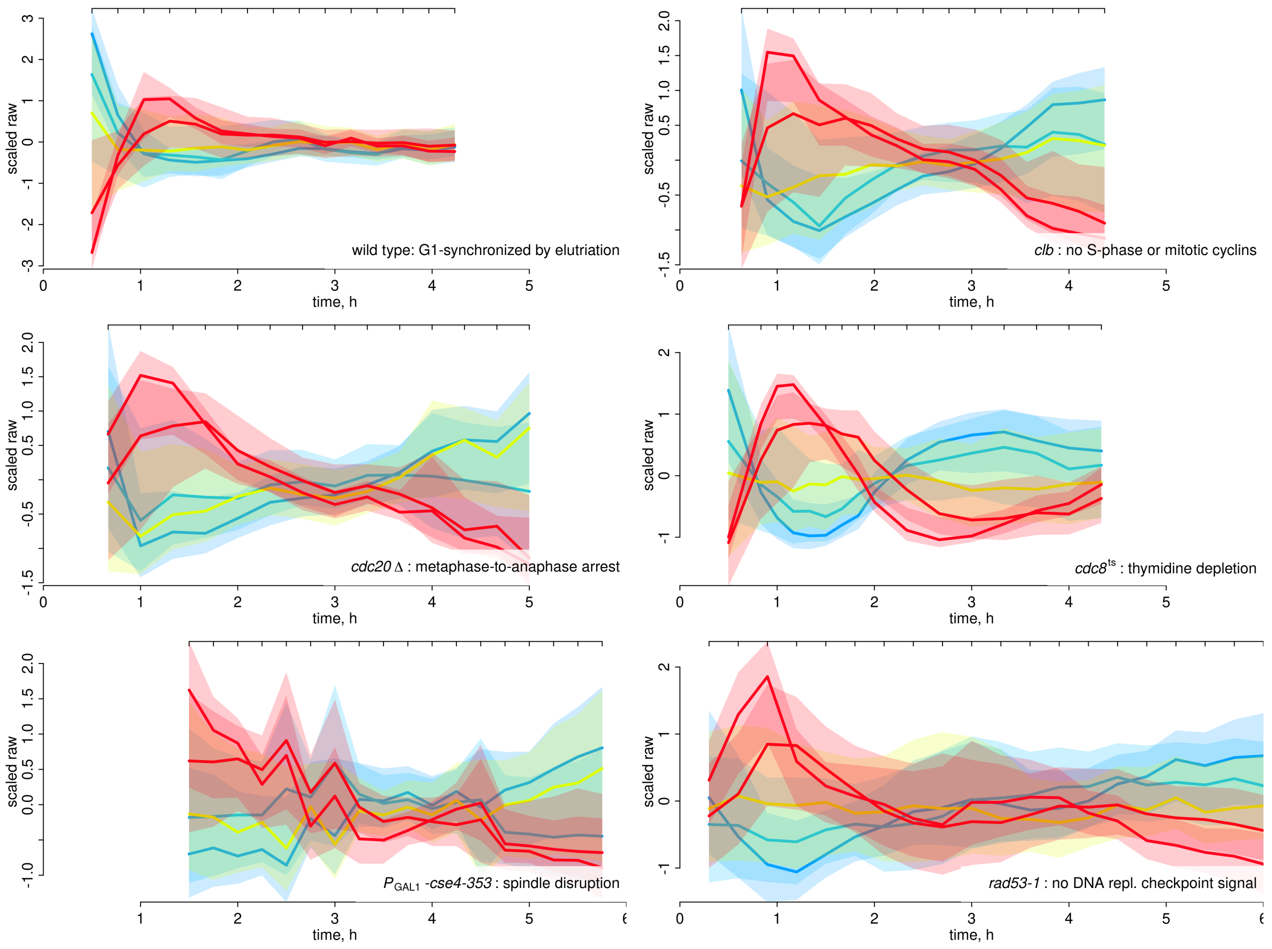


Figure S12: **Cell-Cycle Transcriptomes.** As Figure S10A but for cell cycle arrest experiments from Orlando et al. [2008] and [Bristow et al., 2014].

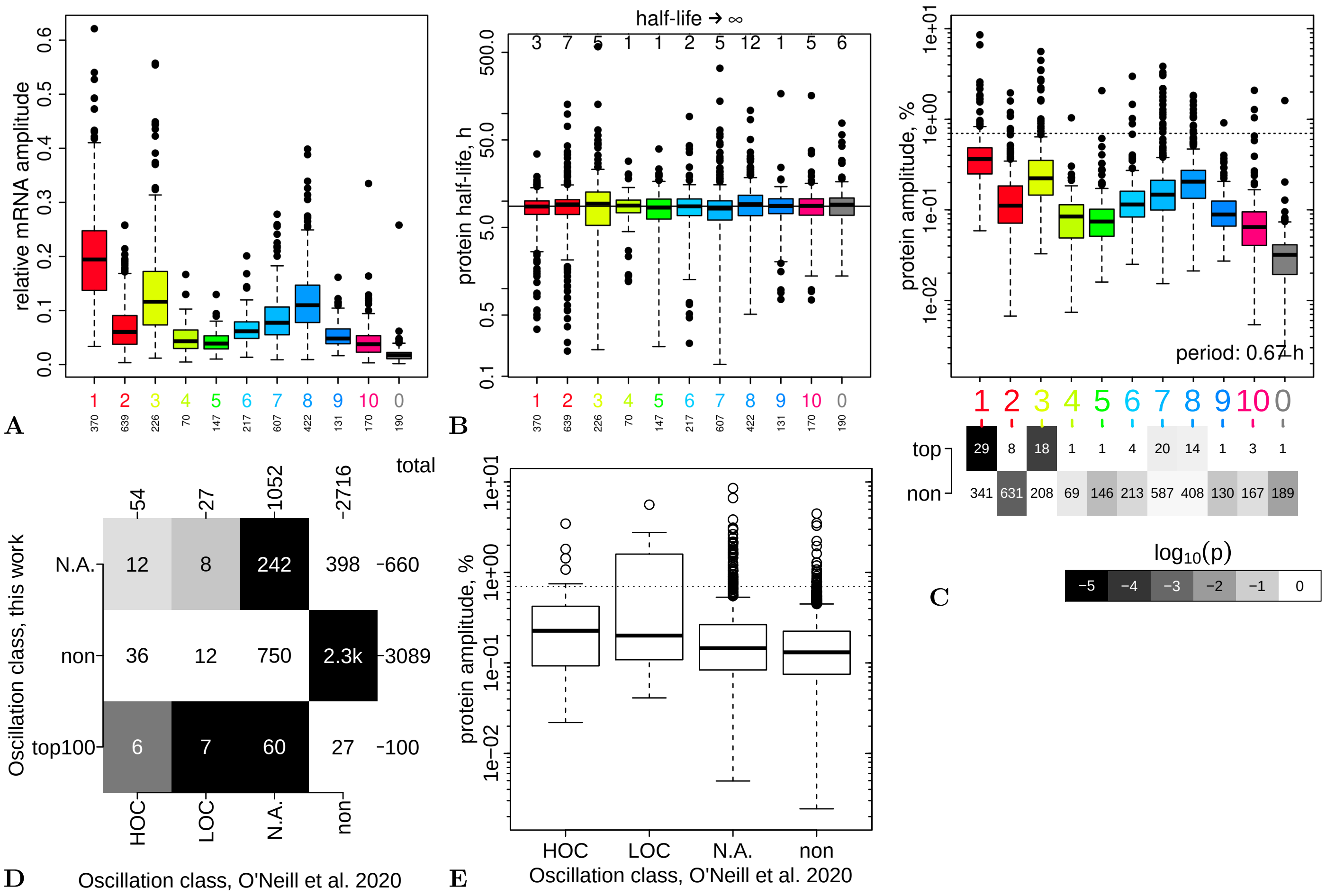


Figure S13: **Prediction of Relative Protein Amplitudes. A–C:** For this analysis 3,189 coding genes covered by a segment with $J > 0.5$ and with available protein half-life data by [Christiano et al., 2014] were considered (Dataset S3). **A:** Cluster distributions of relative mRNA amplitudes $A_\mathrm{R}$ (see Methods) are shown as box-plots. The total number of transcripts in each cluster is shown below the x-axis. **B:** cluster distributions of protein half-lives from [Christiano et al., 2014]. The horizontal line indicates the mean half-live of ca. 9 h; 48 proteins were annotated to have infinite half-lives (top axis). **C:** cluster distributions of relative protein amplitudes ($A_\mathrm{P}$), calculated from mRNA amplitudes (A) and protein half-lives (B) with the model by Lück et al. [2014] (see Methods). The dashed horizontal line indicates the top-100 cut-off. The bottom panel shows a cluster enrichment analysis of the top 100 proteins with $p_\mathrm{txt} = 5 \cdot 10^{-3}$ (white text) and $p_\mathrm{min} = 10^{-5}$ (black fields). **D & E:** Comparison of our predicted top 100 oscillating proteins (E) and protein amplitudes (F) to the proteome-based oscillators from a long period oscillation, classifed into HOC phase- and LOC phase-specific or non-oscillatory ("non"), by O' Neill et al. [2020]. For (E) all 3,849 coding genes covered by a segment with $J > 0.5$ were considered, and "N.A." indicates the 660 genes in this set that had no protein half-life data for amplitude prediction (our data) or the 1,052 proteins that were not detected in the proteome measurements. The enrichment analysis in (E) was plotted with the same p-value cut-offs as (C). The N.A. class of the proteome data is strongly enriched ($p = 4 \cdot 10^{-12}$) in the top oscillators of our data. This class contains transcription factors (e.g. BDF2, CLB2, GZF3, MET28, SWI5, MSN4) that are expressed at low abundance and may have escaped detection in the proteomics study. The six HOC phase-specific proteins that overlap with our top 100 list ($p = 0.002$) are (the products of): CIT2, FCY2, THI7, PRY2, MCD1, NSA2. The seven LOC phase-specific proteins ($p = 3.7 \cdot 10^{-6}$) are: YKL164C, ATO3, NCE103, YDR343C, MDH2, ITR1, ERG1. Several of these are discussed in the manuscript. See Table S3 for functions. The protein amplitudes in (F) are the same as in (D); the horizontal line indicates the top 100 cut-off.

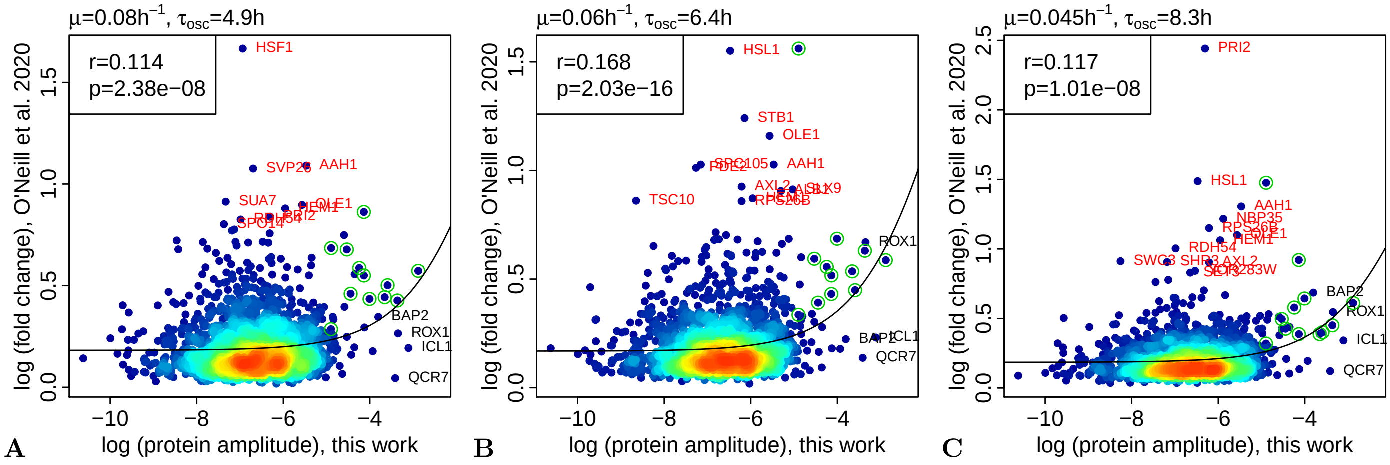


Figure S14: **Genome-Wide Comparison of Protein Amplitudes:** predicted amplitudes (our data, x-axis, Fig. S13) vs. measured protein amplitudes at three different culture growth rates (y-axis) by O' Neill et al. [2020]. On a genome-wide scale the correlation between amplitudes estimated from proteome data and from mRNA amplitudes and protein half-lives is weak but significantly positive. The top left legends provide the Pearson correlation and p-value (R's `cor.test`) and the black line is a linear regression of the non-logged raw data. Green circles indicate the consensus genes, explicitly stated in the caption for Figure S13D. The genes indicated by name (red and black text) are classified as oscillators in one, but not the other data set. All data is available in Dataset S3.

| ID.segment | CL | ONeill | J | ID.gene | Name | rA.RNA | t05.protein | rA.protein | Description |
|---|---|---|---|---|---|---|---|---|---|
| sg0488_3 | 1 | N.A. | 0.92 | YGR055W | MUP1 | 0.62 | 0.57 | 0.09 | High affinity methionine permease, integral mem... |
| sg1555_3 | 1 | N.A. | 0.67 | YDR184C | ATC1 | 0.30 | 0.34 | 0.07 | Nuclear protein, possibly involved in regulatio... |
| sg2038_2 | 3 | LOC | 0.58 | YKL164C | PIR1* | 0.16 | 0.20 | 0.06 | O-glycosylated protein required for cell wall s... |
| sg1666_5 | 3 | non | 0.81 | YER065C | ICL1 | 0.55 | 1.03 | 0.04 | Isocitrate lyase, catalyzes the formation of su... |
| sg1224_17 | 7 | N.A. | 0.56 | YPL187W | MF(ALPHA)1 | 0.12 | 0.23 | 0.04 | Mating pheromone alpha-factor, made by alpha ce... |
| sg1283_8 | 3 | non | 0.72 | YPR065W | ROX1 | 0.22 | 0.49 | 0.04 | Heme-dependent repressor of hypoxic genes; cont... |
| sg1440_13 | 3 | HOC | 0.57 | YCR005C | CIT2 | 0.56 | 1.40 | 0.03 | Citrate synthase, catalyzes the condensation of... |
| sg1634_20 | 7 | non | 0.56 | YDR529C | QCR7 | 0.08 | 0.17 | 0.03 | Subunit 7 of the ubiquinol cytochrome-c reducta... |
| sg0201_8 | 7 | N.A. | 0.78 | YDL070W | BDF2 | 0.28 | 0.71 | 0.03 | Protein involved in transcription initiation at... |
| sg1431_6 | 7 | N.A. | 0.95 | YCL067C | HMLALPHA2* | 0.11 | 0.26 | 0.03 | Silenced copy of ALPHA2 at HML; homeobox-domain... |
| sg0497_14 | 6 | N.A. | 0.73 | YGR111W | | 0.10 | 0.24 | 0.03 | Putative protein of unknown function; green flu... |
| sg1603_5 | 3 | LOC | 0.60 | YDR384C | ATO3 | 0.31 | 0.94 | 0.03 | Plasma membrane protein, regulation pattern sug... |
| sg1010_5 | 3 | N.A. | 0.95 | YNL327W | EGT2 | 0.15 | 0.42 | 0.03 | Glycosylphosphatidylinositol (GPI)-anchored cel... |
| sg1066_8 | 1 | LOC | 0.51 | YNL036W | NCE103 | 0.16 | 0.47 | 0.03 | Carbonic anhydrase; poorly transcribed under ae... |
| sg1238_7 | 1 | N.A. | 0.51 | YPL135W | ISU1 | 0.24 | 0.81 | 0.02 | Conserved protein of the mitochondrial matrix, ... |
| sg1381_11 | 3 | non | 0.86 | YBR068C | BAP2 | 0.23 | 0.86 | 0.02 | High-affinity leucine permease, functions as a ... |
| sg1405_6 | 7 | N.A. | 0.69 | YBR173C | UMP1 | 0.12 | 0.41 | 0.02 | Short-lived chaperone required for correct matu... |
| sg2346_4 | 7 | N.A. | 0.80 | YNL115C | | 0.17 | 0.61 | 0.02 | Putative protein of unknown function; green flu... |
| sg0086_12 | 7 | N.A. | 0.87 | YBR158W | AMN1 | 0.11 | 0.39 | 0.02 | Protein required for daughter cell separation, ... |
| sg2501_15 | 1 | N.A. | 0.76 | YOR342C | | 0.15 | 0.56 | 0.02 | Putative protein of unknown function; green flu... |
| sg0265_3 | 3 | N.A. | 0.58 | YDR222W | YDR222W | 0.45 | 1.96 | 0.02 | Protein of unknown function; green fluorescent ... |
| sg1699_3 | 10 | N.A. | 0.78 | YFL065C | | 0.33 | 1.40 | 0.02 | Putative protein of unknown function; induced b... |
| sg0228_8 | 5 | N.A. | 0.83 | YDR077W | SED1 | 0.06 | 0.22 | 0.02 | Major stress-induced structural GPI-cell wall g... |
| sg2432_4 | 7 | N.A. | 0.52 | YOR028C | CIN5 | 0.16 | 0.65 | 0.02 | Basic leucine zipper (bZIP) transcription facto... |
| sg0748_9 | 2 | non | 0.88 | YKL139W | CTK1 | 0.05 | 0.19 | 0.02 | Catalytic (alpha) subunit of C-terminal domain ... |
| sg1199_4 | 1 | N.A. | 0.86 | YOR355W | GDS1 | 0.20 | 0.88 | 0.02 | Protein of unknown function, required for growt... |
| sg1592_1 | 8 | LOC | 0.73 | YDR343C | HXT6* | 0.27 | 1.27 | 0.02 | High-affinity glucose transporter of the major ... |
| sg1665_6 | 1 | HOC | 0.79 | YER056C | FCY2 | 0.20 | 0.89 | 0.02 | Purine-cytosine permease, mediates purine (aden... |
| sg1591_1 | 8 | N.A. | 0.55 | YDR342C | HXT7* | 0.26 | 1.27 | 0.02 | High-affinity glucose transporter of the major ... |
| sg1740_3 | 7 | N.A. | 0.91 | YGL219C | MDM34 | 0.12 | 0.53 | 0.02 | Mitochondrial component of the ERMES complex th... |
| sg0119_2 | 1 | N.A. | 0.77 | YCL058W-A | ADF1 | 0.23 | 1.18 | 0.02 | Transcriptional repressor encoded by the antise... |
| sg0791_4 | 1 | N.A. | 0.58 | YKR093W | PTR2 | 0.30 | 1.58 | 0.02 | Integral membrane peptide transporter, mediates... |
| sg1209_37 | 3 | N.A. | 0.85 | YPL265W | DIP5 | 0.44 | 2.61 | 0.02 | Dicarboxylic amino acid permease, mediates high... |
| sg0937_8 | 3 | non | 0.87 | YMR011W | HXT2 | 0.21 | 1.12 | 0.02 | High-affinity glucose transporter of the major ... |
| sg1293_2 | 7 | N.A. | 0.76 | YPR119W | CLB2 | 0.11 | 0.52 | 0.02 | B-type cyclin involved in cell cycle progressio... |
| sg1316_4 | 0 | N.A. | 0.56 | YPR203W | | 0.26 | 1.40 | 0.02 | Putative protein of unknown function |
| sg2398_10 | 3 | LOC | 0.76 | YOL126C | MDH2 | 0.45 | 2.80 | 0.02 | Cytoplasmic malate dehydrogenase, one of three ... |
| sg1627_3 | 2 | LOC | 0.86 | YDR497C | ITR1 | 0.08 | 0.37 | 0.02 | Myo-inositol transporter with strong similarity... |
| sg0756_9 | 8 | N.A. | 0.78 | YKL096W | CWP1 | 0.10 | 0.51 | 0.02 | Cell wall mannoprotein that localizes specifica... |
| sg0124_2 | 1 | N.A. | 0.97 | YCL036W | GFD2 | 0.35 | 2.12 | 0.02 | Protein of unknown function, identified as a hi... |
| sg0592_2 | 6 | N.A. | 0.67 | YHR152W | SPO12 | 0.09 | 0.47 | 0.01 | Nucleolar protein of unknown function, positive... |
| sg1599_5 | 7 | N.A. | 0.90 | YDR369C | XRS2 | 0.10 | 0.52 | 0.01 | Protein required for DNA repair; component of t... |
| sg0858_16 | 8 | HOC | 0.88 | YLR237W | THI7 | 0.17 | 0.98 | 0.01 | Plasma membrane transporter responsible for the... |
| sg0171_3 | 8 | non | 0.64 | YDL204W | RTN2 | 0.21 | 1.46 | 0.01 | Protein of unknown function; has similarity to ... |
| sg2478_4 | 10 | N.A. | 0.56 | YOR226C | ISU2 | 0.12 | 0.74 | 0.01 | Conserved protein of the mitochondrial matrix, ... |
| sg0940_12 | 7 | non | 0.92 | YMR028W | TAP42 | 0.03 | 0.14 | 0.01 | Essential protein involved in the TOR signaling... |
| sg1582_3 | 1 | N.A. | 0.70 | YDR309C | GIC2 | 0.08 | 0.50 | 0.01 | Redundant rho-like GTPase Cdc42p effector; homo... |
| sg1420_10 | 2 | non | 0.94 | YBR260C | RGD1 | 0.04 | 0.24 | 0.01 | GTPase-activating protein (RhoGAP) for Rho3p an... |
| sg1910_1 | 7 | N.A. | 0.59 | YIL119C | RPI1 | 0.26 | 1.99 | 0.01 | Putative transcriptional regulator; overexpress... |
| sg1825_21 | 8 | LOC | 0.84 | YGR175C | ERG1 | 0.19 | 1.39 | 0.01 | Squalene epoxidase, catalyzes the epoxidation o... |
| sg1545_1 | 7 | N.A. | 0.77 | YDR146C | SWI5 | 0.07 | 0.45 | 0.01 | Transcription factor that activates transcripti... |
| sg0921_13 | 2 | N.A. | 0.91 | YML065W | ORC1 | 0.08 | 0.57 | 0.01 | Largest subunit of the origin recognition compl... |
| sg2171_4 | 8 | non | 0.85 | YLR286C | CTS1 | 0.17 | 1.39 | 0.01 | Endochitinase, required for cell separation aft... |
| sg0776_6 | 2 | HOC | 0.74 | YKR013W | PRY2 | 0.06 | 0.46 | 0.01 | Protein of unknown function |
| sg0269_3 | 7 | N.A. | 0.50 | YDR262W | | 0.13 | 0.99 | 0.01 | Putative protein of unknown function; green flu... |
| sg0850_4 | 8 | N.A. | 0.66 | YLR190W | MMR1 | 0.07 | 0.56 | 0.01 | Phosphorylated protein of the mitochondrial out... |
| sg1059_11 | 8 | N.A. | 0.69 | YNL066W | SUN4 | 0.17 | 1.45 | 0.01 | Cell wall protein related to glucanases, possib... |
| sg1555_2 | 10 | N.A. | 0.62 | YDR185C | UPS3 | 0.12 | 0.97 | 0.01 | Mitochondrial protein of unknown function; simi... |
| sg1639_13 | 4 | N.A. | 0.60 | YEL076C | | 0.17 | 1.40 | 0.01 | Putative protein of unknown function |
| sg0618_12 | 6 | non | 0.69 | YIL123W | SIM1 | 0.07 | 0.52 | 0.01 | Protein of the SUN family (Sim1p, Uth1p, Nca3p,... |
| sg0830_11 | 3 | non | 0.80 | YLR079W | SIC1 | 0.08 | 0.61 | 0.01 | Inhibitor of Cdc28-Clb kinase complexes that co... |
| sg1279_2 | 1 | non | 0.80 | YPR035W | GLN1 | 0.47 | 6.70 | 0.01 | Glutamine synthetase (GS), synthesizes glutamin... |
| sg0549_9 | 3 | N.A. | 0.85 | YHL027W | RIM101 | 0.09 | 0.71 | 0.01 | Transcriptional repressor involved in response ... |
| sg0922_2 | 1 | N.A. | 0.83 | YML060W | OGG1 | 0.16 | 1.48 | 0.01 | Mitochondrial glycosylase/lyase that specifical... |
| sg0748_13 | 9 | non | 0.53 | YKL137W | CMC1 | 0.16 | 1.57 | 0.01 | Evolutionarily conserved copper-binding protein... |
| sg0267_17 | 3 | N.A. | 0.68 | YDR247W | VHS1 | 0.19 | 1.89 | 0.01 | Cytoplasmic serine/threonine protein kinase; id... |
| sg2399_10 | 7 | N.A. | 0.52 | YOL122C | SMF1 | 0.13 | 1.25 | 0.01 | Divalent metal ion transporter with a broad spe... |
| sg1209_17 | 1 | N.A. | 0.84 | YPL274W | SAM3 | 0.32 | 3.91 | 0.01 | High-affinity S-adenosylmethionine permease, re... |
| sg2036_14 | 6 | N.A. | 0.51 | YKL178C | STE3 | 0.08 | 0.69 | 0.01 | Receptor for a factor pheromone, couples to MAP... |
| sg0030_2 | 1 | N.A. | 0.72 | YBL081W | | 0.23 | 2.53 | 0.01 | Non-essential protein of unknown function; null... |
| sg0851_8 | 7 | non | 0.72 | YLR206W | ENT2 | 0.04 | 0.39 | 0.01 | Epsin-like protein required for endocytosis and... |
| sg1381_16 | 8 | N.A. | 0.68 | YBR062C | | 0.17 | 1.87 | 0.01 | Protein of unknown function that interacts with... |
| sg1000_13 | 8 | non | 0.82 | YMR297W | PRC1 | 0.06 | 0.56 | 0.01 | Vacuolar carboxypeptidase Y (proteinase C; CPY)... |
| sg1362_4 | 1 | N.A. | 0.52 | YBL028C | | 0.21 | 2.41 | 0.01 | Protein of unknown function that may interact w... |
| sg1967_11 | 3 | N.A. | 0.89 | YJL110C | GZF3 | 0.20 | 2.28 | 0.01 | GATA zinc finger protein and Dal80p homolog tha... |
| sg1038_6 | 1 | N.A. | 0.81 | YNL191W | DUG3 | 0.42 | 7.11 | 0.01 | Probable glutamine amidotransferase, forms a co... |
| sg2100_7 | 1 | non | 0.62 | YLL062C | MHT1 | 0.49 | 10.35 | 0.01 | S-methylmethionine-homocysteine methyltransfera... |
| sg0056_2 | 7 | N.A. | 0.60 | YBR038W | CHS2 | 0.07 | 0.74 | 0.01 | Chitin synthase II; catalyzes transfer of N-ace... |
| sg0417_3 | 1 | non | 0.97 | YFR030W | MET10 | 0.42 | 7.88 | 0.01 | Subunit alpha of assimilatory sulfite reductase... |
| sg1642_7 | 3 | non | 0.77 | YEL063C | CAN1 | 0.12 | 1.35 | 0.01 | Plasma membrane arginine permease, requires pho... |
| sg1836_5 | 1 | non | 0.52 | YGR280C | PXR1 | 0.29 | 4.05 | 0.01 | Essential protein involved in rRNA and snoRNA m... |
| sg2522_15 | 2 | N.A. | 0.88 | YPL253C | VIK1 | 0.09 | 0.94 | 0.01 | Protein that forms a complex with Kar3p at the ... |
| sg2017_2 | 1 | non | 0.66 | YJR137C | MET5 | 0.47 | 10.28 | 0.01 | Sulfite reductase beta subunit, involved in ami... |
| sg0804_5 | 1 | N.A. | 0.70 | YLL027W | ISA1 | 0.13 | 1.39 | 0.01 | Mitochondrial matrix protein involved in biogen... |
| sg2430_4 | 7 | N.A. | 0.91 | YOR023C | AHC1 | 0.09 | 0.99 | 0.01 | Subunit of the Ada histone acetyltransferase co... |
| sg0758_7 | 1 | non | 0.98 | YKL078W | DHR2 | 0.37 | 6.51 | 0.01 | Predominantly nucleolar DEAH-box ATP-dependent ... |
| sg0334_6 | 1 | non | 0.83 | YEL071W | DLD3 | 0.54 | 15.50 | 0.01 | D-lactate dehydrogenase, part of the retrograde... |
| sg0642_10 | 1 | N.A. | 0.85 | YIL019W | FAF1 | 0.20 | 2.53 | 0.01 | Protein required for pre-rRNA processing and 40... |
| sg2131_9 | 8 | N.A. | 0.80 | YLR108C | | 0.23 | 3.22 | 0.01 | Protein of unknown function; green fluorescent ... |
| sg0212_5 | 2 | HOC | 0.84 | YDL003W | MCD1 | 0.07 | 0.79 | 0.01 | Essential subunit of the cohesin complex requir... |
| sg0750_6 | 2 | N.A. | 0.89 | YKL125W | RRN3 | 0.14 | 1.66 | 0.01 | Protein required for transcription of rDNA by R... |
| sg1682_1 | 1 | HOC | 0.57 | YER126C | NSA2 | 0.22 | 3.02 | 0.01 | Protein constituent of 66S pre-ribosomal partic... |
| sg2245_16 | 3 | non | 0.85 | YMR015C | ERG5 | 0.17 | 2.11 | 0.01 | C-22 sterol desaturase, a cytochrome P450 enzym... |
| sg1179_4 | 3 | N.A. | 0.77 | YOR264W | DSE3 | 0.12 | 1.48 | 0.01 | Daughter cell-specific protein, may help establ... |
| sg1771_9 | 1 | N.A. | 0.90 | YGL077C | HNM1 | 0.16 | 2.00 | 0.01 | Choline/ethanolamine transporter; involved in t... |
| sg2608_8 | 8 | non | 0.91 | YPR155C | NCA2 | 0.12 | 1.39 | 0.01 | Protein involved in regulation of mitochondrial... |
| sg0486_1 | 7 | non | 0.58 | YGR046W | TAM41 | 0.07 | 0.84 | 0.01 | Mitochondrial protein involved in protein impor... |
| sg0759_48 | 8 | N.A. | 0.82 | YKL062W | MSN4 | 0.14 | 1.88 | 0.01 | Transcriptional activator related to Msn2p; act... |
| sg2301_7 | 1 | non | 0.79 | YMR300C | ADE4 | 0.41 | 9.74 | 0.01 | Phosphoribosylpyrophosphate amidotransferase (P... |
| sg1943_24 | 1 | N.A. | 0.56 | YIR017C | MET28 | 0.41 | 9.58 | 0.01 | Basic leucine zipper (bZIP) transcriptional act... |

Table S3: Top 100 predicted protein oscillators. ID.segment: ID of the segment overlapping with the gene (Jaccard index $J > 0.5$ with the gene ORF). CL.segment: cluster assignment of the segment. ONeill: the HOC/LOC classification by [O' Neill et al., 2020], see Figure S13D,E. J: Jaccard index of segment:gene overlap. ID.gene/Name: systematic and standard names of the gene. t05.protein: protein half-lives (in hours) from [Christiano et al., 2014], and rA: relative oscillation amplitudes (% of mean) for RNA (our data) and proteins (calculated via the model by [Lück et al., 2014]). See Figure S13 for details. The table is sorted by the relative protein amplitude (rA.protein). Description: protein annotation in the gff3 genome release file, cut at 50 letters. (*) these genes didn't have an official standard name in the genome release used for analysis, but do have the indicated name at the current SGD website (2020).

| description | category | genome | cluster | enrichment | cluster % of bin | p-value |
|---|---|---|---|---|---|---|
| plasma membrane | GO:0005886 | 131 | 15 | 4.4 | 11.5 | 9.2E-07 |
| transmembrane transport | GO:0055085 | 33 | 8 | 9.3 | 24.2 | 1.3E-06 |
| fungal-type cell wall | GO:0009277 | 38 | 7 | 7.1 | 18.4 | 4.2E-05 |
| glucose transmembrane transporter activity | GO:0005355 | 5 | 3 | 23.1 | 60.0 | 1.6E-04 |
| fructose transmembrane transporter activity | GO:0005353 | 5 | 3 | 23.1 | 60.0 | 1.6E-04 |
| mannose transmembrane transporter activity | GO:0015578 | 5 | 3 | 23.1 | 60.0 | 1.6E-04 |
| structural constituent of cell wall | GO:0005199 | 6 | 3 | 19.2 | 50.0 | 3.2E-04 |
| sulfite reductase (NADPH) activity | GO:0004783 | 2 | 2 | 38.5 | 100.0 | 6.7E-04 |
| sulfite reductase complex (NADPH) | GO:0009337 | 2 | 2 | 38.5 | 100.0 | 6.7E-04 |
| integral component of plasma membrane | GO:0005887 | 9 | 3 | 12.8 | 33.3 | 1.3E-03 |
| extrinsic component of mitochondrial inner membrane | GO:0031314 | 10 | 3 | 11.5 | 30.0 | 1.8E-03 |
| glyoxylate cycle | GO:0006097 | 3 | 2 | 25.7 | 66.7 | 2.0E-03 |
| hexose transmembrane transport | GO:0008645 | 3 | 2 | 25.7 | 66.7 | 2.0E-03 |
| pentose transmembrane transporter activity | GO:0015146 | 3 | 2 | 25.7 | 66.7 | 2.0E-03 |
| amino acid transport | GO:0006865 | 4 | 2 | 19.2 | 50.0 | 3.9E-03 |
| amino acid transmembrane transporter activity | GO:0015171 | 4 | 2 | 19.2 | 50.0 | 3.9E-03 |
| cellular bud | GO:0005933 | 28 | 4 | 5.5 | 14.3 | 5.4E-03 |
| extracellular region | GO:0005576 | 15 | 3 | 7.7 | 20.0 | 6.2E-03 |
| regulation of cyclin-dependent protein serine/threonine kinase activity | GO:0000079 | 5 | 2 | 15.4 | 40.0 | 6.4E-03 |

Table S4: GO terms enriched with $p < 0.01$ and with >1 hits in the 100 top protein oscillators (Tab. S3). Simple cumulative hypergeometric distribution (Fisher's exact) tests were performed for each GO category without any multiple testing correction.

| par. | value | unit | description | source |
|---|---|---|---|---|
| | | | **Transcription/Translation Rates** | |
| $k_{nt}$ | 2000 | nt/min | mRNA elongation rate | [Mason and Struhl, 2005] |
| $\ell_{aa}$ | 10 | aa/s | peptide elongation rate | [Morisaki et al., 2016] |
| $L_{aa}$ | | aa | protein length | SGD |
| $L_{nt}$ | | nt | mRNA length: ORF+100 nt | SGD |
| | | | **Ribosomes per Cell** | |
| $B$ | 187e3±56e3 | C | ribosomes per yeast cell | [von der Haar, 2008] |
| $B_0$ | 109e3 | C | fitted intercept | [Waldron and Lacroute, 1975] |
| $\beta$ | 334e5 | C h | and slope | see Fig. S18C |
| | | | **Medians of Cytosolic RP Genes** | see Fig. S18A |
| $k$ | 264 | C $h^{-1}$ | transcription rate, duplicated genes | $2 \times k_{nt}/L_{nt}$ |
| $\ell$ | 250 | $h^{-1}$ | translation rate | $\ell_{aa}/L_{aa}$ |
| $n_B$ | 3 | | ribosome density: ribosomes per mRNA | [Arava et al., 2003] |
| $\delta_r$ | 2.79 | $h^{-1}$ | mRNA degradation rate | [Christiano et al., 2014] |
| $\delta_p$ | 0.07 | $h^{-1}$ | protein degradation rate | [Geisberg et al., 2014] |
| | | | **Strain-/Condition-specific** | |
| $\tau_{\text{hoc}}$ | 0.7 | h | time spent in HOC | |
| $\mu_f$ | 0.11 | $h^{-1}$ | IFO 0233, 0.1–0.15 | [Hansson and Häggström, 1983] |
| | | | IFO 0233, >0.1 | [Satroutdinov et al., 1992] |
| | 0.25 | $h^{-1}$ | DBY12007 | [Burnetti et al., 2016] |
| | 0.27 | $h^{-1}$ | CEN.PK122 | [van Dijken et al., 2000] |
| | 0.30 | $h^{-1}$ | LBGH 1022 | [Rieger et al., 1983] |

Table S5: **Model Parameters.** Parameter values (rounded) used for all model calculations and plots, if not specified otherwise. C is the concentration unit which we here define dimensionless as parts (molecules) per cell. Several values were intially found at `https://bionumbers.hms.harvard.edu` [Milo et al., 2010] and the original publications tracked from there (column **source**). Equations are shown for calculated parameters (without time conversion). Gene-specific transcription and translation rates were calculated from the gene and protein lengths (SGD genome annotation, release R64-1-1, where 100 nt were added to account for untranslated 5' and 3' regions) and polymerase and ribosome elongation rates, ribosome densities, mRNA and protein degradation rates were taken from the supporting information of the indicated publications. The medians of all annotated RP genes were calculated as indicated in Figure S18A, and the transcription rate was doubled to account for the duplication of RP genes in budding yeast. The critical dilution rate $\mu_f$ for strain DBY12007 was derived from an empirical oscillation model [Burnetti et al., 2016]. The value for strain CEN.PK122 was also used for the calculation of the relative growth rate for the `nocetti16` data set in Fig. 3E, where the strain was only specified as CEN.PK.

| | $\mu_f/h^{-1}$ | $\tau_{\text{hoc}}$/h | B=const/C | $\tau_{\text{hoc}}$/h | $\tau_{\max}$/h | $B_0$/C | $\beta$/(C h) |
|---|---|---|---|---|---|---|---|
| IFO 0233 | 0.121 | 0.6 | 355698 | 0.5 | 3 | 169468 | 1539090 |
| DBY12007 | 0.275 | 0.9 | 186854 | 0.6 | 6 | 101681 | 309719 |
| CEN.PK122 | 0.297 | | | 0.6 | 12 | 50840 | 415993 |
| LBGH 1022 | 0.330 | | | 0.8 | 48 | 16947 | 428340 |

Table S6: Parameters for the fitted lines in Fig. 4B and S17D. The critical dilution rates $\mu_f$ were taken 10% higher than those reported (Table S5) and $\tau_{\text{hoc}}$ was slightly shifted to manually fit data points. The left columns show parameters for the "constant ribosomes" model (Eq. S19), the right columns show parameters used for the "variable ribosomes" model (Eq. S25). The ribosome concentrations and slopes were obtained via the $\mu_f$-constraints (Eq. S23, S26), with all other parameters as provided in Table S5. Additionally, the extended model with variable ribosome concentration requires to estimate a $\tau_{\max}$ at $\mu = 0$ (Eq. S26).

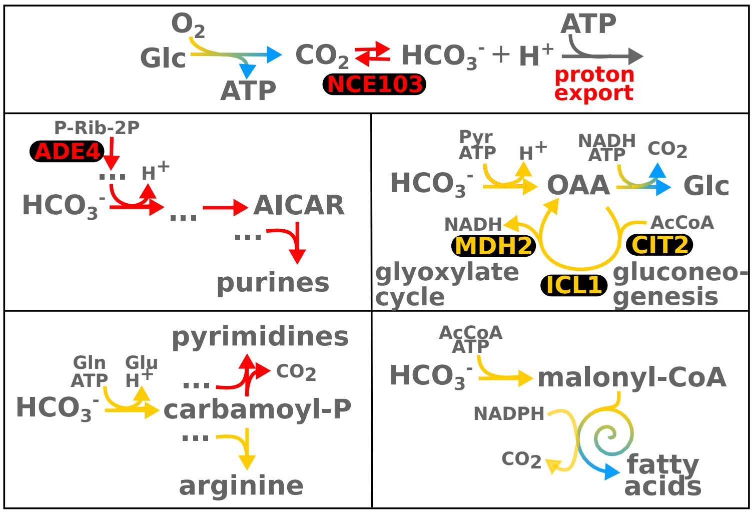


**A**

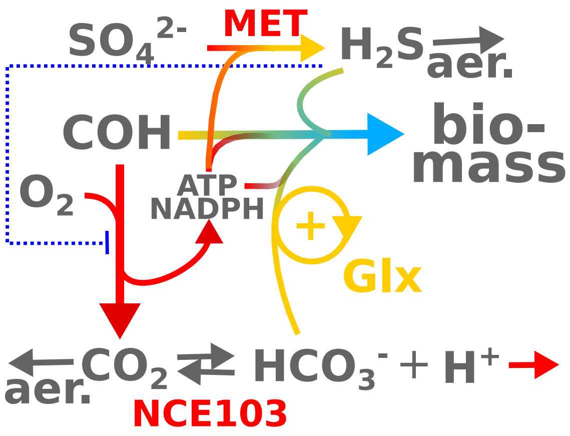


**B**

Figure S15: **Short-lived Enzymes of Biosynthetic Pathways. A:** Bicarbonate is a substrate of essential biosynthetic pathways, and the single copy enzyme NCE103 (top panel) is essential in aerated cultures, both on liquid and solid media [Aguilera et al., 2005]. Simplified pathway cartoons of the involved reactions were derived from detailed networks at the *Saccharomyces* genome database (SGD) and indicate the major substrates, products and energetic requirements (ATP, NADPH). Arrow colors indicate the expression cohorts of pathway enzymes (Fig. S9). Short-lived enzymes with high RNA amplitude (Tab. S3, Dataset S3) are indicated. Enzymes: NCE103 - carbonic anhydrase, ADE4 - phosphoribosylpyrophosphate amidotransferase (PRPPAT); MDH2 - malate dehydrogenase, cytoplasmic/peroxisomal isozyme (see text); ICL1 - isocitrate lyase, peroxisomal isozyme; CIT2 - citrate synthase, peroxisomal isozyme. Metabolites: Glc - Glucose, ATP - adenosine-triphosphate, P-Rib-2P - 5-phospho-$\alpha$-D-ribose 1-diphosphate, AICAR - 5-amino-1-(5-phospho-D-ribosyl)imidazole-4-carboxamide, Pyr - pyruvate, OAA- oxaloacetate, NADH - nicotinamide adenine dinucleotide reduced, AcCoA - acetyl coenzyme-A, malonyl-CoA - malonyl coenzyme-A, Gln - glutamine, Glu - glutamate, carbamoyl-P - carbamoyl-phosphate; all others are defined by their chemical formula. Summary and Reasoning: In HOC phase there is a deficit in $CO_2$ release (RQ $< \frac{2}{3}$, Fig. 1). At the same time proton export is at maximum, and intracellular pH increases in both short period [Satroutdinov et al., 1992, Keulers et al., 1996] and long period [O' Neill et al., 2020] systems. Carbonic anhydrase activity and pH increase could explain the "missing" $CO_2$ by a quick equilibrium shift to $HCO_3^-$, as previously suggested by Keulers et al. [1996]. During the biosynthetic pulse at the transition to LOC phase (Murray et al. [2007] and Appendix B), bicarbonate could fuel several biosynthetic pathways. Note, that $CO_2$ is lost in subsequent steps in pyrimidine and fatty acid synthesis, i.e., their is no net sequestration of $CO_2$ in these pathways. The anaplerotic reaction (middle right panel, $HCO_3^-$ and pyruvate to oxaloacetate) is not necessarily linked to the glyoxylate cycle, or gluconeogenesis, but the glyxoxylate cycle comprises three predicted oscillatory enzymes. **B: Putative feedback and feedfoward relations that could be involved in the switch from catabolic to anabolic flux at the HOC-to-LOC transition.** Carbonic anhydrase and bicarbonate-dependent pathways could interact with other short-lived enzymes in a multi-branch feedforward switch from catabolic to anabolic flux. All bicarbonate-dependent reactions also depend on ATP (A), fatty acid synthesis additionally requires NADPH. The positive feedback via the autocatalytic [Nakatsukasa et al., 2015, Barenholz et al., 2017] glyoxylate cycle (Glx, see (A)) could provide carbon backbones, the sulfate uptake pathway (MET) reduces sulfate to sulfide, requiring 6 NADPH and 4 ATP, and $H_2S$ also acts as a (negative) feedback inhibitor of respiration [Wolf et al., 2001]; specifically, beef heart mitochondrial complex IV (cytochrome C oxidase) was inhibited with a $K_i = 0.2\,\mu M$ [Nicholls, 1975, Szabo et al., 2014] compared to $[H_2S]_{max} \approx 3\,\mu M$ in our data (Fig. 1). These speculative hypotheses could be tested by dynamic extensions of flux balance analysis or resource balance analysis models, using the quantitative data we provide in Datatable S1, but final proof would require dedicated experiments and metabolic flux analysis. respiratory breakdown -¿ overflow at pyruvate node -¿ glyoxylate switch; align with transl. bursting par.

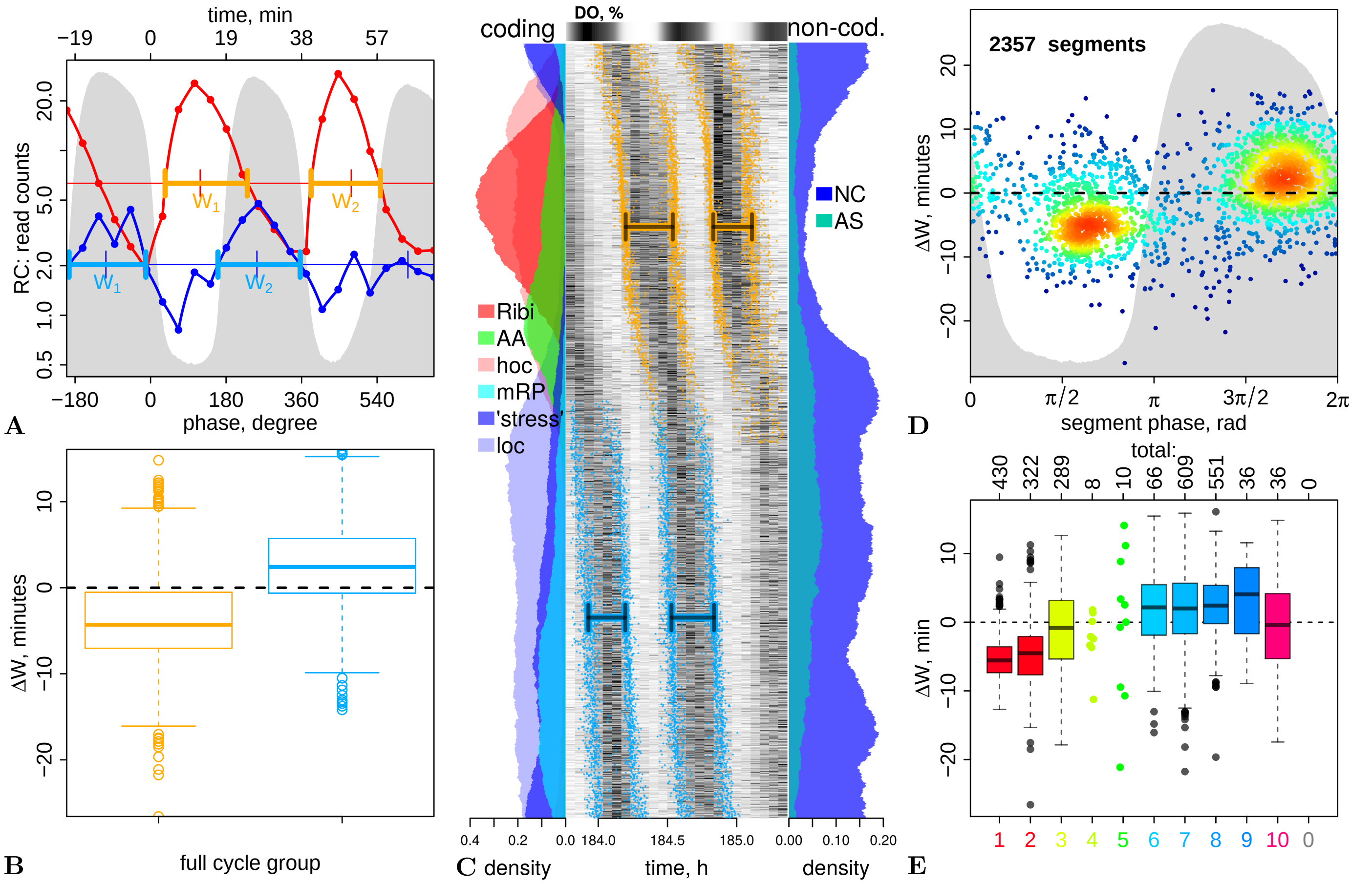


Figure S16: **Shift in Relative Peak Widths.** **A:** Transcript abundance peak width analysis shown on two examples: the segments covering the SER3 gene (red line) and its upstream noncoding RNA SRG1 (blue lines). The points are the measured samples, the lines are values interpolated to 1° resolution (0.105 min) resolution, used for peak detection and peak width calculation. The thin horizontal lines lines are the temporal medians, and the thick arrows are the peak widths $W_1$ and $W_2$. **B:** Box-plots of $\Delta W = W_2 - W_1$ separately for segments with full cycles in the first (blue) or second (orange) half of the experiment. **C:** The phase-ordered heatmap as in the main paper (Fig. 2A) but indicating the borders for peak width calculations for each segment in the same colors as in (B). **D:** 2D histogram of $\Delta W$ vs. peak phase. **E:** cluster-wise distributions of $\Delta W$. The total counts of segments for wich two peaks could be defined are shown on the top axis.

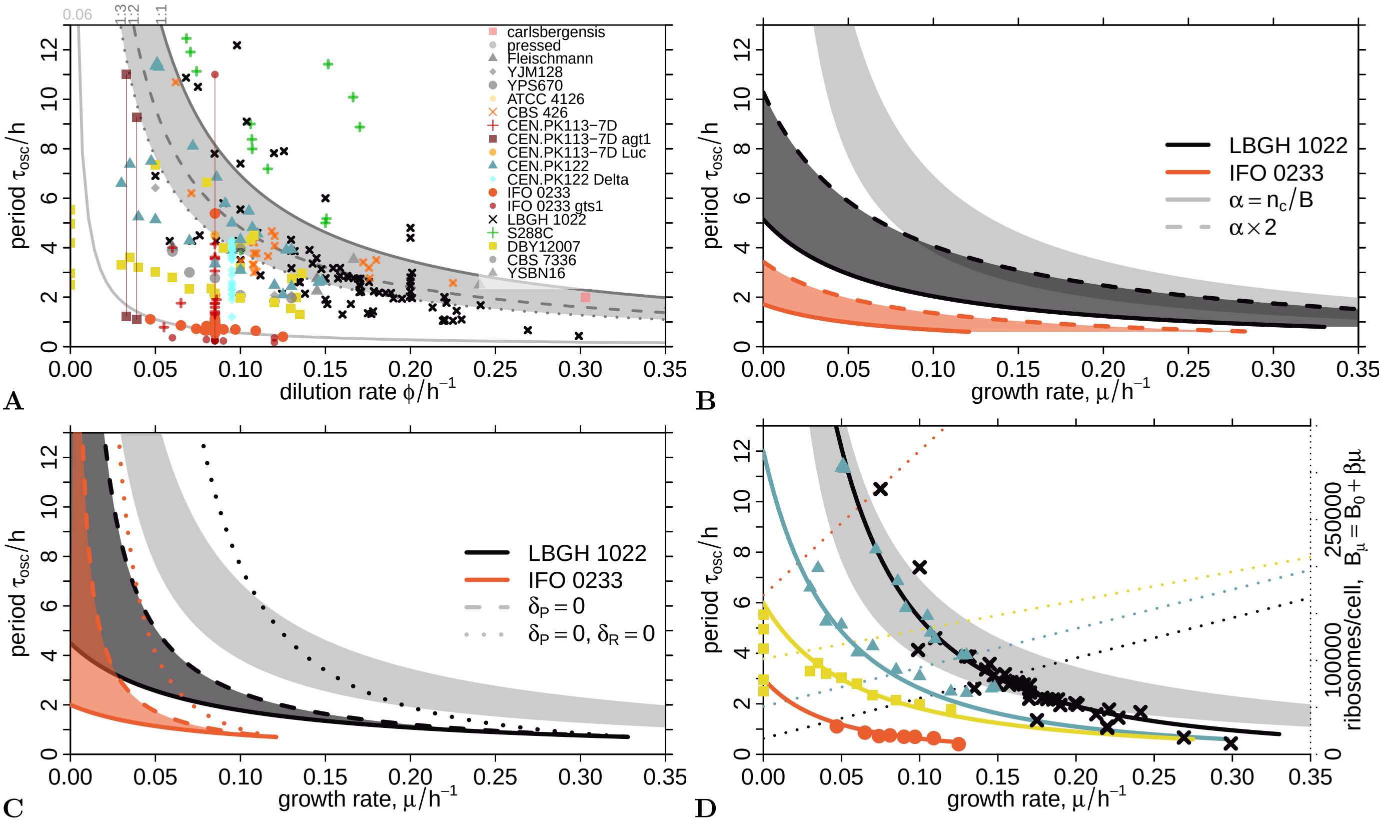


Figure S17: **Strain-Specific Parameter Constraints and Parameter Variation. A:** Periods ($\tau_{\rm osc}$) observed at specific continuous culture dilution rates ($\phi$) were digitized from plots or tables from 49 different publications and collected in a table (`periods.csv`) at `https://gitlab.com/raim/ChemostatData` [Machné, 2017]. The collection contains 308 data points from 19 budding yeast strains, incl. various mutant strains (legend), 9 data points are outside the shown plot range. Vertical lines connect experiments from the same publication. The gray background area and three black lines indicate periods expected from mode 1:1, 1:2, and 1:3 of asymmetric CDC synchronization (Eq. S31-S32, [Bellgardt, 1994]), the light gray line indicates the probabilistic mode 1:2 model with $p = 0.06$ (Eq. S34, [Duboc and von Stockar, 2000]). The gray background is shown in all plots as a reference. **B:** Effect of doubling the numerator term (here: $\alpha$) on $\tau_{\rm osc}$ (Eq. S19), from $\mu_f$-constrained $\alpha$ (Eq. S23) with all other rates as in Tab. S5, for the two indicated strains. **C:** As (B) but showing the effect of setting the protein (dashed line) degradation rate or both mRNA and protein degradation rates (dotted line) to 0. **D:** as Fig. 4B (main article) but with periods $\tau_{\rm osc}$ predicted (solid lines) using the variable ribosome model (Eq. S25), with parameters indicated in Tab. S5, where linear ribosome variation parameters (dotted lines) were calculated from $\mu_f$-constraints (Eq. S26), with manual adjustments to fit the data (Tab. S6, right columns). Period data were from experiments with variation of period under consistent conditions for four different strains (LBGH 1022 [von Meyenburg, 1969b, Heinzle et al., 1983], CEN.PK122 [Beuse et al., 1998, Burnetti et al., 2016] DBY12007 [Slavov et al., 2011, Burnetti et al., 2016] and IFO 0233 [Murray et al., 2001]); strain-specific point colors and symbols as in (A) and Fig. 4B of the main article.

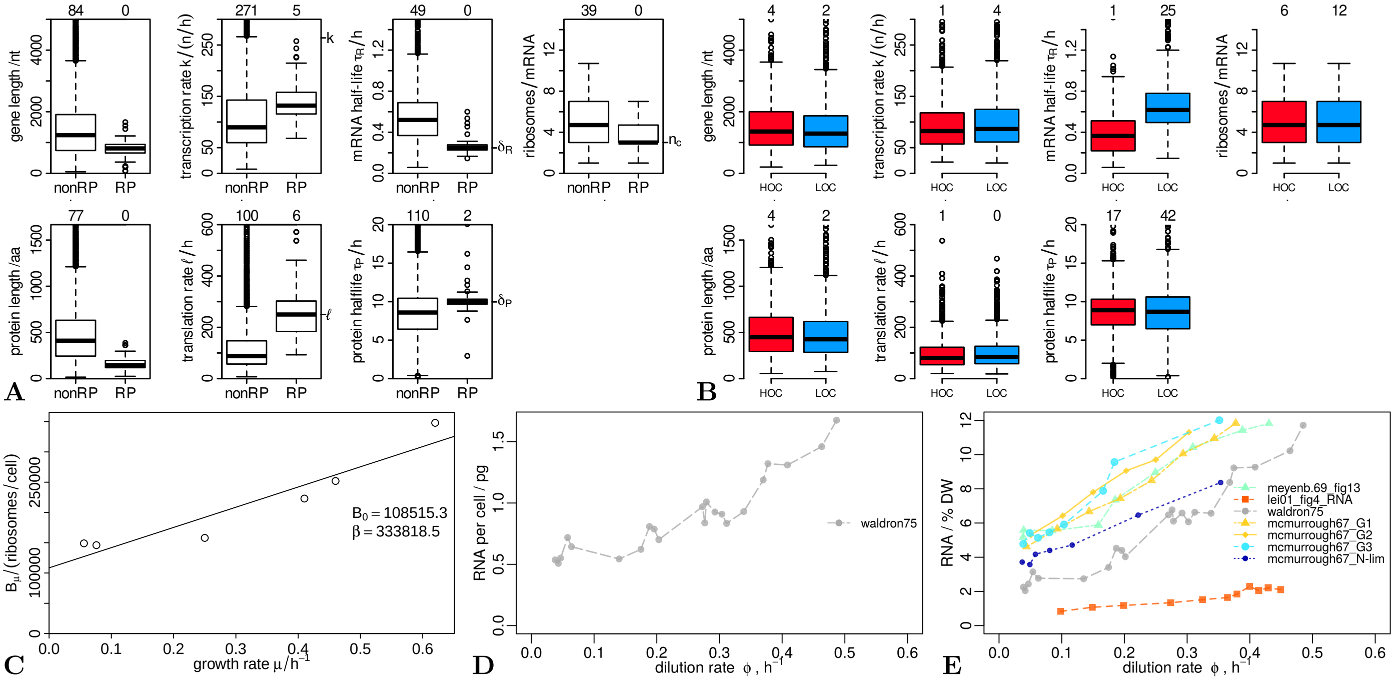


Figure S18: **Distributions of Production and Degradation Rates, and Ribosomes Per Cell. A–B:** Distribution of gene-specific parameters (production and degradation rates) used for calculation of the median RP gene values, indicated on the right y-axis and in Tab. S5. The median RP transcription rate $k$ was doubled to account for RP genes duplication. Protein half-lives indidacted as ">=100" in the original data were discarded (set to `NA`). Protein half-live data that were given for two duplicates were used for both duplicates. The boxplots are zoomed on the main distributions and the number of genes with higher values are indicated on the top x-axis. **B:** same as (A) but separately for genes classified as HOC or LOC, as used for the mRNA and protein abundance predictions. **C:** Ribosome count per cell for different growth rates as measured by Waldron and Lacroute [1975], and the linear regression used for the base values of the variable ribosome model extension (Tab. S5 and Fig. 4A of the main manuscript). **D:** RNA mass per cell from the same publication as in (C), indicating that at lower growth rates, ribosomes may be considered constant (also see (C). **E:** Additional RNA per biomass data sets [McMurrough and Rose, 1967, von Meyenburg, 1969b, Lei et al., 2001], all collected in table `growthrates.csv` at `https://gitlab.com/raim/ChemostatData` [Machné, 2017]. Note that von Meyenburg [1969b] also observed an approximately constant ribosome biomass fraction at low growth rates with strain LBGH 1022 and media very similar to the one used herein.

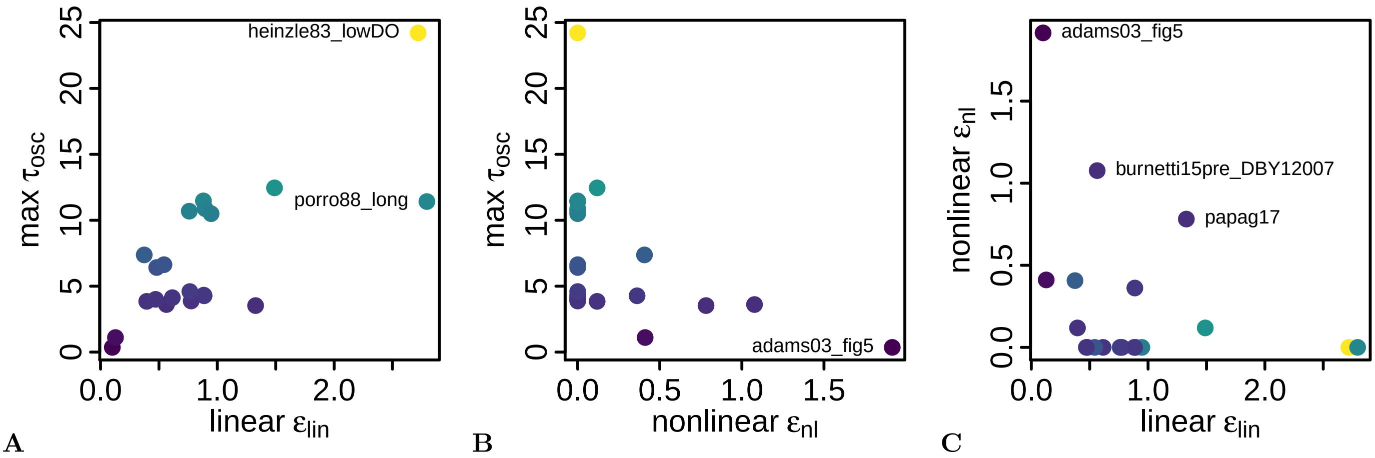


Figure S19: **Period Fit Coefficients.** The fitted nonlinear ($\epsilon_{nl}$) and linear ($\epsilon_{lin}$) helper parameters from Fig. S20, plotted against and colored by the maximal period in each data set (**A, B**) or against each other (**C**). This indicates that especially long-period data sets require higher production ($\epsilon_{lin}$) and lower degradation ($\epsilon_{nl}$) terms in Eq. S19.

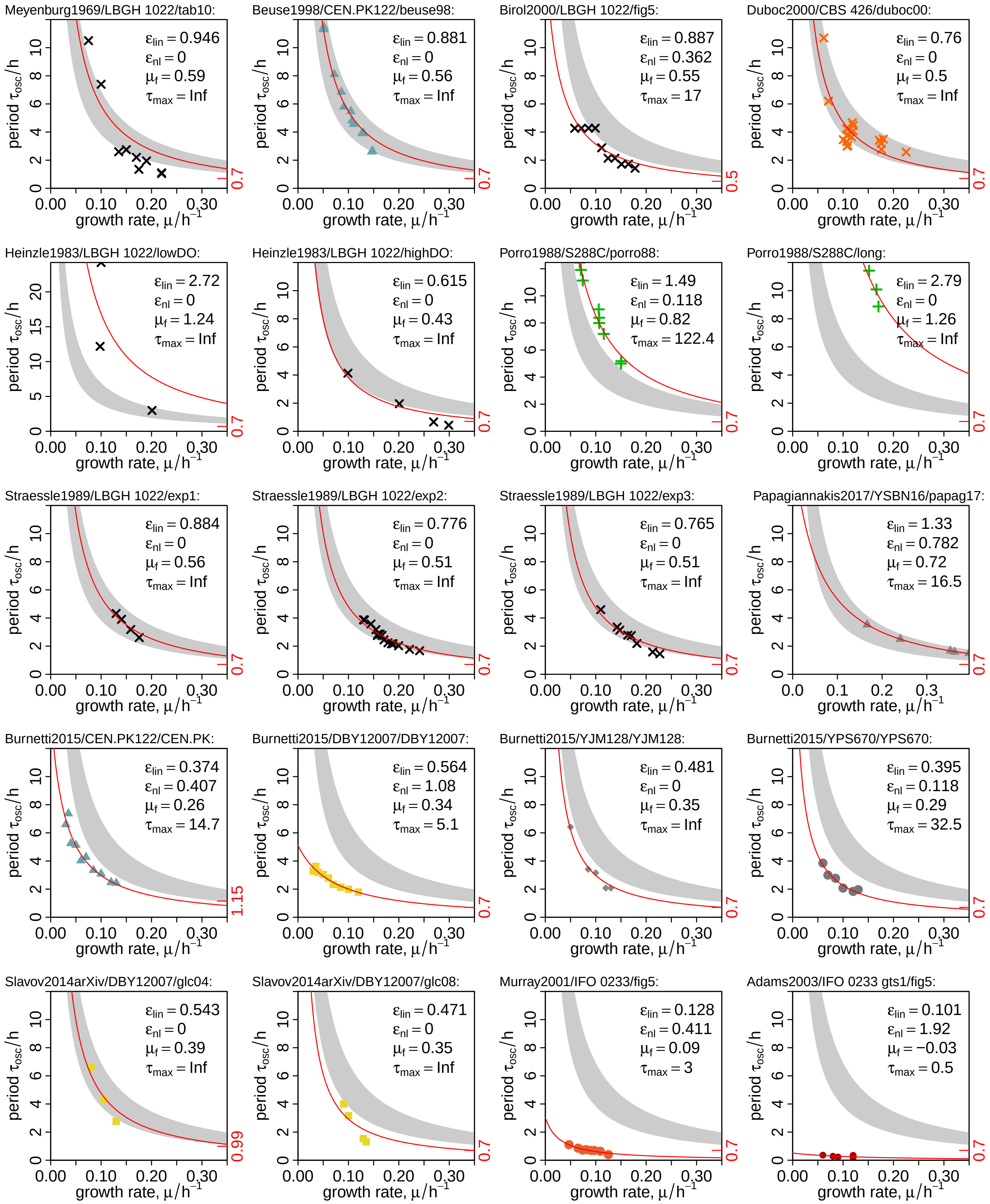


Figure S20: **Period Fits.** Equation S19 was extended by two scaling parameters, where $\epsilon_{lin}$ scales the numerator and $\epsilon_{nl}$ the protein degradation rate $\delta_p$: $\tau_{\text{osc}} = \epsilon_{lin} \frac{\tau_{\text{hoc}} k}{\mu+\delta_r} \frac{\ell}{\mu+\delta_p \epsilon_{nl}} \frac{n_B}{B}$. The parameters in Table S5 were used for non-linear least square fits (R's `nls`) of the scaling parameters to period data from literature (Fig. S17). Data sources: all data sets where one publication had more than 2 periods at different growth rates were fit [von Meyenburg, 1969b, Heinzle et al., 1983, Porro et al., 1988, Strässle et al., 1989, Duboc and von Stockar, 2000, Beuse et al., 1998, Birol et al., 2000, Murray et al., 2001, Adams et al., 2003, Slavov et al., 2014, Burnetti et al., 2016, Papagiannakis et al., 2017]. To allow for `nls` convergence with the same parameter set (Tab. S5), a lower RP mRNA degradation rate was required ($\delta_r = 1.1\,\text{h}^{-1}$), observed by Munchel et al. [2011] for cells in glucose-rich conditions. Additionally, three data sets required initialization with a higher or lower $\tau_{\text{hoc}}$, indicated on the right axis. For "Experiment 4", diamond symbols in Fig. 9 of Strässle et al. [1989], `nls` failed to converge and data is not shown.

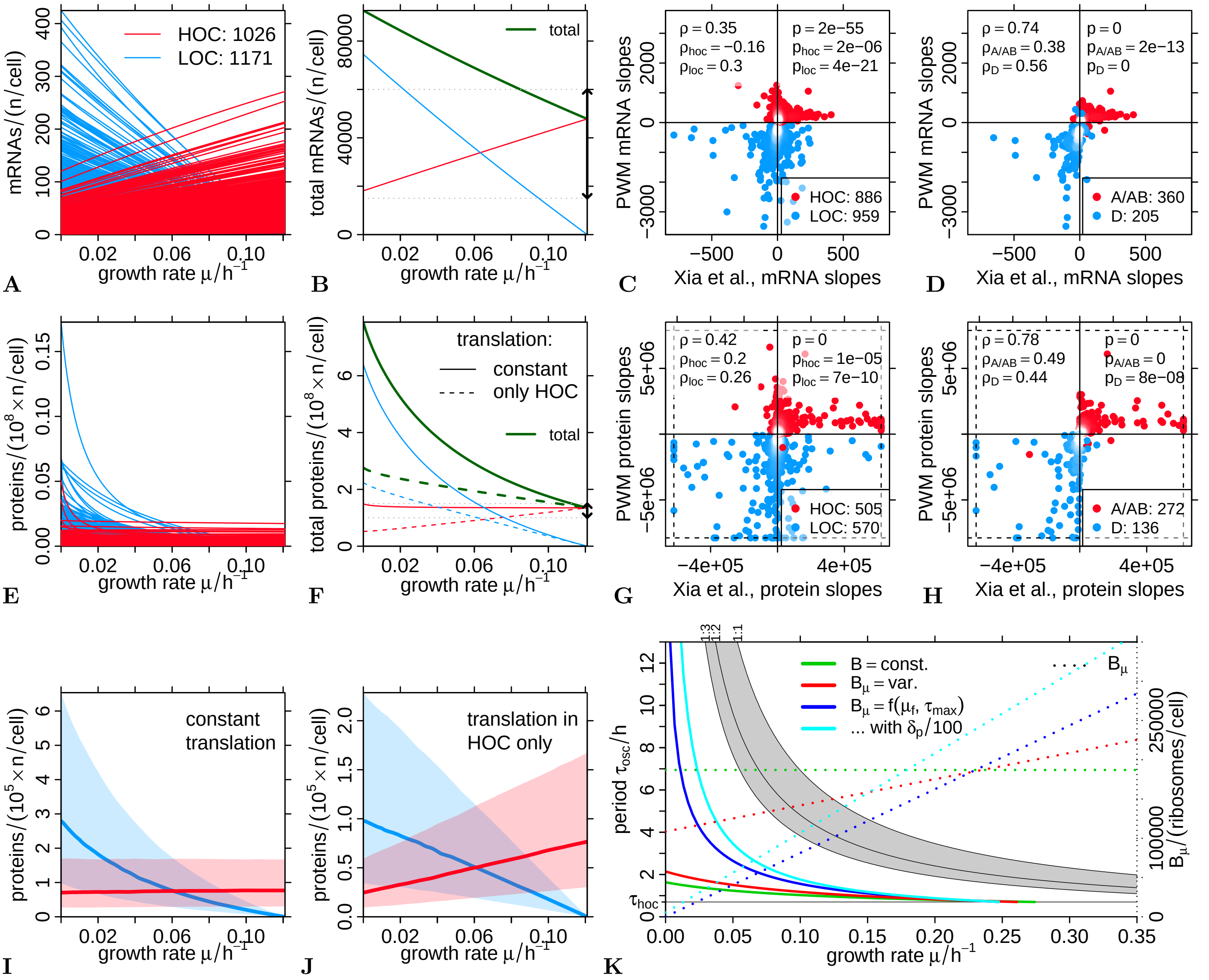


Figure S21: **mRNA and Protein Abundance Prediction.** The fractions $\phi_{\text{hoc}}$ and $\phi_{loc} = 1 - \phi_{\text{hoc}}$ were calculated for the oscillation model calibrated to IFO 0233 period data as shown in Figure 4B of the main article. Equations S15 and S16 and gene-specific rates (Fig. S18B) were then used to predict mRNA (**A**) and protein (**E**) per cell abundances of HOC- and LOC-classified genes for different growth rates. **B & F:** Total mRNA and protein per cell abundances by clusters (colored lines) and total (dark green lines). The dashed line in (F) are protein abundances assuming translation only in HOC phase (Eq. S27). The vertical arrows on the right y-axis are published estimates of total mRNA [Hereford and Rosbash, 1977, Zenklusen et al., 2008] and protein [Milo, 2013] abundances. **C - H:** The slopes of all mRNAs and proteins were calculated by linear regressions of the data shown in (A) and (E) and plotted against linear regression-derived slopes of absolute transcript (C, D) and protein (G, H) data from chemostat cultures at different growth rates by Xia et al. [2022]. The protein data were scaled to the same total amount in each sample (the mean of all experiments, $7.9 \cdot 10^7$), and only slopes where the linear regression had $r^2 \geq 0.25$ wer considered. Spearman rank correlations ($\rho$) and p-values (R function `cor.test`) for all genes and for the HOC- and LOC-specific subsets ($\rho_{\text{hoc/loc}}$) are indicated in the legends; $p = 0$ indicates a reported $p < 2.2 \cdot 10^{-16}$, i.e. below machine precision. For (C) and (G) all genes with a HOC/LOC classification for this work and available data in the comparison set were used. For (D) and (H) only the subsets of HOC- and LOC-specific clusters (A and AB, and D, respectively) from the consensus clustering by Machné and Murray [2012] were used. The number of genes in each analysis is indicated in the legend. For the plot of the protein analysis the data was cut at the dashed lines and data points on these lines have higher slopes. **I & J:** Cluster averages as in Fig. 4C of the main article (solid line: medians, transparent ranges: 25% and 75% quantiles) but for protein abundances, with the base model (I, Eq. S16) or with the additional assumption that translation only occurs in HOC (J, Eq. S27). **K:** Period predictions using the same parameters as in Fig. 4A (main article) but with the additional assumption that translation only occurs in HOC (Eq. S27), and additionally (cyan line) decreasing protein degration $\delta_p$ by 1/100.
